# Constitutively active RAS prolongs Cdc42 signalling while attenuating MAPK signalling during fission yeast mating

**DOI:** 10.1101/2025.03.06.641927

**Authors:** Emma J. Kelsall, Akatsuki Kimura, Ábel Vértesy, Kornelis R. Straatman, Mishal Tariq, Raquel Gadea, Chandni Parmar, Gabriele Schreiber, Shubhchintan Randhawa, Takashi Y. Ida, Cyril Dominguez, Edda Klipp, Kayoko Tanaka

## Abstract

The small GTPase RAS is a signalling hub activating multiple pathways, which may respond differently to a constitutively active RAS mutation. We explored this issue using fission yeast, where RAS-mediated pheromone signalling (PS) activates the MAPK^Spk1^ and Cdc42 pathways. We observed that in cells with the yeast RAS mutant, *ras1.G17V*, the MAPK^Spk1^ activation was attenuated similarly to that in wildtype cells, whereas Cdc42 activation was prolonged. We built a mathematical model implementing PS negative-feedback circuits and competition between the two Ras1 effectors, MAPKKK^Byr2^ and Cdc42-GEF^Scd1^. The model robustly predicted the MAPK^Spk1^ activation dynamics of an additional 20 PS mutants. In support of the model, a recombinant Cdc42-GEF^Scd1^ competed with MAPKKK^Byr2^ for Ras1 binding and overexpression of the Ras binding domain of either Cdc42-GEF^Scd1^ or MAPKKK^Byr2^ inhibited both downstream pathways. Our study has established that the constitutively active RAS signalling propagates differently to downstream pathways where the system prevents MAPK overactivation.

## Introduction

Proto-oncogene Ras GTPase family members are widely conserved and play pivotal roles in cell growth, differentiation and apoptosis^1^. The physiological impact of Ras mutations is highlighted in the resultant tumorigenesis and developmental disorders^2, 3^. More than 99% of identified oncogenic RAS mutations occur at codons 12, 13 and 61 of human Ras isoforms^2^ and impair efficient GTP hydrolysis and/or enhance the GDP-GTP nucleotide exchange activity^4, 5^. This results in the accumulation of GTP-bound Ras, and when overexpressed, the oncogenic Ras leads to constitutive activation of the downstream effector pathways, such as the ERK signalling pathway^6, 7^. Interestingly, however, previous studies using mouse embryonic fibroblasts (MEFs) demonstrated that oncogenic Ras expressed at its endogenous level does not cause ERK pathway hyperactivation upon a growth factor stimulation^8, 9, 10^, although the *KRAS^G12D^* MEFs showed enhanced proliferation, morphological changes and partial transformation^9^. We also observed the lack of hyperactivation of the ERK pathway in a human cell culture model that carries heterozygous oncogenic G12X mutations despite increased cell proliferation and motility^11^. Meanwhile, small GTPases, including RalA/B, Cdc42 and Rac, are required in oncogenic-RAS-driven tumorigenesis^12, 13, 14, 15, 16, 17, 18^ and the oncogenic RAS may cause small GTPase hyperactivation. However, whether the lack of ERK hyper-activation and an enhanced small GTPase activation are a conserved fundamental feature of the oncogenic RAS signalling is unclear. We wished to address the question using the model organism fission yeast, where a unique Ras homologue, Ras1, plays a crucial role in pheromone signalling by activating two downstream pathways, a pheromone MAPK cascade and Cdc42 pathway, leading to the mating of haploid cells^19, 20^(Fig. 1A, B).

**Figure 1.**
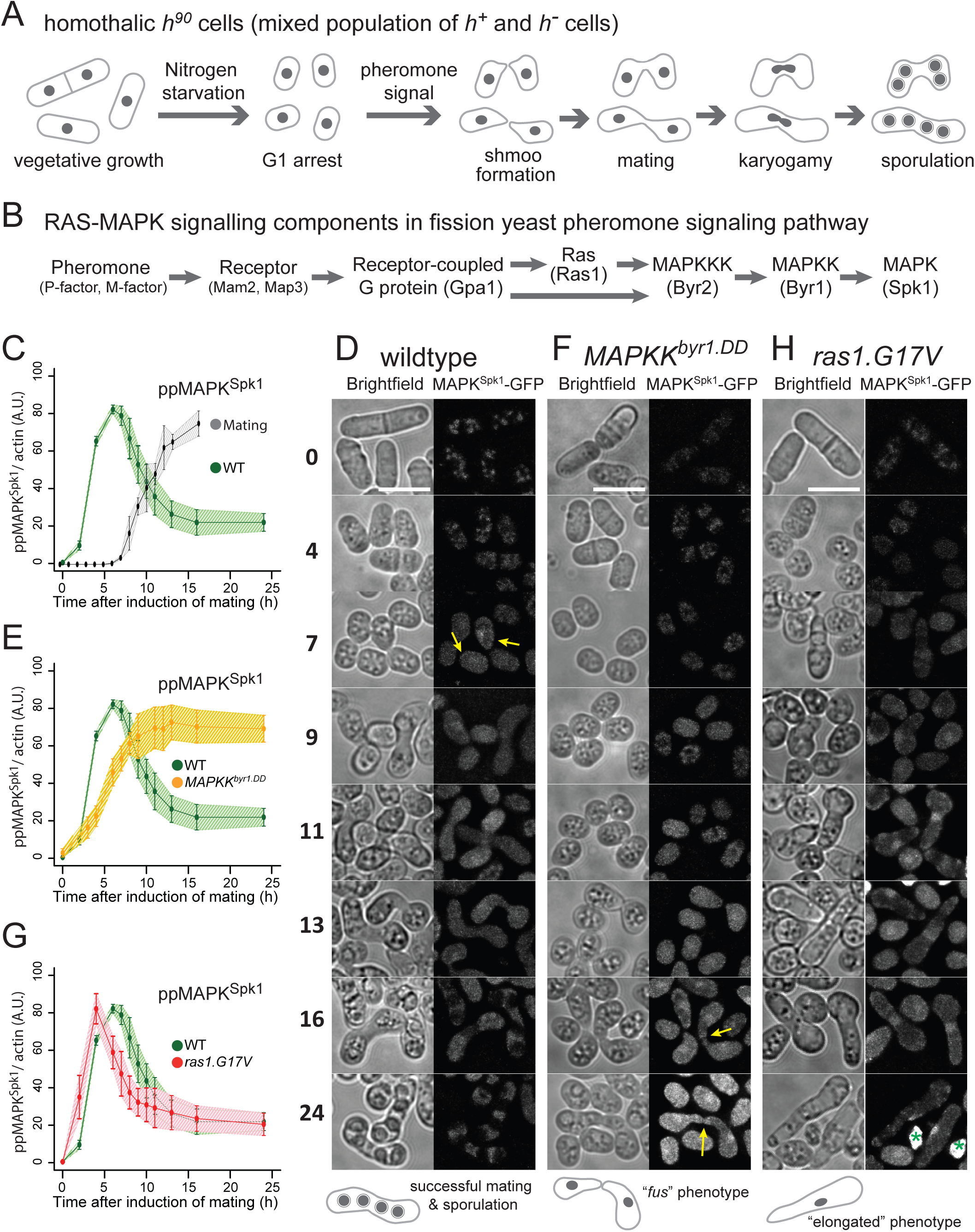
Distinct modes of MAPK^Spk1^ temporal phosphorylation profile and morphological changes during sexual differentiation in wildtype, *MAPKK^byr1.DD^* and *ras1.G17V* mutants. (A) A pictorial representation of wildtype fission yeast sexual differentiation. (B) A list of key signalling components of the fission yeast pheromone signalling pathway. The diagram reflects the prediction that Gpa1 and Ras1 separately contribute to MAPKKK^Byr2^ activation, although the precise mechanism is unknown (Xu et al., 1994). At the same time, Ras1 activation is expected to be at least partly under the influence of active Gpa1 because the *ste6* gene, encoding a Ras1 activator, is strongly induced upon successful pheromone signalling (Hughes et al., 1994). (C)-(H) Cells were induced for sexual differentiation by the plate mating assay system as described in the materials and methods. (C), (E) and (G) Quantified **pp**MAPK^Spk1^ signal from western blots of wildtype (KT3082) (C), *MAPKK^byr1.DD^* (KT3435) (E) and *ras1.G17V* (KT3084) (G) cells. Three biological replicates, shown in Supplementary Fig. S2, were used for quantitation (error bars are ±SD. Individual data is shown in Supplementary Fig. S3). Actin was used as a loading control, and quantitation was carried out using the Image Studio ver2.1 software (Licor Odyssey CLx Scanner). For the wildtype samples in (C), the % of cells mating is also indicated (n=400, three biological replicates). The wildtype **pp**MAPK^Spk1^ result (C) is also presented in (E) and (G) as a reference. (D), (F) and (H) Cellular morphology (brightfield) and localization of MAPK^Spk1^-GFP over a 24-hour time-course in wildtype (D), *MAPKK^byr^*^1^*^.DD^* (F) and *ras1.G17V* (H) cells. The time after induction of mating in hours is indicated on the left. At each time point, a brightfield image and a GFP signal image were taken and processed as described in materials and methods. Green asterisks in the time 24 h in the *ras1.G17V* cell image (H) indicate auto-fluorescence signals from inviable cell debris, which were produced through cytokinesis failure or cell lysis. Yellow arrows in panels (D) and (F) indicate transient accumulation of MAPK^Spk1^-GFP at the shmoo tips. Scale bars represent 10µm.

Upon nutritional starvation, fission yeast cells of opposite mating types (*h^+^* and *h^-^*) exchange mating pheromones^19^. Gpa1, the α-subunit of the pheromone receptor-coupled G-protein, relays the pheromone signal into the cell^21^ through Ras1, leading to the activation of the MAPK cascade consisting of Byr2 (MAPKKK), Byr1 (MAPKK) and Spk1 (MAPK)^22, 23, 24, 25, 26, 27, 28^. The *ras1.G17V* mutant, an equivalent of mammalian oncogenic *RAS.G12V* mutation, produces an excessively elongated shmoo, or a conjugation tube, upon exposure to the mating pheromone ^24^, implying that Ras1.G17V may amplify the pheromone signal^19^.

Ras1 also regulates cell morphology during vegetative growth: whilst deletion of either *gpa1*, *MAPKKK^byr^*^2^, *MAPKK^byr1^* or *MAPK^spk1^*does not result in any obvious phenotypes during vegetative cell growth^21, 29, 30^, *ras1Δ* cells lose the typical rod-shaped morphology to become rounded^24, 31^.

Studies based on recombinant protein assays and yeast-2-hybrid analysis identified Scd1, a GDP-GTP exchange factor (GEF) for Cdc42, which regulates the actin cytoskeleton and cell morphology, as well as MAPKKK^Byr2^, as Ras1 interacting proteins^32, 33, 34^. These signalling components were all found in the cell cortex before mating^20, 35, 36, 37, 38^. These observations suggest that Ras1 regulates the pheromone MAPK^Spk1^ and the Cdc42 pathways. However, how a constitutive activation of Ras1 affects the two downstream pathways has yet to be understood. By establishing conditions to induce highly synchronous mating of fission yeast cells, we were able to quantify the MAPK^Spk1^ and Cdc42 activation dynamics during the physiological mating process in wildtype and in mutants of various mating phenotypes. We build a mathematical model that serves as a prototype of a branched Ras-mediated signalling pathway, demonstrating that the *ras1.G17V* mutation does not induce constitutive activation of the MAPK^Spk1^ but rather a prolonged activation of Cdc42. The model also highlights the physiological importance of the bipartite activation of MAPKKK^Byr2^: a Ras1-dependent and a Ras1-independent mechanism, the latter of which employs the adaptor protein Ste4^39, 40^. Finally, our study reveals the crucial role played by Cdc42 in the *ras1.G17V* mutant causing the *ras1.G17V* phenotype.

## Results

### A highly synchronous mating assay allows quantifying the MAPK^Spk1^ activity during the fission yeast mating process

To monitor fission yeast pheromone signalling throughout the mating process (Fig. 1A and B), we established a protocol to induce highly synchronous mating, employing cells where the endogenous MAPK^Spk1^ is tagged with GFP-2xFLAG and conducted quantitative Western blotting of phosphorylated MAPK^Spk1^ (supplementary Fig. S1). Under this condition, homothallic *h*^90^ cells started to mate 7 hours after induction of mating (Fig. 1C, grey line). The phosphorylated (active) MAPK^Spk1^ (**pp**MAPK^Spk1^) signal was first detected three hours after induction of mating and reached its peak at about five to seven hours when cell fusion was also initially observed (Fig. 1C, green line. Original membrane images and quantitations presented in Supplementary Fig. S2 and S3). The **pp**MAPK^Spk1^ then gradually decreased to a non-zero level as meiosis continued towards sporulation. It was also noted that the total MAPK^Spk1^-GFP was essentially not expressed during the vegetative cycle, but it was promptly induced by nitrogen starvation (Supplementary Fig. S2 and S3). The *mapk^spk1^* gene is a known target of the transcription factor Ste11^41^, which itself is activated by MAPK^Spk1^ ^42^. This positive feedback loop likely facilitates a swift increase of MAPK^Spk1^ expression upon nitrogen starvation.

We found that MAPK^Spk1^-GFP localised to both the cytosol and the nucleus, with some nuclear accumulation, before gradually disappearing as the mating process came to an end (Fig. 1D and Supplementary Fig. S4). Transient foci of GFP signals were also found at the cell cortex, as has been reported for MAPKK^Byr1^, the activator of MAPK^Spk1^ ^36^ (Fig. 1D, yellow arrows and Supplementary Fig. S4).

### Constitutively active MAPKK^Byr1.DD^ mutant causes constitutive activation of MAPK^Spk1^

Activation of MAPKK family kinases is mediated by dual phosphorylation of conserved Ser/Thr residues^43^, and when the corresponding Ser214/Thr218 were substituted to aspartic acid, the resultant fission yeast MAPKK^Byr1.DD^ was expected to act as a constitutively active MAPKK^44^. Indeed, the level of **pp**MAPK^Spk1^ in the MAPKK^Byr1.DD^ mutant remained high after reaching its highest intensity at around 13 hours after induction (Fig. 1E, yellow line, Supplementary Fig. S2 and S3), although the initial increase of the **pp**MAPK^Spk1^ signal was slower compared to the wildtype strain. The **pp**MAPK^Spk1^ level remained high even 48 hours after the induction of mating (Supplementary Fig. S5). The result highlights that the endogenously expressed MAPKK^Byr1.DD^ is proficient in retaining the high **pp**MAPK^Spk1^ level, and the counteracting de-phosphorylation by phosphatases Pmp1 and Pyp1^45^ is not efficient in downregulating the **pp**MAPK^Spk1^ signal in the presence of MAPKK^Byr1.DD^. Consistent with the observed slower increase in **pp**MAPK^Spk1^, the nuclear localisation of MAPK^Spk1^ was also delayed compared to the wildtype cells (Fig. 1F). A strong nuclear MAPK^Spk1^-GFP signal was then observed in the paired “*fus”* (fusion deficient) cells, an intriguing phenotype of the MAPKK^Byr1.DD^ cells (Fig.1F)^36, 44^. Interestingly, the projection tips of the paring cells often show increased MAPK^Spk1^-GFP signal (Fig. 1F, yellow arrows and Supplementary Fig. S6).

### The *ras1.G17V* mutation causes immediate but transient MAPK^Spk1^ activation

The fission yeast equivalent of human oncogenic *RAS.G12V* is *ras1.G17V,* which induces an excessively elongated shmoo, and the cells fail to recognize a partner and become sterile^22, 24^. The “elongated shmoo” phenotype was interpreted as an excess activation of the Ras1 downstream pathway(s), leading to a prediction that the *ras1.G17V* causes over-activation of MAPK^Spk1^ ^19, 20^. However, the **pp**MAPK^Spk1^ signal intensity declined in a comparable manner to wildtype cells, indicating that down-regulation of **pp**MAPK^Spk1^ is effective in the *ras1.G17V* mutant, unlike in the *MAPKK^byr1.DD^*mutant (Fig. 1G, red line, Supplementary Fig. S2 and S3). Correspondingly, the nuclear MAPK^Spk1^-GFP signal also declined 16-24 hours after induction of mating (Fig. 1H). Interestingly, an immediate increase of the **pp**MAPK^Spk1^ signal upon induction of mating was observed (Fig. 1G, red line).

### The elongated *ras1.G17V* shmoos develop with a detectable level of MAPK^Spk1^: neither amplitude nor duration of ppMAPK^Spk1^ signal influences the *ras1.G17V* phenotype

Having observed that *MAPKK^byr1.DD^* and *ras1.G17V* show different MAPK^Spk1^ activation profiles (sustained vs transient) and different morphological phenotypes (“*fus*” vs elongated), we examined a correlation between the MAPK^Spk1^ activation profiles and the cell morphology. The *ras1.G17V MAPKK^byr1.DD^* double mutant showed constitutive activation of MAPK^Spk1^ (Fig. 2A, light blue line, Supplementary Fig. S2 and S3). The nuclear **pp**MAPK^Spk1^-GFP signal in the *ras1.G17V MAPKK^byr1.DD^* double mutant was present 24 hours after induction of mating, unlike the *ras1.G17V* single mutant cells (Fig. 2B), confirming that the *ras1.G17V MAPKK^byr1.DD^* double mutant cells retained a high **pp**MAPK^Spk1^ level. Thus, *MAPKK^byr1.DD^* is overall epistatic to *ras1.G17V* in terms of the MAPK^Spk1^ activation status.

**Figure 2.**
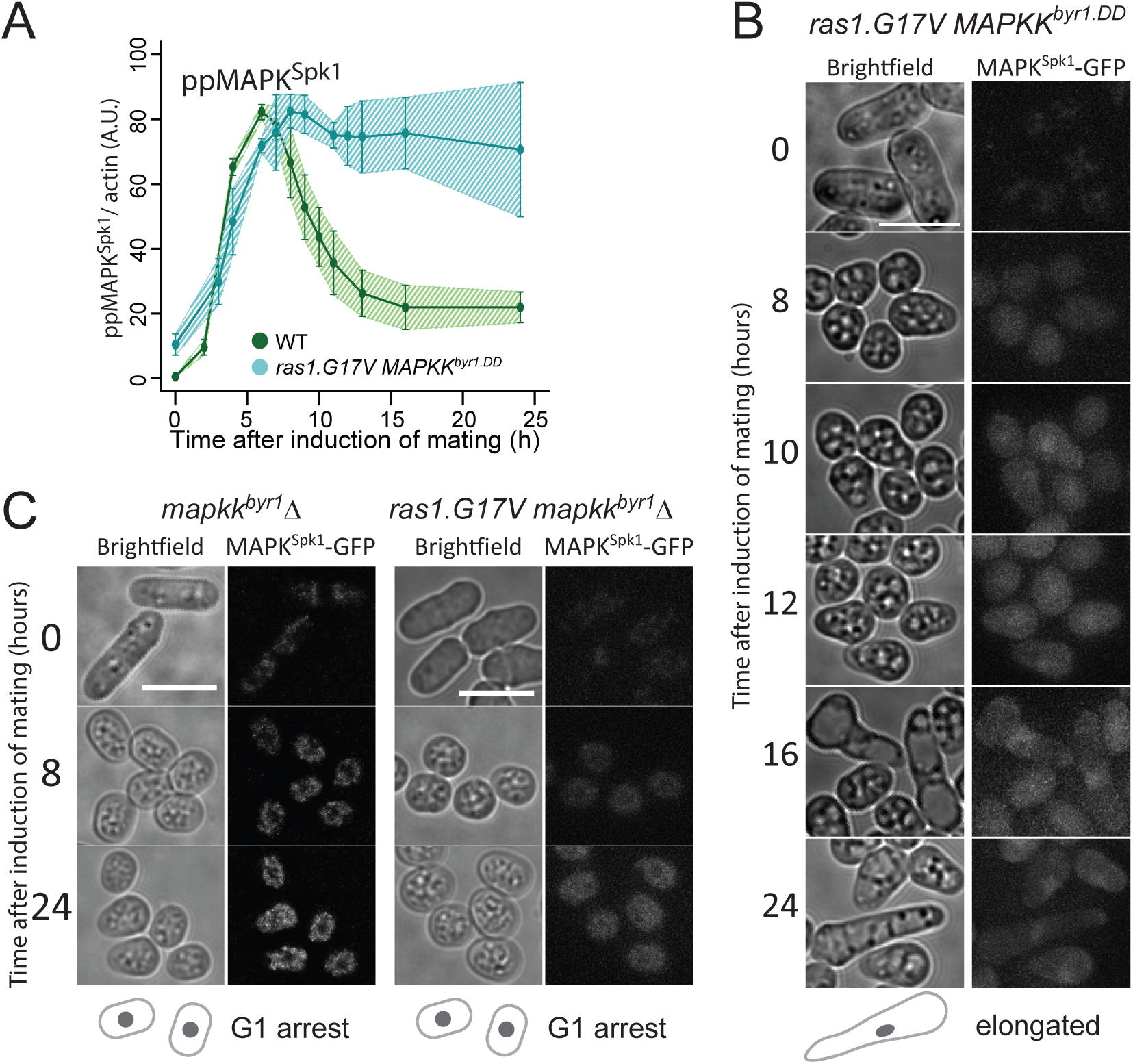
In the *ras1.G17V MAPKK^byr^*^1^*^.DD^* double mutant, the MAPK^Spk^^1^ phosphorylation profile follows *MAPKK^byr^*^1^*^.DD^* single mutant phenotype whilst cell morphology mimics the *ras1.G17V* single mutant phenotype. (A) MAPK^Spk1^ phosphorylation status in the *ras1.G17V MAPKK^byr^*^1^*^.DD^* double mutant cells (KT3439). Cells were induced for mating by the plate mating assay system as described in the materials and methods. Quantitated **pp**MAPK^Spk1^ signal (arbitrary unit) from western blots is presented. Original membrane images are presented in Supplementary Fig. S2. Three biological replicates were used for quantitation (error bars are ±SD). α-tubulin was used as a loading control, and quantitation was carried out using the Image Studio ver2.1 software (Licor Odyssey CLx Scanner). The wildtype **pp**MAPK^Spk1^ result (Fig. 1C) is also presented as a reference. (B) The terminal mating phenotype of *ras1.G17V MAPKK^byr^*^1^*^.DD^* double mutant is a phenocopy of *ras1.G17V* single mutant which shows the “elongated” morphology. Images were taken of *ras1.G17V MAPKK^byr^*^1^*^.DD^* double mutant (KT3439) in the same way as in Fig. 1. Time after induction of mating in hours is indicated on the left. (C) There is no morphological change in the absence of MAPK^Spk1^ signalling. Cell images of *MAPKK^byr1^Δ* (KT4300) and *ras1.G17V MAPKK^byr1^Δ* (KT5215) strains are shown. Images were taken in the same way as in Fig. 1. Time after induction of mating in hours is indicated on the left of each series. The scale bars represent 10µm.

However, for cell morphology, the *ras1.G17V MAPKK^byr1.DD^* double mutant cells showed the “elongated” *ras1.G17V* phenotype (Fig. 2B), demonstrating that *ras1.G17V* was epistatic to *MAPKK^byr1.DD^* regarding cell morphology.

Interestingly, the morphological change in the *ras1.G17V MAPKK^byr1.DD^*double mutant was first noticed 16 hours after induction of mating, much later than the *ras1.G17V* single mutant, but as a comparable timing as the “*fus*” formation of the *MAPKK^byr1.DD^* single mutant (Fig. 2B). Given the slower increase of the **pp**MAPK^Spk1^ signal in the *MAPKK^byr1.DD^* mutant, we predicted that the “elongated” shmoo formation still requires a certain level of MAPK^Spk1^ activity. Indeed, when *MAPKK^byr1^* was deleted in the *ras1.G17V* mutant, not only was the MAPK^Spk1^ activation abolished (Supplementary Fig. S1G), but also shmoo formation was abolished (Fig. 2C). Based on these observations, we concluded that the Ras1 status, but not the MAPK^Spk1^ activation profile, determines the cell morphology. Yet, the *ras1.G17V* “elongated shmoo” phenotype still requires a certain level of MAPK^Spk1^ activity, which defines the timing of the shmoo formation.

### Cdc42 is required for the shmoo formation and full activation of the MAPK^Spk1^

During the vegetative cycle, *ras1Δ* cells show spherical cell morphology^22, 24^ where polarised localisation of active Cdc42 is compromised^46^(Supplementary Fig. S7) which can be detected by CRIB-GFP, a specific binder of the active GTP-bound form of Cdc42^47^. The current understanding is that Ras1 activates Cdc42, which leads to the activation of downstream Ste20-like kinase, Pak1/Shk1^48, 49, 50, 51^, resulting in actin reorganisation and shmoo formation under mating conditions^35, 52^.

The elongated shmoo phenotype was lost when the *scd1*, encoding a Cdc42-GEF, was deleted in the *ras1.G17V* mutant. Instead, the cells showed a mating-deficient phenotype similar to the *cdc42-GEF^scd1^Δ* single mutant, supporting the model that Cdc42 acts downstream of Ras1 to cause morphological changes (Fig 3A).

**Figure 3.**
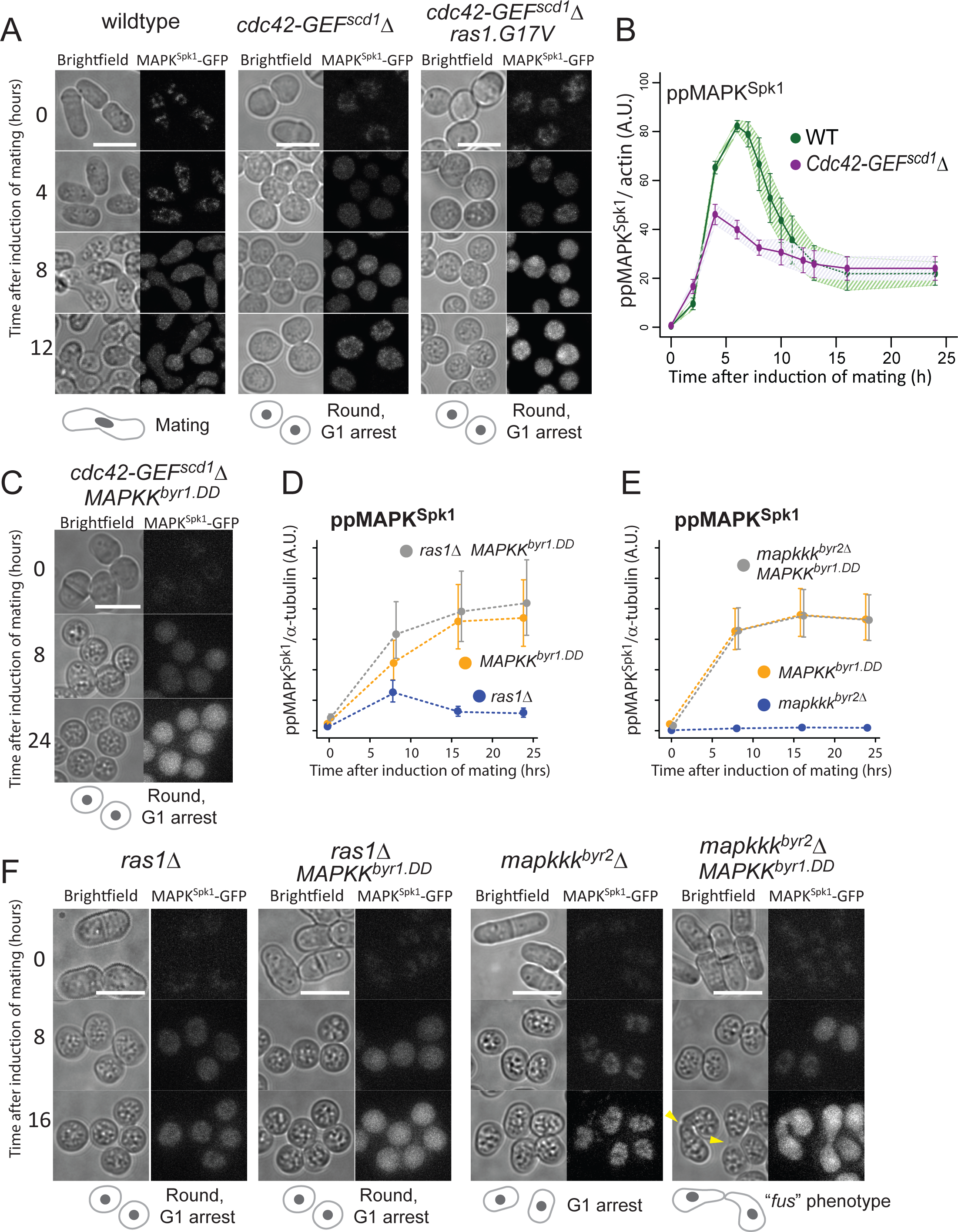
Ras1 activates both MAPK^Spk1^ and Cdc42 pathways during pheromone signaling. (A) The deletion of *scd1* causes round cell morphology and sterility. Images of WT (KT3082), *scd1Δ* (KT4061) and *scd1Δ ras1.G17V* double mutant (KT4056), expressing the MAPK^Spk1^-GFP, were taken in the same way as in Fig.1. Numbers on the left represents hours after induction of mating. (B) The **pp**MAPK^Spk1^ levels during the sexual differentiation in *scd1Δ* (KT4061) cells. Results of three biological replicates (error bars are ±SD) are presented. Original membrane images are presented in Supplementary Fig. S2. The wildtype **pp**MAPK^Spk1^ result presented in Fig.1C is also shown in green as a reference. (C) Cell images of *scd1Δ MAPKK^byr^*^1^*^.DD^* double mutant (KT4047). The images were taken in the same way as in Fig.1. Numbers on the left represent hours after induction of mating. (D) The **pp**MAPK^Spk1^ levels in *ras1Δ* (KT4323), *MAPKK^byr1.DD^*(KT3435) and *ras1Δ MAPKK^byr1.DD^* (KT4359) cell extracts. Original Western blotting data is presented in Supplementary Fig. S8A. (E) The **pp**MAPK^Spk1^ levels in *mapkkk^byr2^Δ* (KT3763), *MAPKK^byr1.DD^* (KT3435) and *mapkkk^byr2^Δ MAPKK^byr1.DD^* (KT4010) cell extracts. Original Western blotting data is presented in Supplementary Fig. S8B. For (D) and (E), quantification was carried out using the Image Studio ver2.1 (Li-cor). (F) Cell images of the strains mentioned in (D) and (E) were taken in the same way as in Fig.1. Numbers on the left represent hours after induction of mating. For all the images presented in (A), (C) and (F), the scale bars represent 10µm.

Interestingly, the **pp**MAPK^Spk1^ level was substantially reduced in the *cdc42-GEF^scd1^Δ* mutant compared to the wildtype (Fig. 3B, purple line, Supplementary Fig. S2 and S3). The result agreed with a previous study that predicted active Cdc42 to contribute to the activation of MAPKKK^Byr233^. Even so, some MAPK^Spk1^ activation occurred in the *cdc42-GEF^scd1^Δ* mutant, and a nuclear MAPK^Spk1^-GFP signal was detectable (Fig. 3A).

A reduced but substantial level of MAPK^Spk1^ activation in the *cdc42-GEF^scd1^Δ* mutant means that the mating deficiency of this mutant is unlikely to result from the lack of MAPK^Spk1^ activation. Indeed, the introduction of *MAPKK^byr1.DD^* to the *cdc42-GEF^scd1^Δ* mutant did not restore the mating deficient phenotype, even though nuclear MAPK^Spk1^-GFP highly accumulated (Fig. 3C). The result shows that active Cdc42 function is required for the mating process regardless of the MAPK^Spk1^ activation status. Taken together with the essential role of MAPK^Spk1^, we concluded that the mating pheromone signalling feeds into two pathways, MAPK^Spk1^ and Cdc42.

### Ras1 activates two effector pathways, MAPK^Spk1^ and Cdc42

In order to further clarify the role of Ras1, we examined the MAPK^Spk1^ activation status and cell morphology in the following four strains: *ras1Δ* mutant, *ras1Δ MAPKK^byr1.DD^* double mutant, *MAPKKK*^byr2^*Δ* mutant and *MAPKKK*^byr2^*Δ MAPKK^byr1.DD^* double mutant. As mentioned earlier, the vegetatively growing *ras1Δ* cells show a round morphology with reduced cortical signal of CRIB-GFP, demonstrating that Cdc42 activation is compromised (Supplementary Fig. S7). The deletion of *ras1* also causes substantial reduction, but not complete elimination, of the **pp**MAPK^Spk1^ (Fig. 3D, blue line, and Supplementary Fig. S8A); thus, Ras1 plays an important role in activating both Cdc42 and MAPK^Spk1^ pathways. Introduction of the *MAPKK^byr1.DD^* mutation into the *ras1Δ* mutant cells induces the constitutive **pp**MAPK^Spk1^ (Fig. 3D, grey line and Supplementary Fig. S8A) but does not affect the round cell morphology, and cells remain sterile (Fig. 3F, the 2^nd^ left panel). In striking contrast, the sterile phenotype of the *MAPKKK^byr2^Δ*, associated with a complete lack of shmoo formation (Fig. 3F, the 2^nd^ right panel), was converted to the “*fus*” phenotype when combined with the *MAPKK^byr1.DD^* mutation (Fig. 3F, the far right panel). As expected, the *MAPKKK^byr2^Δ MAPKK^byr1.DD^*double mutant shows MAPK^Spk1^ constitutive activation (Fig. 3E and Supplementary Fig. S8B). Thus, unlike the cases of *scd1Δ* or *ras1*Δ, the lack of *MAPKKK^byr2^* can be bypassed by constitutive activation of MAPK^Spk1^, indicating that the sole role of MAPKKK^Byr2^ is to activate the MAPK^Spk1^, unlike its upstream activator, Ras1, which also activates Cdc42 pathway.

### Ras1.G17V causes accumulation of Cdc42-GTP at the cell cortex

Having observed a relatively mild influence of Ras1.G17V towards the MAPK^Spk1^ activation profile, we next examined whether the Cdc42 pathway was affected by the *ras1.G17V* mutation. As previously observed, dynamic foci of CRIB-GFP appeared on the cell cortex upon induction of mating^52^(Fig. 4A and B). In our experimental condition, more than 80% of the wildtype cells showed the cortical CRIB-GFP signal at 4.5 hours after induction of mating (Fig. 4B). The cortical CRIB-GFP foci became concentrated at the mating site and quickly disappeared once cells fused successfully to form zygotes (Fig. 4A and B). In striking contrast, in the *ras1.G17V* mutant cells, the cortical CRIB-GFP signal persisted, often at the elongated tip end of the cells, even 12.5 hours after induction of mating (Fig. 4A and B). The signal could still be seen in about 40% of the cells 22.5 hours after induction of mating (Fig. 4A and B). The result shows that the Cdc42 pathway is excessively activated in the *ras1.G17V* mutant, and the tip localisation of Cdc42^GTP^ indicates that the signature “elongated” *ras1.G17V* morphological phenotype is caused by hyperactivation of the Cdc42 pathway.

**Figure 4.**
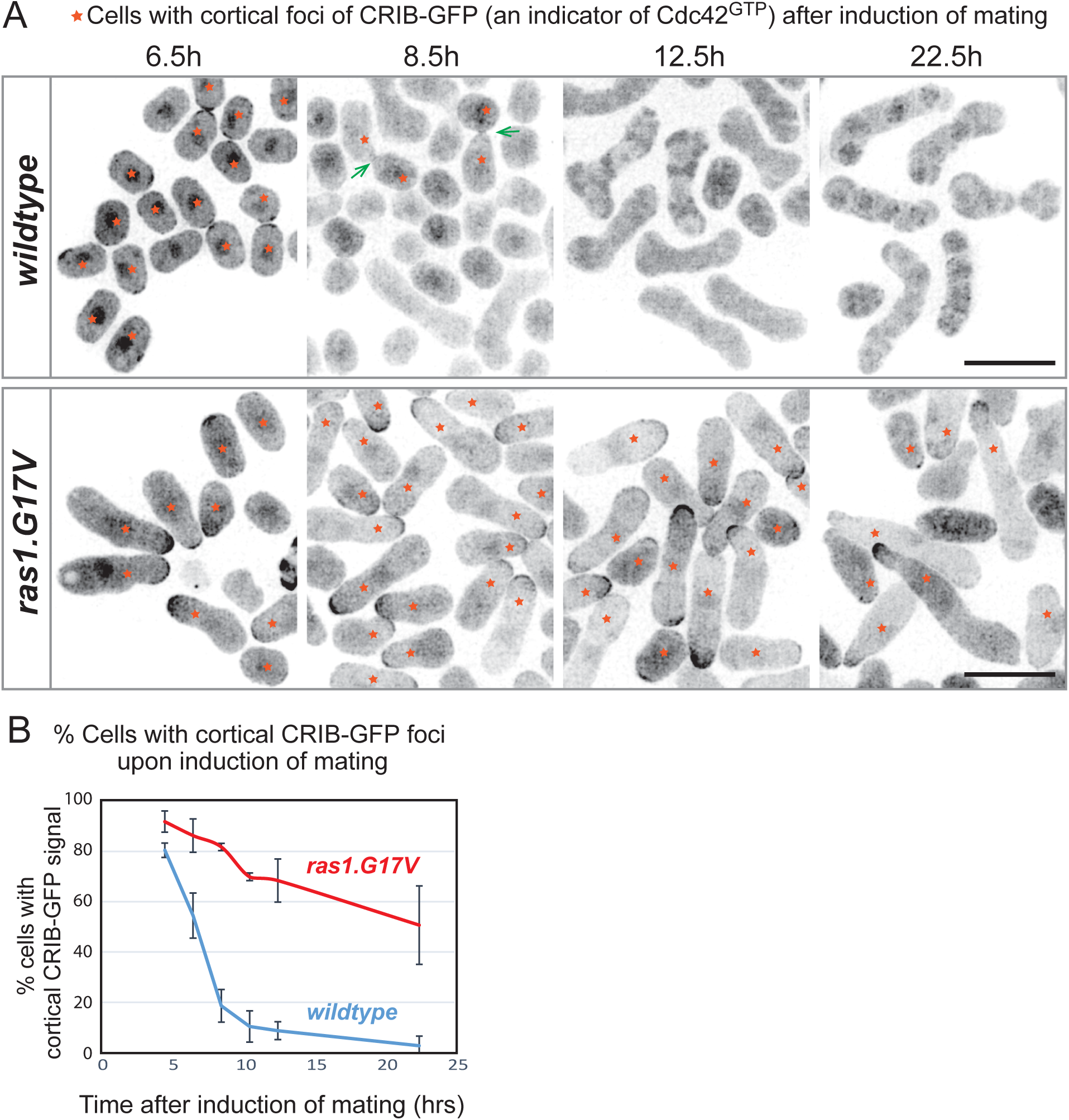
Ras1.G17V induces cortical Cdc42^GTP^ accumulation. (A) Cell morphology and localization of Cdc42^GTP^, indicated by CRIB-GFP signal, during the sexual differentiation process. Wildtype (KT5077) and *ras1.G17V* (KT5082) mutant cells were induced for sexual differentiation by the plate mating assay condition (Materials and Methods), and live cell images were taken at the indicated time after the induction. Representative CRIB-GFP signal images are presented. Orange stars indicate cells with cortical CRIB-GFP foci. Rapidly disappearing CRIB-GFP signals at the fusion site of wildtype mating cells are indicated by green arrows at the time 8.5h image. The scale bar represents 10 µm. (B) Quantification of the results presented in (A). At each time point (4.5h, 6.5h, 10.5h, 12.5h and 22.5h after induction of sexual differentiation), 150 cells were examined to determine whether they have cortical CRIB-GFP foci. % cells with cortical CRIB-GFP foci are presented. The experiment was repeated three times, and the mean values and SDs were plotted in the graph.

The effect of *ras1.G17V* mutation towards Cdc42 activation was also observed during vegetative growth, producing the strongest cortical CRIB-GFP signal at the cell tip in the *ras1.G17V* mutant (Supplementary Fig. S7). The effect of *ras1.G17V* on cell morphology was best recognised when the *rgl4* that encodes a GTPase activation protein negatively regulating Cdc42^46, 47, 53^ was deleted, as the resultant *rga4Δ ras1.G17V* double mutant cells were larger and round with an increased CRIB-GFP signal (Supplementary Fig. S7). These results fit the hypothesis that Ras1.G17V has an enhanced capability to activate Cdc42.

### Two Ras effectors, MAPKKK^Byr2^ and Cdc42-GEF^Scd1^, compete with each other for Ras1

We next examined whether the two Ras effectors, MAPKKK^Byr2^ and Cdc42-GEF^Scd1^, compete with each other for Ras1. First, we overexpressed the Ras binding domains (RBDs) of MAPKKK^Byr2^ and Cdc42-GEF^Scd1^ in the *ras1.G17V* cells and examined the activation status of MAPK^Spk1^ and Cdc42. For MAPKKK^Byr2^, the region spanning the residues 65-180 (Byr2-RBD) was used following previous structural studies^54^, and for Cdc42-GEF^Scd1^, the region spanning the residues 760-872 (Scd1-PB1) was used based on its amino acid sequence similarity with the Ras Associating (RA) domain of mammalian RalGDS.

Overexpression of Byr2-RBD in the *ras1.G17V* cells substantially reduced the **pp**MAPK^Spk1^ level, whereas the total MAPK^Spk1^ level was only marginally affected (Fig. 5A and Supplementary Fig. S9), indicating that the Byr2-RBD competed against the endogenous full-length MAPKKK^Byr2^ in activating the MAPK^Spk1^. Strikingly, overexpression of Scd1-PB1 also inhibited the MAPK^Spk1^ activation, exhibiting its capability to interfere with the MAPK^Spk1^ pathway (Fig. 5A and Supplementary Fig. S9).

**Figure 5.**
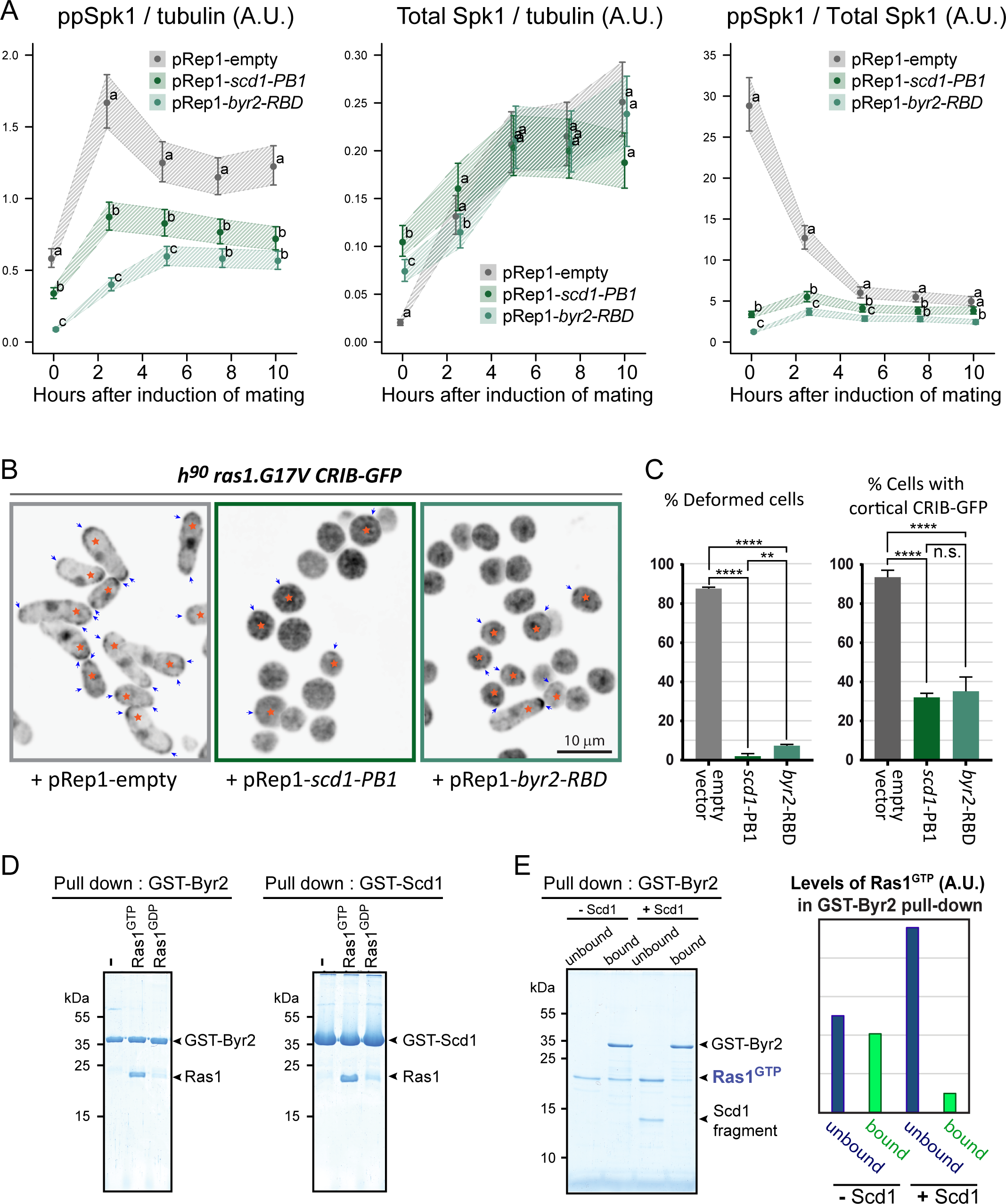
Two Ras effectors, MAPKKK^Byr2^ and Cdc42-GEF^Scd1^, compete each other for Ras1. (A) Cells harboring *ras1.G17V* and MAPK^Spk1^-GFP-2xFLAG (KT5940) were transformed with either pRep1 empty vector, pRep1-*scd1*(760-872)-2xFLAG or pRep1-*byr2*(65-180)-2xFLAG and cultured in MM+N without thiamine for 24 hours. Cells were induced for sexual differentiation by the plate mating assay system, and the levels of ppMAPK^Spk1^-GFP, total MAPK^Spk1^ and α-tubulin (loading control) were examined by Western blotting. Three biological replicates, shown in Supplementary Fig. S9, were quantified using the Image Studio ver2.1 software (Licor Odyssey CLx Scanner). The data was analyzed by generalized linear mixed models and their least-squares means (± 1SE) are shown. Different letters (a, b and c) indicate significant differences (α = 0.05) among the three samples, pRep1-empty, pRep1-scd1-PB1 and pRep1-byr2-RBD, within each hour after induction of mating. When all three samples were deemed distinct, they were denoted differently: a, b and c. When all the samples were deemed indistinguishable, they were denoted with the same category: a. (B) Cell morphology and localization of Cdc42^GTP^, indicated by the CRIB-GFP signal, during the sexual differentiation process were compromised by the expression of Scd1-PB1 or Byr2-RBD. Cells harbouring *ras1.G17V* and CRIB-GFP (KT5938) were induced for mating/sexual differentiation by the plate mating assay condition (Materials and Methods), and live cell images of the CRIB-GFP signals were taken 16 hours after induction of the sexual differentiation. Navy arrows indicate cortical CRIB-GFP foci. Cells with the cortical CRIB-GFP foci (marked by an orange star) are counted as “cortical CRIB-GFP positive” cells. The scale bar is 10 µm. (C) Quantitation of the results presented in (B). 300 cells were scored for cell deformity (left panel) and the cortical CRIB-GFP signal (right panel). % cells is presented. The experiment was repeated three times, and the mean and SD values of the % cells are plotted in the graphs. Ordinary one-way ANOVA followed by *post hoc* Dunnett’s multiple comparisons test showed that the expression of scd1-PB1 or byr2-RBD reduces the % deformed cells and % cells with cortical CRIB-GFP. (D) GTP-loaded Ras1.G17V (1-172) directly binds to Byr2 (65-180) and Scd1 (760-872). *In vitro* GST pull-down assays of bacterially expressed Ras1.G17V (1-172), GST-Byr2 (65-180) and GST-Scd1 (760-872) were conducted as described in materials and methods. GTP-loaded Ras1.G17V (1-172) was found to bind to both GST-Byr2 (65-180) and GST-Scd1 (760-872). (E) Two Ras1 effectors, Byr2 and Scd1, compete for GTP-loaded Ras1.G17V (1-172). *In vitro* GST pull-down assays of bacterially expressed Ras1.G17V (1-172) and GST-Byr2 (65-180) were conducted as in (D). The addition of the Scd1 (760-872) fragment interfered with Ras1-Byr2 binding (the 4^th^ lane). Quantitated signal intensities of the Ras1.G17V (1-172) band in the gel are shown in the right panel.

When Cdc42 activation was examined in the *ras1.G17V* cells, overexpression of Scd1-PB1 reduced the cortical CRIB-GFP signal and abolished deformed cells (Fig. 5B and C), indicating that the endogenous Scd1 function was interfered with the over-expressed Scd1-PB1. Byr2-RBD overexpression also resulted in a comparable phenotype (Fig. 5B and C). Collectively, two Ras1 effectors, MAPKKK^Byr2^ and Cdc42-GEF^Scd1^, compete with each other for Ras1 *in vivo*.

Next, we conducted *in vitro* binding competition assays using bacterially expressed GST-tagged fragments of Byr2-RBD and Scd1-PB1, both of which bound Ras1.G17V (1-172) loaded with GTP (Fig. 5D). The binding of Ras1.G17V^GTP^ to GST-Byr2-RBD was substantially decreased when the GST-Scd1-PB1 was added (Fig. 5E). The result indicates the biochemical competitive nature of MAPKKK^Byr2^ and Cdc42-GEF^Scd1^ for active Ras1.

### Ras1 and an adaptor protein Ste4 are both necessary to fully activate MAPKKK^Byr2^

Although Ras1 plays a significant role in activating MAPK^Spk1^, a low but detectable level of **pp**MAPK^Spk1^ was still induced in the *ras1Δ* mutant (Fig. 3D and Supplementary Fig. S8A), indicating that there is a Ras1-independent mechanism to activate MAPK^Spk1^. Previous studies proposed an adaptor protein, Ste4, to be involved in the activation of MAPKKK^Byr2^ ^33, 39, 40, 55^. We therefore examined whether Ste4 is required for MAPK^Spk1^ activation.

We detected virtually no MAPK^Spk1^ phosphorylation in the *ste4Δ* mutant (Fig. 6A, yellow line and Supplementary Figure S8C), revealing that the adaptor^Ste4^ is a prerequisite for the MAPK^Spk1^ activation. The introduction of *ras1.G17V* mutation neither restored the MAPK^Spk1^ activation nor mating (Fig. 6A, B). Thus, activation of Ras1 cannot take over Ste4 function. In a striking contrast, the *ste4Δ MAPKK^byr1.DD^* double mutant induces the constitutive MAPK^Spk1^ activation and the “*fus”* phenotype as the *MAPKK^byr1.DD^*single mutant cells (Fig. 6A and B). Collectively, a full MAPKKK^Byr2^ activation requires both Ras1 and Ste4 functions. Interestingly, Ste4’s role is limited to the MAPKKK^Byr2^ activation process as MAPKKK^Byr1.DD^ can suppress the *ste4Δ* mutant phenotype, whereas MAPKKK^Byr1.DD^ could not bypass the Ras1 function (Fig. 3F).

**Figure 6.**
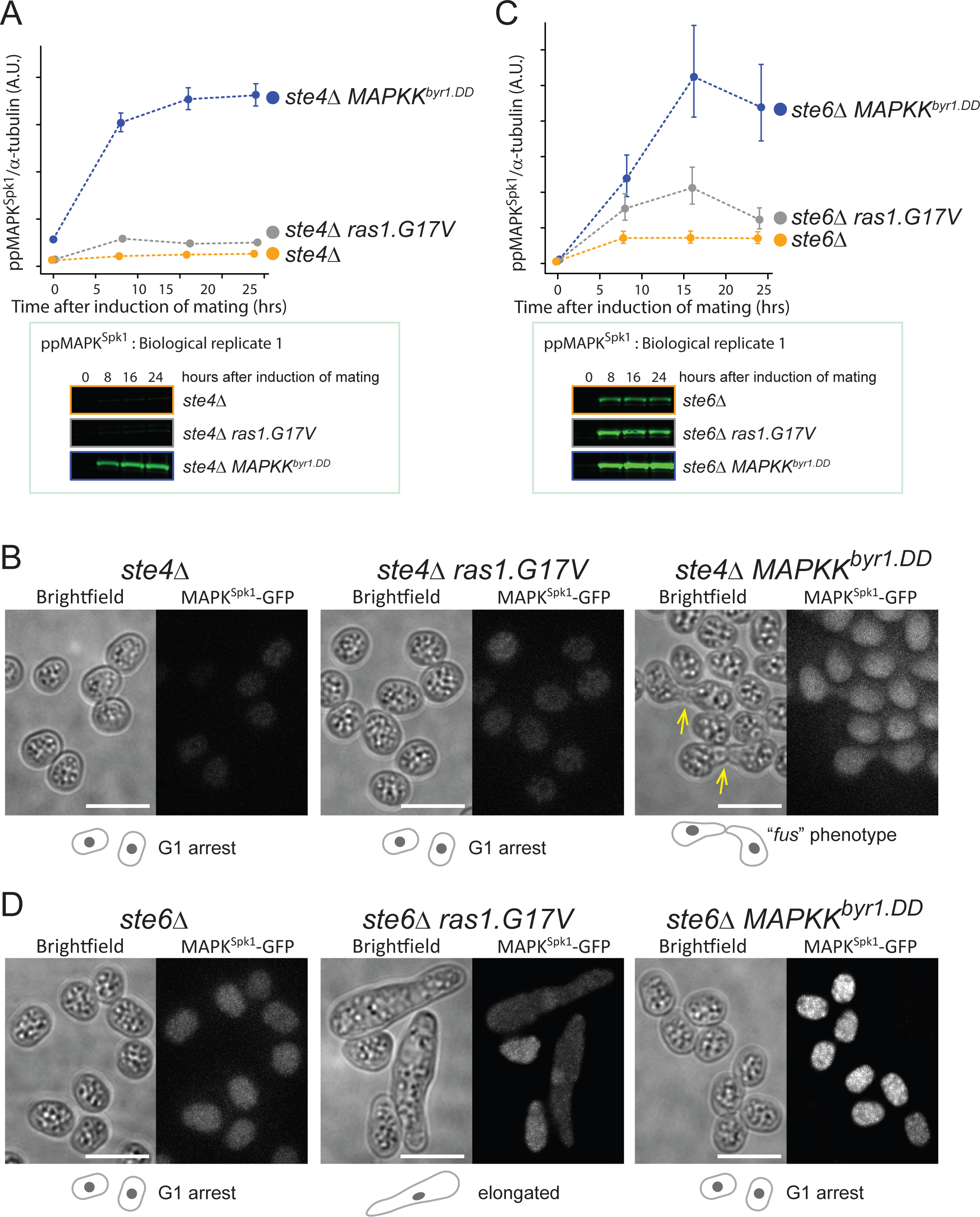
Distinct contributions of Ste4 and Ste6 to MAPK^Spk1^ activation. (A) Ste4 is essential for MAPK^Spk1^ activation. MAPK^Spk1^ phosphorylation status in *ste4Δ* (KT4376), *ste4Δ ras1.G17V* (KT5143) and *ste4Δ MAPKK^byr1.DD^* (KT5136) at times-points 0, 8, 16 and 24 hours after the induction of sexual differentiation were examined. Quantified results of three biological replicates (error bars are ±SEM) are presented. The ppMAPK^Spk1^ Western blotting result of Biological Replicate 1 is presented as a representative result where the ppMAPK^Spk1^ levels are close to 0 in the *ste4Δ* (KT4376) and *ste4Δ ras1.G17V* (KT5143) strains. All the original Western blotting membranes are presented in Supplementary Fig. S8C. (B) The *MAPKK^byr1.DD^* but not *ras1.G17V* mutation suppresses the incapability of *ste4Δ* to cause pheromone-induced morphological change. Cell images of *ste4Δ* (KT4376), *ste4Δ ras1.G17V* (KT5143) and *ste4Δ MAPKK^byr1.DD^* (KT5136) strains were taken 24 hours after induction of mating. The scale bar represents 10 µm. (C) Lack of Ste6 does not result in the complete loss of MAPK^Spk1^ phosphorylation. MAPK^Spk1^ phosphorylation status in *ste6Δ* (KT4333), *ste6Δ ras1.G17V* (KT4998) and *ste6Δ MAPKK^byr1.DD^* (KT5139) at times-points 0, 8, 16 and 24 hours after the induction of sexual differentiation were examined. Quantified results of three biological replicates (error bars are ±SEM) are presented. The ppMAPK^Spk1^ Western blotting result of Biological Replicate 1 is presented as a representative result where the ppMAPK^Spk1^ level of the *ste6Δ* (KT4333) is clearly detectable. All the original Western blotting membranes are presented in Supplementary Fig. S8D. (D) The *MAPKK^byr1.DD^* but not *ras1.G17V* mutation suppresses the incapability of *ste6Δ* to cause pheromone-induced morphological change. Cell images of *ste6Δ* (KT4333), *ste6Δ ras1.G17V* (KT4998) and *ste6Δ MAPKK^byr1.DD^* (KT5139) strains were taken 24 hours after induction of mating. The scale bar represents 10 µm.

### Ste6, a Ras1 GTP-GDP exchange factor, contributes to both the MAPK^Spk1^ and the Cdc42 pathway activation

The activation of Ras1 is mediated by two GDP-GTP exchange factors (GEFs), Ste6 and Efc25^56, 57^. Ste6 is essential for mating but is dispensable during vegetative growth, whereas Efc25 is dispensable for mating but is required to maintain cell morphology during vegetative growth^56,57^. There has been an interesting proposition that Ste6 may specifically help Ras1 to activate the MAPK^Spk1^ pathway but not the Cdc42 pathway, whilst Efc25 specifically facilitates Ras1 to activate the Cdc42 pathway^58^. We examined this hypothesis by monitoring the MAPK^Spk1^ activation status and conducting genetic epistasis analysis of *ras1.G17V* and *MAPKK^byr1.DD^* in the *ste6Δ* mutant.

In the *ste6Δ* cells, MAPK^Spk1^ phosphorylation occurred at a detectable level (Fig. 6C yellow line and Supplementary Figure S8D). When the *MAPKK^byr1.DD^* mutation was introduced, the MAPK^Spk1^ phosphorylation level was maximized (Fig. 6C and Supplementary Figure S8D) and the nucleus MAPK^Spk1^-GFP signal accumulated (Fig. 6D). However, the sterile phenotype remained (Fig. 6D). In contrast, the *ras1.G17V* mutation rescued the “pheromone-insensitive sterile” morphology of *ste6Δ*, as previously reported^56^, exhibiting the “elongated” phenotype (Fig. 6D) even though the increase of the ppMAPK^SPK1^ accumulation was not as high as the case of the *MAPKK^byr1.DD^*mutation (Fig. 6C and Supplementary Figure S8D). The result indicates that, unlike the *ste4Δ* mutant, the mating deficiency of *ste6Δ* is not caused by a mere lack of MAPK^Spk1^ activation but by a lack of Ras1 activation, which mediates *both* the MAPK^Spk1^ and Cdc42 pathway activation in response to the pheromone signalling.

### Activation of Gpa1 represents the pheromone signalling

Gpa1, which plays the primary role in pheromone signalling^21^, is expected to act upstream of Ste4 and ste6. In agreement, in the *gpa1Δ* mutant, we detected no MAPK^Spk1^ activation nor morphological change (Fig. 7A, yellow line, Fig. 7B, left panel, and Supplementary Fig. S10A). The *ras1.G17V* did not rescue the lack of *MAPK^Spk1^* activation nor did it induce a shmoo-like morphological change (Fig. 7A, grey line, Fig. 7B, the second right panel, and Supplementary Fig. S10A), supporting our earlier observation that a Ras1-independent mechanism, involving Ste4, is essential for MAPK^Spk1^ activation. Meanwhile, the *MAPKK^byr1.DD^*caused the constitutive activation of MAPK^Spk1^ (Fig. 7A, blue line and Supplementary Fig. S10A), but cells showed no morphological change (Fig. 7B, the second left panel). When both *ras1.G17V* and *MAPKK^byr1.DD^* mutations were introduced into the *gpa1Δ* strain, MAPK^Spk1^ was activated, and a shmoo-like morphological change occurred (Fig. 7A, red line, Fig. 7B, the right panel and Supplementary Fig. S10A). Therefore, the activation of both the MAPK^Spk1^ pathway and the Ras1 pathway is sufficient to take over the Gpa1 function.

**Figure 7.**
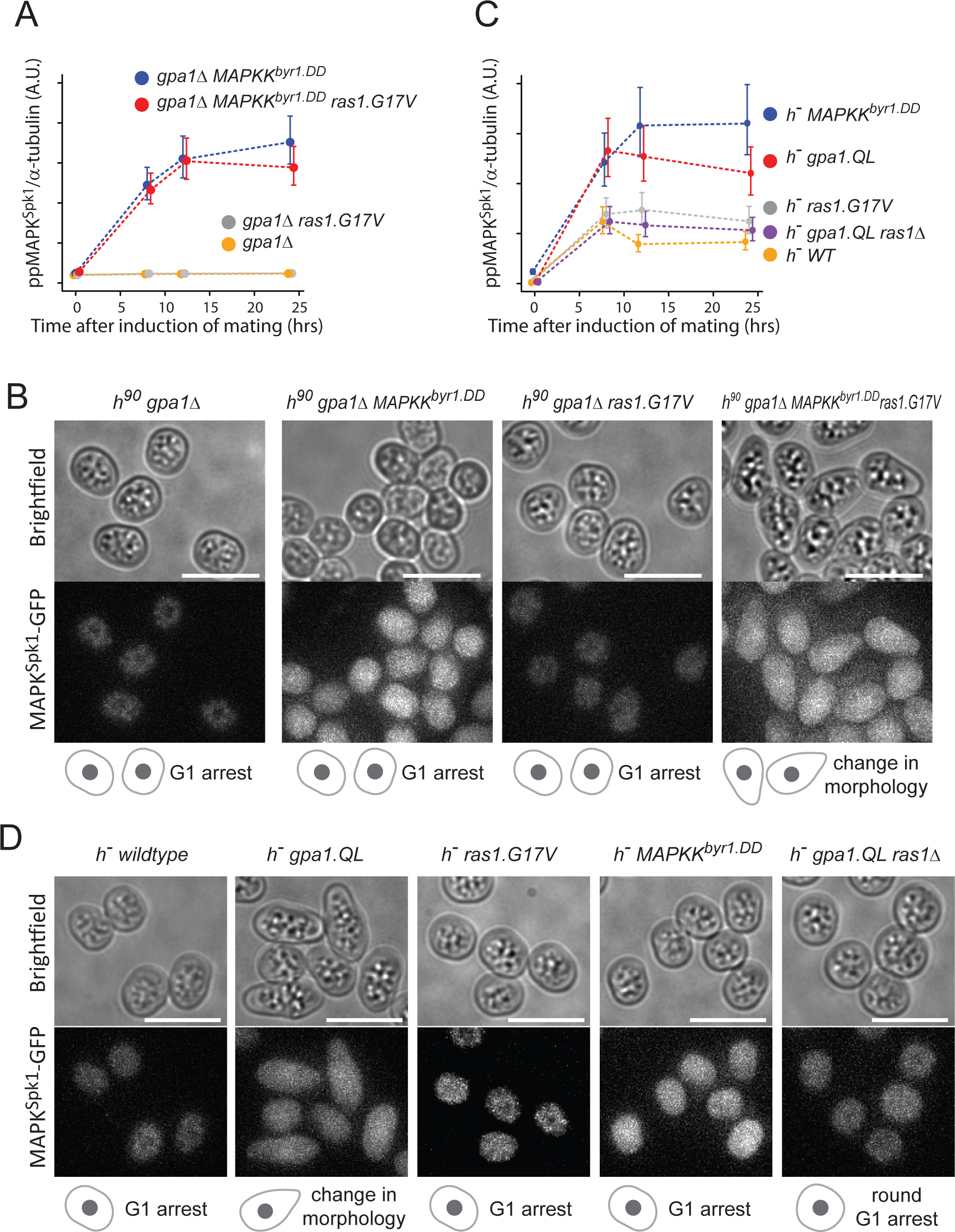
Gpa1 plays the central role of the pheromone signal transduction by activating both MAPK^Spk1^ and Ras1 pathways. (A) MAPK^Spk1^ phosphorylation status in homothallic *gpa1Δ* (KT4335), *gpa1Δ ras1.G17V* (KT5023), *gpa1Δ MAPKK^byr1.DD^* (KT4353) and *gpa1Δ ras1.val17 MAPKK^byr1.DD^* (KT5035) at times-points 0, 8, 12, 16 and 24 hours after the induction of sexual differentiation were examined. Results of three biological replicates (error bars are ±SEM) are presented. Original Western blotting membranes are presented in Fig. S10A. (B) Cell images of the above-mentioned strains at 16 hours after the induction of sexual differentiation. All the cell images were taken and processed as in Fig. 1. The scale bar represents 10 µm. (C) MAPK^Spk1^ phosphorylation status in *h^-^* WT (KT4190), *h^-^ gpa1.QL* (KT5059), *h^-^ ras1.G17V* (KT4233), *h^-^ gpa1.QL ras1Δ* (KT5070) and h *MAPKK^byr1.DD^* (KT4194) at times-points 0, 8, 12 and 24 after the induction of sexual differentiation were examined. Results of three biological replicates (error bars are ±SEM) are presented. Original Western blotting membranes are presented in Supplementary Fig. S10B. (D) Cell images of the above strains at 12 h after the induction of sexual differentiation. All the cell images were taken and processed as in Fig. 1. The scale bar represents 10 µm.

The signalling hierarchy of Gpa1, Ras1 and the MAPK^Spk1^ cascade was further validated in a heterothallic *h^-^* cell population. This population serves as a reference for measuring responses to nitrogen starvation without mating factor signalling as it lacks a mating partner and, consequently, the mating pheromone^41^. Interestingly, upon nitrogen starvation, the *h^-^* wildtype cells displayed a basal level of **pp**MAPK^Spk1^ activation, although no morphological changes were observed (Fig. 7C, yellow line, Fig. 7D, the left panel, and Supplementary Fig. S10B). In contrast, *h^-^* cells with the constitutively active *gpa1.QL* mutation exhibited a “shmoo-like” morphological change as previously reported^21^ and showed strong MAPK^Spk1^ activation (Fig. 7C, red line, Fig. 7D, the second left panel and Supplementary Fig. S10B). Conversely, the *h^-^ ras1.G17V* mutant showed no apparent morphological alternation and exhibited only a basal level of **pp**MAPK^Spk1^, comparable to that observed in the *h^-^* wildtype strain (Fig. 7C, grey line, Fig. 7D, the middle panel, and Supplementary Fig. S10B). Meanwhile, the *h^-^ MAPKK^byr1.DD^* mutant induced strong constitutive MAPK^Spk1^ activation, confirming that the MAPKK^Byr1.DD^ can activate MAPK^Spk1^ regardless of the pheromone signal input (Fig. 7C, blue line and Supplementary Fig. S10B). Yet, the cell morphology remained unchanged (Fig. 7D, the second right panel). Collectively, these results support the model in which Gpa1 serves as the central transducer of the pheromone signalling. It was noted that the Gpa1^QL^-induced phenotype was largely dependent on Ras1 function as the *h^-^ gpa1.QL ras1Δ* double mutant exhibited only a basal level of **pp**MAPK^Spk1^, and round cell morphology (Fig. 7C, purple line, Fig. 7D, the right panel and Supplementary Fig.

S10B).

### Delayed negative feedback explains the transient increase of the total and phosphorylated MAPK^Spk1^ in ordinary differential equations-based mathematical modelling

Our results, along with previous findings, can be summarised in a diagram illustrating that the pheromone signalling activates Ras1, leading to the activation of two competing effectors, MAPKKK^Byr2^ and Cdc42-GEF^Scd1^ (Fig. 8A). In addition, the MAPKKK^Byr2^ activation also requires Ste4 function, and a reciprocal activation occurs between MAPK^Spk1^ and Cdc42 for their full activation (Fig. 8A). We further explored the regulatory relationships among the signalling components that influence the levels of total MAPK^Spk1^ ([tSpk1]), **pp**MAPK^Spk1^ ([ppSpk1]) and active Cdc42 ([aCdc42]) by constructing mathematical models and finding optimal parameters for the quantitative Western blot data (Fig. 1C, E, G, Fig. 2A, Fig. 3B and Supplementary Fig. S3) and the CRIB-GFP imaging data (Fig. 4)(see Material and Methods section and Supplementary Figures S11 and S12 for details).

**Figure 8.**
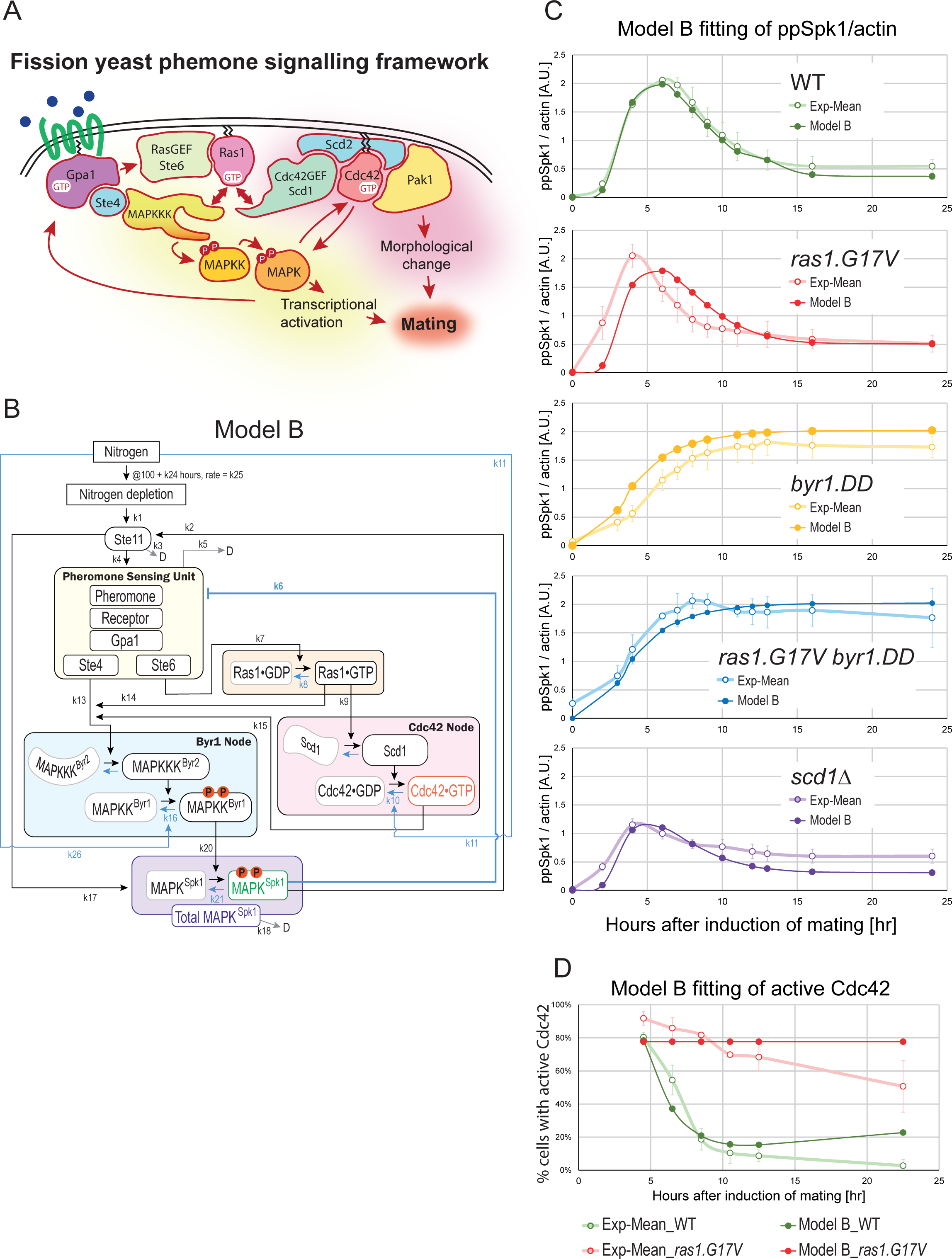
Mathematical modelling of the fission yeast pheromone signalling dynamics. (A) Schematic diagram of the fission yeast pheromone signalling pathway. The diagram highlights the bipartite mechanism to activate MAPKKK^Byr2^, the competition of two Ras1 effectors, MAPKKK^Byr2^ and Cdc42GEF^Scd1^, and the interplay between the MAPK^Spk1^ and Cdc42 pathways. Successful mating requires both the MAPK^Spk1^ and Cdc42 pathway activation. (B) Model B components and framework. For the detailed implementation of the mutants, see Supplementary Fig. S11 and Materials and Methods. The measured components, total MAPK^Spk1^, **pp**MAPK^Spk1^ and Cdc42^GTP^, are shown in purple, green and red, respectively. (C) Model B fittings for the **pp**MAPK^Spk1^ levels in wildtype, *ras1.G17V*, *MAPKK^byr1.DD^*, *ras1.G17V MAPKK^byr1.DD^* and *Cdc42GEF^scd1^Δ* mutants. Mean values (open circles) and SD values (error bars) of the experimental results are shown in pale colours, and the model-fitted values are shown as filled circles in darker colours. Model B fittings for the total MAPK^Spk1^ are shown in Supplementary Fig. S13. (D) Model B fitting of the active Cdc42 levels in wildtype and *ras1.G17V* mutants. Mean values (open circles) and SD values (error bars) of the experimental results are shown in pale colours, and the model-fitted values are shown in filled circles in darker colours.

We postulated up to 29 biochemical processes accompanied by a rate parameter *k* to represent the documented relationships among the signalling components (Supplementary Table S1, Fig. 8B, Supplementary Fig. S11, S12A and Supplementary Fig. S14A; for further details, see Material and Methods section). We obtained initial parameter values using an optimisation toolbox of MATLAB software (Mathworks, Natick, MA, USA) and subsequently fitted these values to better align with the experimental results through the use of a Markov chain Monte Carlo method (MCMC)^59^.

Our initial model, Model A, did not yield a set of parameters that aligned with the experimental results (Supplementary Fig. S12); [tSpk1] and [ppSpk1]did not decrease following an initial increase. To replicate the transient increase of [tSpk1] and [ppSpk1], a delayed negative feedback regulation was deemed necessary^60^. Given that the *MAPKK^byr1.DD^* mutant entirely lacks downregulation (Fig. 1E), we considered downregulations occurring downstream of MAPKK^byr1^ (such as Pyp1 and Pmp1) to be physiologically insignificant. On the other hand, Sxa2 (a serine carboxypeptidase against a mating pheromone P-factor) and Rgs1 (a regulator of Gpa1), both of which are induced upon successful pheromone signalling^41, 61, 62, 63^, receptor internalization^64^ and regulation of the *mapk^spk1^* transcript or other components by antisense RNA^65^ fit well to the criteria for the delayed negative feedback. We consolidated all these elements collectively as a single circuit and provided it with the rate constant *k6* (Model B, Fig. 8B). Incorporating this negative feedback allowed us to obtain a set of parameters that effectively reproduced the transient increase of [tSpk1] and [ppSpk1]in wildtype, *ras1.G17V* and *scd1Δ* condition (Fig. 8C and Supplementary Figure S13). Furthermore, the fit for the active Cdc42 was also significantly improved (Fig. 8D).

### Hypothetical regulations to further improve the model fit to the experimental results

Model B phenocopied the experimental results of tSpk1 and **pp**Spk1 in wildtype and *scd1Δ* strains and qualitatively reproduced most data points in other strains. However, it did not predict the early increase of the **pp**Spk1 levels in the *ras1.G17V* and *ras1.G17V MAPKK^byr1.DD^* strains (Fig. 8C).

We revised Model B to generate Model C, where we introduced two aspects (Supplementary Fig. S14). First, we added *ras1.G17V* strain-specific negative regulatory paths from **pp**Spk1 to modulate *k_9_* and *k_14_*, the rate constants of two Ras1 downstream reactions. Second, to distinguish the outcomes between *MAPKK^byr1.DD^* and *MAPKK^byr1.DD^ ras1.G17V* mutants, we added a hypothetical positive regulation of **pp**Spk1 by Cdc42 with a rate constant *k*27. As a result, model C reproduced both (i) the earlier **pp**Spk1 peak with the height comparable to the WT in the *ras1.G17V*, and (ii) the difference between *MAPKK^byr1.DD^*and *MAPKK^byr1.DD^ ras1.G17V* strain (Supplementary Fig. S14C). The modeling analyses suggested that the constitutively active form of Ras1, Ras1.G17V, may have a qualitatively different biochemical activity compared to the wildtype Ras1.

### Prediction ability of our quantitative model

To examine the validity of Models B and C, we tested whether the models can predict the **pp**Spk1 levels of other experimental results, presented in Figures 3D, 3E, 6A, 6C, 7A and 7C, involving 20 strains of different genotypes (Supplementary Fig. S15). Both Models B and C faithfully simulated Fig. 3E and 6A, involving *MAPKK^byr1.DD^*, *byr2Δ*, *ste4Δ* and *ras1.G17V*. Model B also produced a simulated outcome highly comparable to the results presented in Fig. 3D and 7A, involving *MAPKK^byr1.DD^*, *ras1Δ*, *gpa1Δ*, and *ras1.G17V*, whereas Model C’s predictions for these experiments were less quantitative. On the other hand, Model C predicted Fig. 7C results, involving heterothallic strains of wild type, *MAPKK^byr1.DD^*, *gpa1.QL*, *ras1.G17V* and *ras1Δ*, better than Model B.

It was noted that both Model B and C predictions for Fig. 6C experiment involving the strains with the *ste6Δ* mutation were the least accurate among all the tested experiments, although Model B correctly predicted high **pp**Spk1 for *ste6Δ MAPKKK^byr1.DD^* and low **pp**SPK1 for *ste6Δ* and *ste6Δ ras1.G17V* at a later stage of about 15 hours after induction of mating and Model C predicted a low **pp**Spk1 level in *ste6Δ*, and a high **pp**Spk1 level in *ste6Δ MAPKKK^byr1.DD^*.

In summary, both models predicted the dynamics of **pp**Spk1 to a good extent in the 20 strains. However, Model C did not show a significantly improved prediction capability compared to Model B, despite two additional modelling parameters, *k28* and *k29.* Therefore, the simpler Model B would represent the core framework of fission yeast pheromone signaling.

## Discussion

By quantitating the MAPK^Spk1^ and Cdc42 activation status during the mating process and conducting epistasis analysis between various signalling mutants, we established that Ras1 coordinates activation of two downstream pathways, the MAPK^Spk1^ cascade and the Cdc42 pathway, and revealed that the *ras1.G17V* mutant phenotype (elongated shmoo formation) is caused by the prolonged activation of Cdc42, rather than constitutive activation of MAPK^Spk1^. We built a mathematical model, incorporating assumptions that two Ras1 effectors, Cdc42-GEF^Scd1^ and MAPKKK^Byr2^, compete for active Ras1, and there is a delayed negative feedback for the MAPK^Spk1^ axis. The bipartite mechanism to activate MAPKKK^Byr2^ and the interplay between the MAPK^Spk1^ and Cdc42 pathways are also implemented in the model. The model faithfully recapitulates MAPK^Spk1^ and Cdc42 activation profiles in the wildtype and all mutant strains examined in this study.

The attenuated MAPK^Spk1^ activation in the presence of Ras1.G17V indicated that an efficient feedback mechanism is in place to counteract the effect of Ras1.G17V. The same trend has been reported in the mammalian models where oncogenic *KRAS* was expressed at the physiological levels ^9, 10, 11^. Therefore, it is likely that the MAPK cascade attenuation is generally robust against upstream constitutive RAS signalling. As the *MAPKK^byr1.DD^* mutant caused constitutive activation of MAPK^Spk1^, the effective negative regulation likely occurs upstream of, or at the same level as, MAPKK^Byr1^. In humans, ERK is shown to phosphorylate RAF proteins, the prototype MAPKKKs, to contribute to ERK signal attenuation^66, 67, 68^. Whether fission yeast MAPK^Spk1^ can directly downregulate MAPKKK^Byr2^ is an important future question.

We show that the adaptor^Ste4^ is a prerequisite for MAPK^Spk1^ pathway activation. Hence, the adaptor^Ste4^ can be a target of the negative feedback loop against **pp**MAPK^Spk1^. This mechanism is shared by budding yeast, where Ste50, a Ste4 orthologue, modulates MAPKKK^Ste11^ ^69, 70^. In humans, although an obvious Ste4/Ste50 orthologue is missing, multiple RAF-interacting proteins, including 14-3-3 proteins, as well as the formation of heterodimers between BRAF and CRAF, have been studied for their Ras-independent mechanism to activate RAF proteins^71^. Collectively, MAPK cascades retain a conserved feature of being resistant to oncogenic RAS mutations at physiological conditions.

Meanwhile, we found that MAPK^Spk1^ activity is required for the Ras1.G12V-induced elongated shmoo formation (Fig. 2C), and in turn, the Cdc42 function is also required for full activation of MAPK^Spk1^ (Fig. 3B), as previously proposed^33^. Therefore, the two Ras1 downstream pathways, Cdc42 and MAPK^Spk1^, are not entirely separable. The situation is reminiscent of the *K-ras^G12D^*MEFs that show morphological anomalies without hyperactivation of ERK and AKT^9^, where the MEFs still responded to inhibitors against MAPK and PI3K pathways, reverting the morphology to the one similar to wildtype^9^. It could be that a basal level of MAPK and PI3K pathway activation may be a prerequisite for the *K-ras^G12D^*–induced morphological anomalies.

The molecular mechanism of how MAPK^Spk1^ contributes to Cdc42 activation will require further studies. As MAPK^Spk1^ activates the master transcriptional regulator Ste11^41, 42, 72^, which up-regulates the pheromone signalling components, at least a part of the mechanism likely occurs through transcriptional regulation. In addition, the localisation of signalling components may be regulated by MAPK^Spk1^. Interestingly, during vegetative growth, a stress-activated MAPK Sty1 negatively regulates Cdc42 even when protein synthesis was inhibited by cycloheximide, indicating direct phosphorylation events of Cdc42 and/or its regulators^73^. In budding yeast, MAPK^Fus3^ brings Cdc24, the GEF for Cdc42, to the shmoo site by phosphorylating an adaptor protein Far1^74^, which otherwise sequesters Cdc24 into the nucleus^75, 76^. Fission yeast does not have an obvious Far1 orthologue, but MAPK^Spk1^ may also directly phosphorylate Cdc42 regulatory proteins such as Cdc42-GEF^Scd1^, Scd2 or GAP-Cdc42^Rga4^, all of which function at the shmoo site during the mating^36, 52^(Fig.4 A and B). In agreement with this hypothesis, a transient MAPK^Spk1^ and MAPKK^Byr1^ localisation on the cell cortex was observed during the mating process (Fig. 1)^36^. Localisation of MAPK at the growing cell tips was also observed in other fungi, including *N. crassa* ^77, 78, 79, 80^ and budding yeast, where MAPK^Fus3^ can directly phosphorylate Bni1, a formin that organises actin filaments, to facilitate shmoo formation^81^. Furthermore, a recent phosphoproteome study using cancer cells carrying KRAS oncogenic mutations revealed ERK-dependent phosphorylation of Rho GTPase signalling components^82^. It will be important to fully uncover the role of MAPK at the plasma membrane in regulating cell morphology.

In this study, we have revealed the vital contribution of Cdc42 to induce the *ras1.G17V* phenotype in fission yeast pheromone signalling. The prolonged Cdc42 activation is likely supported by the robust scaffold-mediated positive-feedback on Cdc42^48, 83^. In mammalian systems, small GTPases, such as Cdc42, RalA/B and Rac, play critical roles in oncogenic Ras signalling^13, 14, 15, 16, 17, 18^. Therefore, oncogenic RAS-induced misregulation of small G proteins may be a common basis for the oncogenicity of mutated RAS-induced signalling. Targeting this process may, therefore, be an effective strategy against oncogenic RAS-driven tumourigenesis.

## Materials and methods

### Yeast strains and media

Genotypes of the *Schizosaccharomyces pombe* strains used are listed in the supplementary Table S2. Gene disruption and 2xFLAG-GFP-tagging of genes were performed using the direct chromosomal integration method described previously^84, 85, 86^. Fission yeast media (YE, MM-N and SPA) and basic genetic manipulations are described in^87^.

### Plate mating assay for synchronous mating

In order to induce synchronous mating, we established the “Plate mating assay” system as follows: Cells were grown in YE supplemented with adenine (0.2g/L) until the cell density reaches 8-10 x 10^6^ cells/ml. Cells were then washed with MM-N (1% glucose, supplemented with Leucine 40mg/L) using a filtration unit. These cells were resuspended to a cell density of 0.8-1x10^7^cells/ml in MM-N. In the timecourse experiments, this moment was considered time 0. 8x10^7^cells were re-filtered onto a PDVF membrane of 47 mm diameter (Millipore DVPP04700) in order to evenly place the cells across the defined area. The filters bearing the cells over their surface were carefully transferred onto sporulation agar plates (SPA+Leucine 40mg /L, 50ml SPA agar / 14cm diameter for 6 membranes), cell side up, and incubated at 30°C. During the time course experiments, at each time-point, one membrane, which initially had 8x10^7^cells at time 0, was removed from the SPA plate and placed into 5 ml of ice-cold 20% trichloroacetic acid (TCA) in a 50 ml falcon tube. The tube was shaken vigorously to remove cells from the membrane. The tube was briefly centrifuged, and the membrane was removed from the tube. The tube was then centrifuged for 1 minute at 2000rpm to pellet cells. The supernatant was discarded, cells were resuspended in 1 ml 20% TCA and transferred to a tube capable of storage at -80°C. The cells were pelleted again using a bench top centrifuge, supernatant discarded, and the pellet snap frozen in liquid nitrogen. Tubes were stored at -80°C, ready for protein extraction. A cartoon representation of the outline of the plate mating assay procedure is presented in Supplementary Fig. S1A.

### Preparation of whole cell extracts

This method is based on that described by Keogh lab Protocols (https://sites.google.com/site/mckeogh2/protocols) with a modification to use Urea buffer at the final step to increase protein solubility. All steps were performed on ice unless otherwise stated. Cell pellets (typically 8x10^7^cells) were thawed on ice and resuspended in 250 µl 20% TCA. 250 µl of acid washed glass beads (SIGMA) were added and sampled chilled on ice for 5 minutes. Cells were broken in a FastPrep24 cell beater (MP Biomedicals, speed 6.5 x 1 minute x 3 with 1 minute intervals). Cell extracts were collected by piercing the bottom of tubes with a needle and placing them into collection tubes and centrifuged at 2000 rpm for 1 minute. Beads were washed with 300 µl of 5% TCA and centrifuged at 2000 rpm for 1 minute, this was repeated twice. The contents of the collection tube were transferred to a 1.5 ml tube and centrifuged at 14K rpm for 10 minutes at 4°C. The liquid was discarded and the cell pellet washed with 750 µl of 100% ethanol. Tubes were centrifuged briefly (∼1min 14000rpm) again. Ethanol was completely removed with a pipette. The pellet was then resuspended in 100 µl of Urea buffer (8M urea, 500 mM Tris pH9.4, 1% SDS, 5mM EDTA). The sample was centrifuged and 40 ul of the supernatant was mixed with 80 µl of 3x protein loading buffer ^88^. The sample was then heated for 5 minutes at 95°C and stored at -80°C until use.

### Western Blotting

Protein extracts were subject to SDS-PAGE and were transferred to Immobilon-FL PVDF membrane (Millipore). Membranes were blocked for a minimum of 1-hour in Odyssey Blocking Buffer (OBB) (Li-cor) diluted 1:1 in PBS. Primary antibody incubation was carried out overnight in OBB 1:1 in PBS with anti phospho-p44/42 MAPK (Erk1/2) (Thr202/Tyr204) rabbit monoclonal antibody (#4370 Cell Signaling technology. Used at 1:4000 dilution), anti GFP antibody ((0.4mg/ml), Roche Cat No 11814460001. Used at a 1:2000 dilution), anti FLAG M2 antibody (Sigma F1804, 1μg/μl, used at 1:2000 dilution), anti actin antibody (Life Technologies MA1-744; mouse monocolonal RRID:AB_2223496, 1:2000 dilution) and anti α-tubulin antibody, TAT1 (generous gift from Keith Gull, 1:3000). Membranes were washed 3x10 minutes with 20 ml TBST (50 mM Tris, pH 7.4 – 150 mM NaCl – 0.1% Tween 20) whilst gently shaking followed by secondary antibody incubation (IRDye 680LT goat anti-mouse antibody, Li-cor 926-32211(1.0 mg/ml), 1:16,000 dilution and IRDye 800CW goat anti-rabbit secondary antibody Li-cor 926- 68020 (1.0 mg/ml), 1:16000 dilution in 0.01% SDS, 0.1% Tween 20 in OBB 1:1 PBS) for 1 hour in the dark to prevent fluorophore bleaching. This was followed by 2x 10 minute TBST washes and a 1x 10 minute TBS (50 mM Tris, pH 7.4 – 150 mM NaCl) wash before scanning on the Odyssey CLx infrared Imaging System (Li-cor). Protein quantitation was conducted using either the Image Studio V2.1 software supplied with the Odyssey CLx scanner or Fiji (Image J, ^89^).

### Quantitation of MAPK^Spk1^ phosphorylation

To detect the activated MAPK^Spk1^, whole cell extracts were prepared and Western blotting analysis was carried out using the anti-phospho ERK antibody (#4370 Cell Signalling Technology) that recognises the dual phosphorylation at the conserved TEY motif. Specificity of the antibody against the phosphorylated MAPK^Spk1^ (**pp**MAPK^Spk1^) in the fission yeast cell extracts was confirmed before proceeding with further experiments as shown in Supplementary Fig. S1. As described above in the “**Plate mating assay for synchronous mating**”, during the time course experiments, at each time-point, one membrane, which initially had 8x10^7^cells at time 0, was used to prepare the whole cell extracts. Then an equal volume of each sample was loaded to the Western blotting. In this manner, we observed that the change of the protein level throughout the mating process per a defined starting material. Recent proteiomis studies showed that during the sexual differentiation, both actin and α-tubulin levels stay constant until sporulation starts ^90^. During the sporulation process, actin stays constant whereas the level of α-tubulin decreases ^90^. Therefore, we used actin as an interanal control when the Western blotting involves wildtype cells, which undergo sporulation. We also used α-tubulin as an internal control when sporulation deficient mutant strains were compared each other. The benefit of α-tubulin was that the signal intensity was stronger and therefore, quantitation can be more sensitive. For all Western blotting analysis, three biological replicates of time-course experiments were used.

### Cell Imaging for bright field and Spk1-GFP signals

A Nikon Eclipse Ti-E microscope equipped with CoolLED PrecisExcite High Power LED Fluorescent Excitation System, an Andor iXon EM-DU897 camera and aCFI Plan Apo VC 100x/1.4 objective was used for cell imaging to capture the morphology of cells and the localisation of the GFP-tagged MAPK^Spk1^ during mating time-courses. For cell images presented in Fig. 4, a 2D array scanning laser confocal microscope (Infinity 3, VisiTech) was used (see below). Each time point comprises 15-25 serial images with 0.4 µm intervals along the Z axis taken to span the full thickness of the cell. All of the Spk1-GFP images were deconvolved using the Huygens Essential Deconvolution software (Scientific Volume Imaging). Deconvolved images were Z-projected (maximum intensity), cropped and combined using Fiji ^89^.

### Imaging and quantitation of CRIB-GFP signal

A 2D array scanning laser confocal microscope (Infinity 3, VisiTech) on a NikonTi-E microscope stand equipped with a Hamamatsu Flash 4.0V2 sCMOS camera and a Plan Apo 100x/1.45 objective was used to capture CRIB-GFP signal to examine activation status of Cdc42. For images presented in Fig. 4A, 25 serial Z images with step size of 0.25 µm were taken. Images were deconvolved using Huygens Essential (Scientific Volume Imaging) and maximum intensity projections are presented.

For images presented in Supplementary Fig. S7A, for each image, 30-45 serial images with 0.2 µm interval along the Z axis were taken to span the full thickness of the cell. Using FIJI software maximum intensity Z-projections were created and used for quantitation. Intensity of GFP signal on the cell cortex was measured using FIJI, by applying line measurement with a line width of 1 μm which traced the inner cell cortex. The measurement was done along one of the cell tips that shows a stronger GFP signal as indicated in an example image in Fig. Supplementary Fig. S7B. 40 cells without septum were measured for each strain. Quantified CRIB-GFP intensity traces were analysed in R (R Core Team http://www.R-project.org/). First, rolling averages over +/-5 measurement points were calculated. Next, the X-coordinates of these traces were aligned by their peak intensity. Finally, the average curve from all aligned traces per strain was calculated, and displayed in Supplementary Fig. S7B with respective standard error of the mean curves (dashed lines).

### Glutathione-S-transferase (GST) tag pull-down assays

Fission yeast cDNA fragments encoding Ras1.G17V (1-172), Byr2 (65-180) and Scd1 (760-872) were cloned in the pLEICS1 (Ras1.G17V) and pLEICS4 (Byr2 and Scd1) vectors (Protex [Protein Expression Laboratory], University of Leicester) to produce N-terminally His6-tagged Ras1.G17V (1-172), N-terminally GST-tagged Byr2 (65-180) and N-terminally GST-tagged Scd1 (760-872). Constructs were expressed in *E.coli* BL21 Rosetta cells in LB. All bacteria cell lysates were prepared in phosphate-buffered saline (PBS), pH 7.4.

His6-Ras1.G17V (1-172) was purified using Ni Sepharose 6 Fast Flow (GE Healthcare) and dialysed into buffer A (20 mM Tris-Cl (pH 7.5), 100 mM NaCl, 5mM MgCl2, 1mM β-mercaptoethanol). For GTP loading, buffer A of the Ras1.G17V (1-172) preparation was firstly replaced by Exchange buffer (20 mM Tris-Cl (pH 7.5), 1mM EDTA, 1mM β-mercaptoethanol) using a filtration unit (Amicon Ultra Centrifugal Filter Unit, 10K cut off, Millipore). GDP-GTP exchange reaction was carried out by adding GTP and EDTA to a final concentration of 8 mM and 12 mM respectively and incubating the sample at 37°C for 10 min. The exchange reaction was terminated by adding MgCl2 to a final concentration of 20.5 mM. The resultant GTP-loaded Ras1.G17V (1-172) was washed with buffer A using the filtration unit and immediately used for the GST pull-down assays.

GST-Byr2 (65-180) and GST-Scd1 (760-872) were purified using glutathione sepharose 4B (GE Healthcare). The proteins bound on the glutathione beads were washed with buffer A and used for the GST pull-down assays together with the Ras1.G17V (1-172) preparation. For the competition assays shown in Fig. 5E, Scd1 (760-872) fragment was prepared by cleaving it out from the N-terminal GST, which was bound to the glutathione beads, using TEV protease.

GST pull-down assays were carried out by incubating Ras1.G17V (1-172) (GDP bound or GTP bound) and GST-Byr2 (65-180) or GST-Scd1 (760-872) with or without Scd1 (760-872) fragment at 4 °C for 30 min. Supernatant of the reaction was saved and mixed with the same volume of the 3x loading buffer ^88^ to generate the “unbound” fraction. The remaining beads were washed twice, resuspended in buffer A of the original volume and mixed with the same volume of the 3x loading buffer to generate the “bound” fraction. Protein samples were run on a SDS-PAGE gel and proteins were visualised by InstantBlue protein stain (Expedeon). Quantitation of the intensities of Ras1.G17V (1-172) bands in Fig. 5E was carried out using Fiji software.

### Statistical analysis

Western blotting data presented in Supplementary Figure S2 was quantified as stated in the “Western blotting” section, and individual replicate results are presented in Supplementary Fig. S3 A, B and C. The mean (± SD) of the ppSpk1/actin triplicate data is presented in Fig 1C, 1E, 1G, 2A, and 3B. The data was analyzed by the generalized additive mixed model (GAMM) by fitting GAMMs on temporal variations in the data points (ppSpk1/actin, total Spk1/actin, and ppSpk1/total Spk1) for individual strain using the mgcv package in R. We assumed that the data were distributed according to a normal distribution (identity link). To account for the repeated measurements of data, the models included the genotype of each strain as a random factor. The smoothed curves were back-transformed from the GAMMs and drawn in the figures with twofold standard errors (Supplementary Fig. S3D).

Western blotting data presented in Supplementary Figure S9, which was quantified and presented in Figure 5A, was analyzed by the generalized linear mixed models (GLMMs), which involved gamma distribution (ln-link). The models included three samples, “pRep1-empty”, “pRep1-scd1-PB1”, “pRep1-byr2-RBD”, and “hours after induction of mating” as fully crossed fixed factors. Mixed models were necessary for these models to account for the repeated measurements from each Western blot membrane data.

### Mathematical Modelling

We constructed models with a system of 32 ordinary differential equations (ODEs) describing the amount of 7 molecular components (Ste11, Ste4/6, active Ras1 ([aRas1]), active Cdc42 ([aCdc42]), phosphorylated active MAPKK^Byr1^ ([pByr1]), total MAPK^Spk1^ ([tSpk1]) and phosphorylated **pp**MAPK^Spk1^ ([ppSpk1])), the nutrient status ([N]) and time (t) in 5 different genetic conditions (wildtype, *ras1.G17V*, *MAPKK^byr1.DD^*, *ras1.G17V MAPKK^byr1.DD^* and *scd1Δ*) (Supplementary Fig. S11). All the signalling components were set to be in close proximity, based on our own observation (Figures 1-4) and a series of localization studies^35, 36, 38^.

We postulated up to 29 biochemical processes involving the signalling components (Supplementary Table S1, Fig. 8B, Supplementary Fig. S12A and Supplementary Fig. S14A) to reflect reported relationships among the signalling components. Each of these processes is accompanied by a rate parameter *k*. We first obtained an initial guess of the parameter values using an optimization toolbox of MATLAB software (Mathworks, Natick, MA, USA) and fitted the parameter values to better reproduce the experimental results using a Markov chain Monte Carlo method (MCMC)^59^.

### Modeling of regulatory networks for Spk1

#### The initial model (Model A)

Model A (Supplementary Fig. S12A) consisted of the signaling components as follows: Genes encoding pheromones, receptors, Gpa1, Ste4, and Ste6 are all known to be under the regulation of Ste11, the master transcriptional regulator for meiotic genes^41, 91, 92, 93, 94^ and these components are grouped into the Pheromone Sensing Unit (PSU). During the vegetative growth, the PSU is set to zero. Nitrogen starvation produces active Ste11^93, 95^ (the rate constant *k1*), which induces the PSU (the rate constant *k4*). The PSU activates Ras1 through Ste6^56^ (the rate constant *k7*), and the resultant active Ras1 activates Cdc42 through Scd1^32^ (the rate constant *k9*). In our models, Scd1 and Cdc42 are grouped into one unit, the Cdc42 Node. The PSU also activates MAPKKK^Byr2^ through Ste4^33, 39, 40, 55^ (the rate constant *k13*), which is further modulated by active Ras1 (the rate constant *k14*) and active Cdc42 (the rate constant *k15*)^33^. Activated MAPKKK^Byr2^ then triggers activation of MAPKK^Byr1^ that activates MAPK^Spk1^ (the rate constant *k20*). In our models, MAPKKK^Byr2^ and MAPKK^Byr1^ are grouped into one unit, the Byr1 Node. The activated MAPK^Spk1^ further activates Ste11^42^ (the rate constant *k2*); thus, MAPK^Spk1^ has a positive feedback loop on its own expression via Ste11. Additional regulatory processes included in our models are a negative regulation of MAPKK^Byr1^ by nitrogen^96^ (the rate constant *k26*) and negative regulation of active Cdc42 during vegetative growth (the rate constant *k11*) by the GTPase activating proteins which are expressed at a higher level in the presence of nitrogen^97^. Specifically, Model A consists of the following differential equations for the control (wild-type):

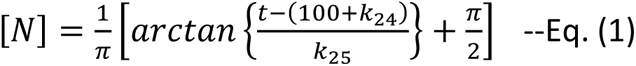

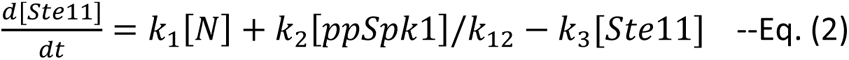

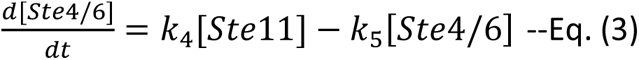

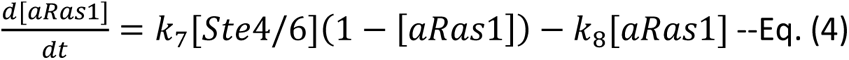

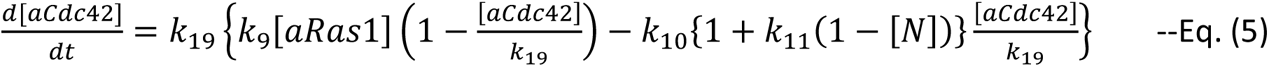

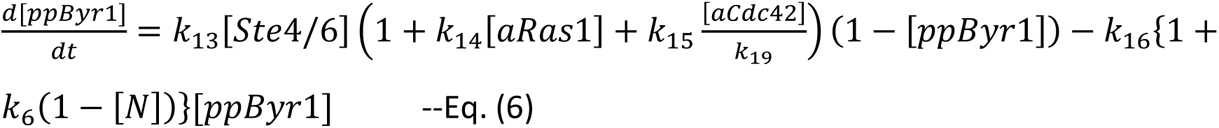

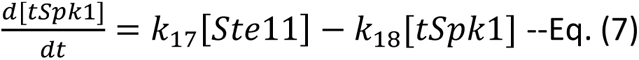

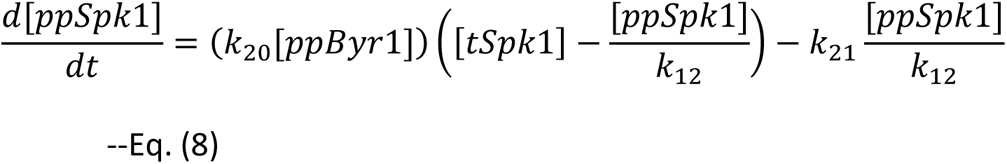

Note that [tSpk1] represents the amount of total Spk1 as the signal intensity of the western blot. Therefore, the amounts of other proteins (e.g. [Ste11]) also represents the relative value normalized to the 1 Unit of [tSpk1], except for [ppSpk1] and [aCdc42]. In case of phosphorylated Spk1, [ppSpk1] represents the amount of phosphorylated Spk1 as the signal intensity of the Western blot of ppSpk1 (but not tSpk1). The amount of phosphorylated Spk1 normalized to the 1 Unit of [tSpk1] will be [ppSpk1]/*k*_12_. In case of active Cdc42, [aCdc42] represents the amount corresponding to the CRIB-GFP [%] (Fig. 4B). The amount of active Cdc42 normalized to the 1 Unit of [tSpk1] will be [aCdc42]/*k*_19_.

For ras1.G17V mutant, we used the same differential equations except that [aRas1] was fixed to a constant value of *k*_22_. Similarly, for byr1.DD mutant, [ppByr1] was fixed to a constant value of *k*_23_. For scd1-delta mutant, [aCdc42] was fixed to zero.

### Model B

In Model B (Fig. 8B), a negative regulation from ppSpk1 to Ste4/6 was added to Model B. Eq. (3’) was used instead of Eq. (3) in Model A.

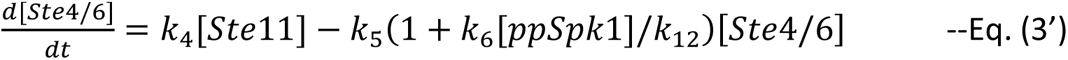

### Model C

In Model C (Supplementary Fig. S14A), three regulations were added to Model B. Two ras1.G17V-specific negative regulations from ppSpk1 to the downstream reactions of Ras1 (i.e. *k*_9_ and *k*_14_) were added to Model B, by using Eqs. (5’) and (6’) instead of Eqs. (5) and (6) only for ras1.G17V and ras1.G17V; byr1.DD conditions. Note that [aRas1] = *k*_22_ and [ppByr1] = *k*_23_ (constant values) in ras1.G17V and byr1.DD conditions, respectively.

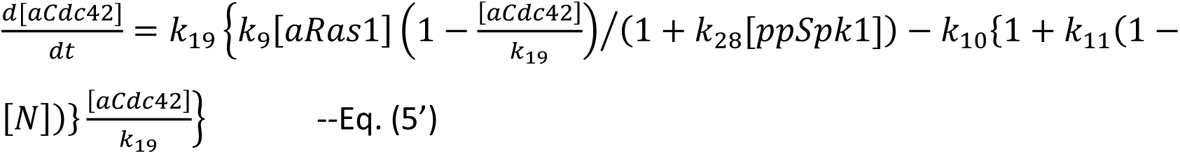

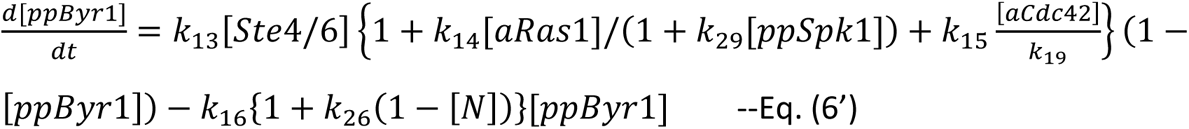

In addition, a regulation to activate aByr1 to ppSpk1 by aCdc42 was added to expect the distinct outcome of byr1.DD and ras1.G17V; byr1.DD as observed in the experiment. Eq. (8’) was used instead of Eq. (8).

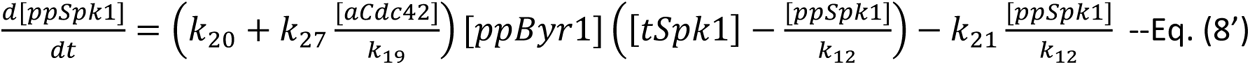

### Initial guess of the parameter values

We first obtained an initial guess of the parameters by using an optimization toolbox of MATLAB software (Mathworks, Natick, MA, USA). We found parameters that optimize an ordinary differential equation (ODE) in the least-squares sense, using the problem-based approach, according to the software website (https://jp.mathworks.com/help/optim/ug/fit-ode-problem-based-least-squares.html?lang=en) within the range of 0.1 ≦ *k*i ≦10, where *k*i represents the *i*-th fitting parameter. The parameters obtained in this way did not reproduce the transient increases of [tSpk1] and [ppSpk1] as observed in our experiment, and thus used as the prior for the MCMC approach. The codes will be uploaded in our Github directory.

### Fitting with an MCMC method

We used a Metropolis algorithm for a Markov chain Monte Carlo (MCMC) method^59^ to search parameters reproducing the experimental behavior using the values of initial guess obtained with MATLAB as the prior. In the algorithm, we change each parameter values by multiplying the previous value (or initial guess for the first step) with exp(*r*), where *r* is a random number following the normal distribution with the mean of 0 and standard deviation of 0.01. We adopted the latter value of 0.01 by testing different values and checked the efficiency of the fitting.

The likelihood of the fitting with a given set of parameters were calculated by the following formulation: 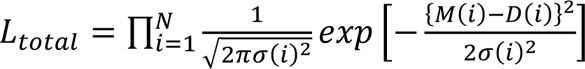, where *N* is the number of data points, *M*(*i*), *D*(*i*) and *σ*(*i*) are the value in the model, the value in the experiments, and the variance in the experiments of the *i*-th data points. If the likelihood *L*_total_ increased from the previous parameters, we adopt the current parameters. Even if the likelihood decreased, we adopt the current parameters with the probability of current *L*_total_ divided by the previous *L*_total_. This step was repeated 100,000 steps. We adopted the parameter set giving the highest likelihood as the best fit parameters. The codes will be uploaded in our Github directory.

## Acknowledgment

The authors thank Tatsuya Maeda, Thibault Mayor, Janni Petersen, Louise Fairall, John Schwabe, David Critchley, Andrey Reviyakin, Gary Willars and Mohan Harihar for helpful suggestions, stimulating discussions and critical reading of the manuscript. We thank Dr. Hiroyuki Kubota (Kyushu Univ) for the discussion on the modeling of signaling networks, Drs. Genta Ueno, Shinya Nakano, Daisuke Murakami, Yusaku Ohkubo (The Institute of Statistical Mathematics) for the discussion on the parameter search methods. We thank PROTEX and PNACL at University of Leicester for their technical assistance. We are grateful to Kazu Shiozaki, Keith Gull and Yeast Genetic Resource Center for providing strains and antibodies. This work was funded by the Wellcome Trust Institutional Strategic Support Fund WT097828/Z/11/Z and WT097828/Z/11/B, the Deutsche Forschungsgemeinschaft (DFG) EXC 81 and Joint Support-Center for Data Science Research, ROIS and JSPS KAKENHI JP18H02414. A.V. was supported by German Academic Exchange Service (DAAD) A0981674.

## Author Contributions

E.J.K., S.R. and K.T. generated yeast strains. E.J.K., G.S. and K.T. monitored activation status of MAPK^Spk1^ and Cdc42. A.V., A.K. and E.K. conducted mathematical modelling. E.J.K., K.R.S. and K.T. conducted image analysis. M.T., R.G., C.P. and C.D. conducted biochemical analysis of Ras1, Byr2 and Scd1. E.J.K., A.V., E.K. and K.T. designed the experiments and interpreted the data.

