## Supplementary Figures, Supplemental Table1 and Supplemental Table 2 for "Constitutively active RAS prolongs Cdc42 signalling while attenuating MAPK signalling during fission yeast mating"

### Supplementary Information

- **Supplementary Figures**
- **Table S1**
- **Table S2**

### Constitutively active RAS prolongs Cdc42 signalling while attenuating MAPK signalling during fission yeast mating

Emma J. Kelsall[1,12], Akatsuki Kimura[2,3,4,12]*, Ábel Vértesy[5,11,12], Kornelis R. Straatman[6], Mishal Tariq[1], Raquel Gadea[1,7], Chandni Parmar[1], Gabriele Schreiber[5], Shubhchintan Randhawa[1,9], Takashi Y. Ida[10], Cyril Dominguez[1,8], Edda Klipp[5]* and Kayoko Tanaka[1]*

Fig. S1

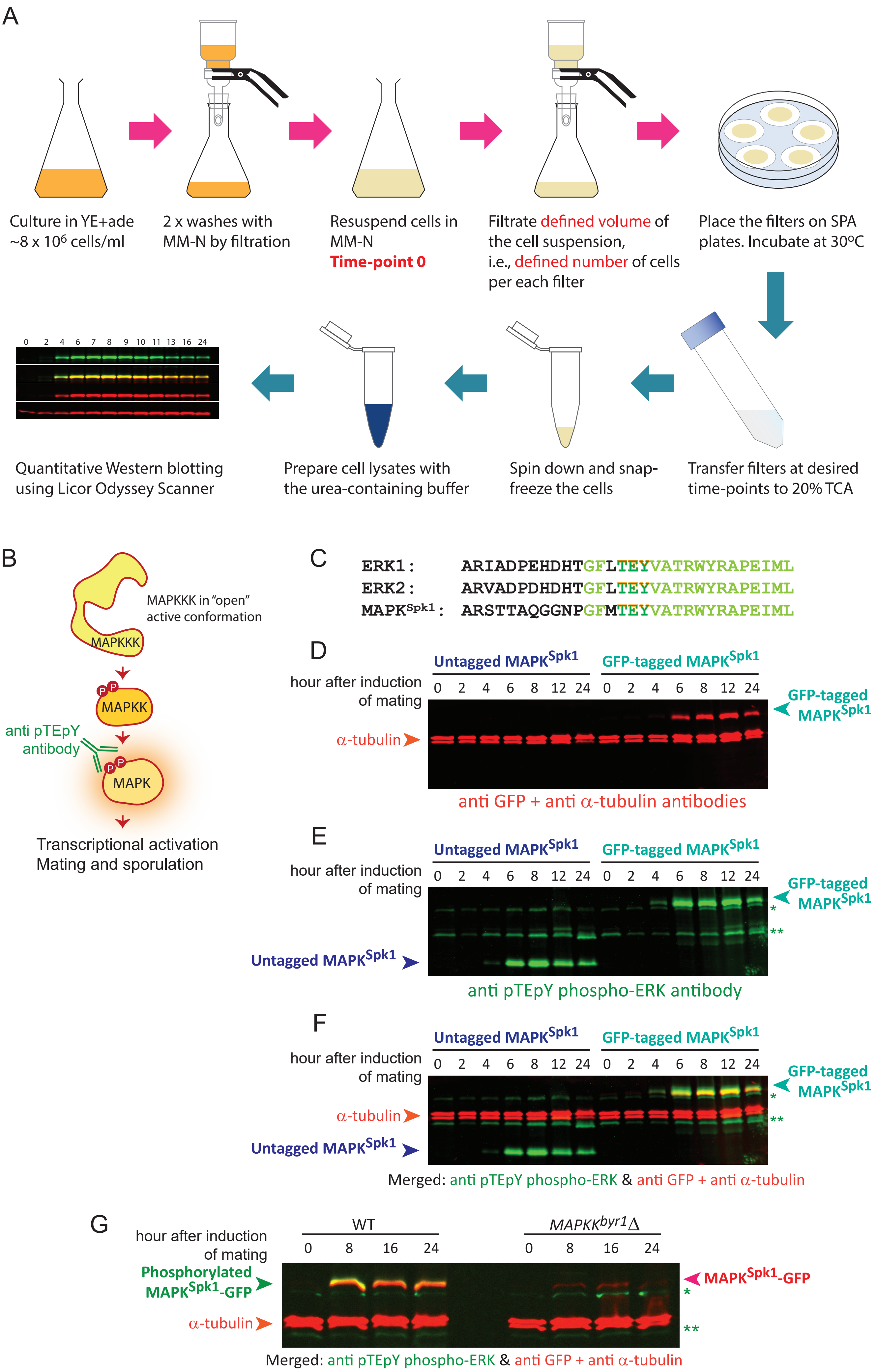
A
Culture in YE+ade ~8 x 10⁶ cells/ml
2 x washes with MM-N by filtration
Resuspend cells in MM-N
Time-point 0
Filtrate defined volume of the cell suspension, i.e., defined number of cells per each filter
Place the filters on SPA plates. Incubate at 30ºC
Transfer filters at desired time-points to 20% TCA
Spin down and snap-freeze the cells
Prepare cell lysates with the urea-containing buffer
Quantitative Western blotting using Licor Odyssey Scanner
0 2 4 6 7 8 9 10 11 13 16 24
B
MAPKKK in "open" active conformation
MAPKKK
MAPKK
anti pTEpY antibody
MAPK
Transcriptional activation
Mating and sporulation
C
ERK1: ARIADPEHDHTGFLTEYVATRWYRAPEIML
ERK2: ARVADPDHDHTGFLTEYVATRWYRAPEIML
MAPKSpk1: ARSTTAQGGNPGFMTEYVATRWYRAPEIML
D
Untagged MAPKSpk1
GFP-tagged MAPKSpk1
hour after induction of mating
0 2 4 6 8 12 24 0 2 4 6 8 12 24
GFP-tagged MAPKSpk1
α-tubulin
anti GFP + anti α-tubulin antibodies
E
Untagged MAPKSpk1
GFP-tagged MAPKSpk1
hour after induction of mating
0 2 4 6 8 12 24 0 2 4 6 8 12 24
GFP-tagged MAPKSpk1
*
**
Untagged MAPKSpk1
anti pTEpY phospho-ERK antibody
F
Untagged MAPKSpk1
GFP-tagged MAPKSpk1
hour after induction of mating
0 2 4 6 8 12 24 0 2 4 6 8 12 24
GFP-tagged MAPKSpk1
*
**
α-tubulin
Untagged MAPKSpk1
Merged: anti pTEpY phospho-ERK & anti GFP + anti α-tubulin
G
hour after induction of mating
WT
MAPKKbyr1Δ
0 8 16 24 0 8 16 24
Phosphorylated MAPKSpk1-GFP
MAPKSpk1-GFP
*
**
α-tubulin
Merged: anti pTEpY phospho-ERK & anti GFP + anti α-tubulin

**Supplementary Figure S1. Plate mating assay and validation of the use of commercially available anti-GFP and anti-phospho-MAPK specific antibodies.**

(A) A graphical representation of the plate mating assay that induces highly synchronous mating/sexual differentiation as stated in the Materials and Methods "Plate mating assay for synchronous mating" section. (B) Schematic outlining of the principal of a phospho-specific MAPK antibody, which specifically recognize the phosphorylated TEY motif (pTEpY) of MAPK. (C) Sequence alignment between mammalian ERK1, ERK2 and *S.pombe* MAPK$^{Spk1}$ at the region of the dual phosphorylation required for activation and surrounding residues. Conserved residues are presented in light green, and TEY is highlighted in orange-green. (D)-(F) A Western blot of a time-course of a WT (KT301) and a MAPK$^{Spk1}$-GFP strain (KT3082) over 24 hours, incubated with anti-GFP, anti-pTEpY phospho-ERK and anti-α-tubulin antibodies. The Li-cor Odyssey system was used to detect the signals. (D) The signals obtained by the 700 nm wavelength scan to detect signals from a primary monoclonal mouse anti-GFP antibody ((0.4mg/ml), Roche Cat No 11814460001. Used at a 1:2000 dilution) and the primary anti-α-tubulin antibody TAT1 (generous gift from Keith Gull, 1:3000), followed by the IRDye 680LT secondary antibody (goat anti-mouse antibody, Li-cor 926-32211(1.0 mg/ml), 1:16,000 dilution). α-tubulin was used as an internal loading control in this instance. (E) The exact same blot as in (D) but showing the 800 nm wavelength scan to detect signals from an anti-pTEpY phospho-ERK antibody (#4370 Cell Signaling Technology. Used at 1:2000 dilution) visualized by the IRDye 800CW goat anti-rabbit secondary antibody (Li-cor 926-68020 (1.0 mg/ml), 1:16000 dilution). (F) An overlay image of the 700 and 800 nm channels that are presented in (D) and (E). (G) The phospho-ERK antibody is phospho-specific. Cell extracts were prepared from the time-course of MAPK$^{Spk1}$-GFP tagged WT (KT3082) and *mapkk$^{byr1}$Δ* strains (KT4300), and MAPK$^{Spk1}$-GFP was detected by anti-GFP (red) while phospho-MAPK$^{Spk1}$ was detected by anti-pTEpY phosphor ERK antibody (green). In the *mapkk$^{byr1}$Δ* strain, phosphorylated MAPK$^{Spk1}$-GFP signal is missing, although MAPK$^{Spk1}$-GFP was detectable at 8, 16 and 24 hours after induction of mating. Note that bands indicated by a single (*) and double (**) asterisks were concluded to be irrelevant to phosphor-MAPK$^{Spk1}$-GFP because they do not exactly overlap with the MAPK$^{Spk1}$-GFP signal (in red) in (F) and (G). Furthermore, these bands exist in non-tagged MAPK$^{Spk1}$ strain seen in (E) and in *mapkk$^{byr1}$Δ* strains in (G), further supporting that the bands are irrelevant to phospho-MAPK$^{Spk1}$-GFP. Considering the molecular weight, the double asterisk band may correspond to another MAPK, MAPK$^{Pmk1}$.

Fig. S2

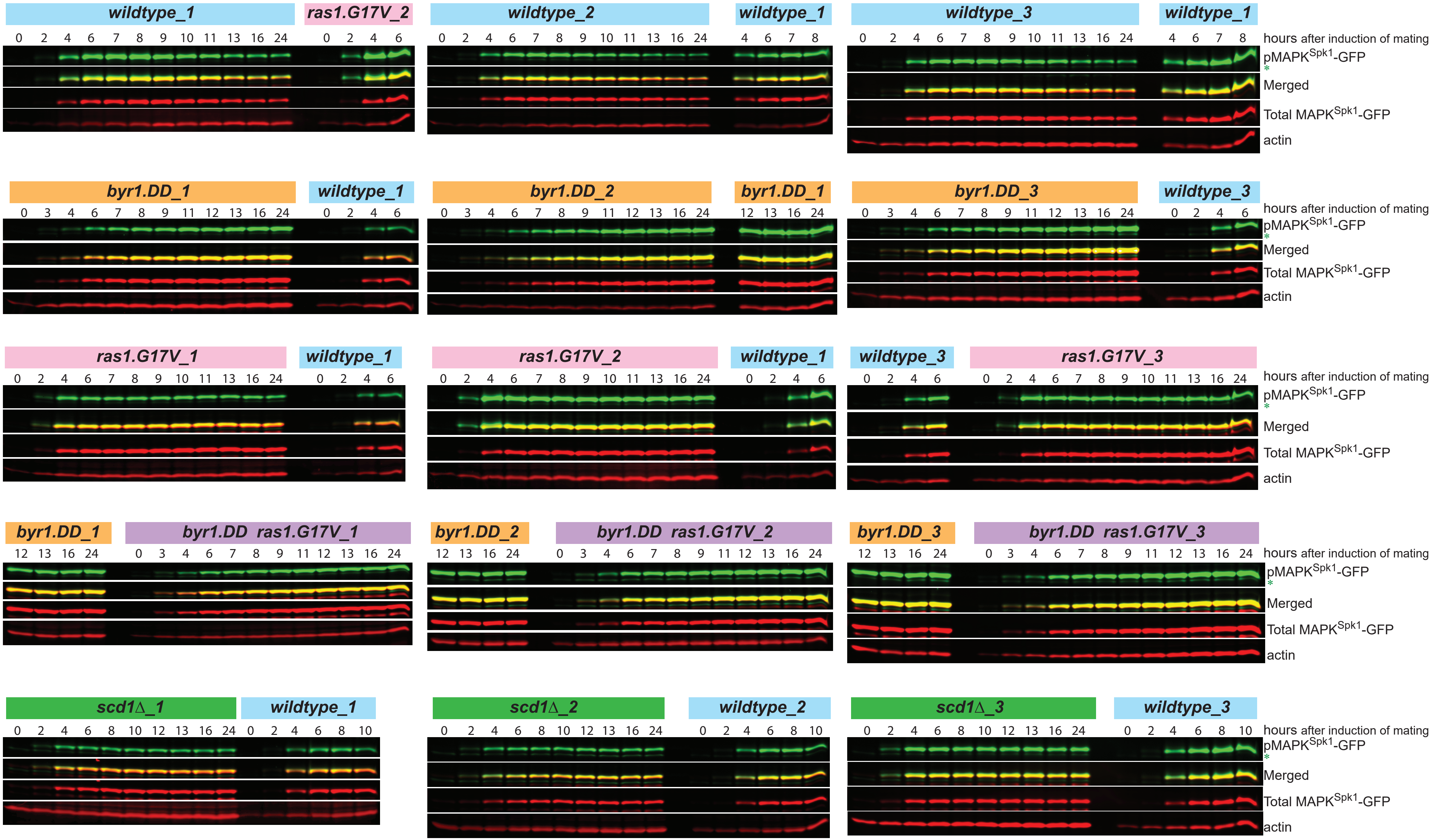

wildtype_1
ras1.G17V_2
wildtype_2
wildtype_1
wildtype_3
wildtype_1
0 2 4 6 7 8 9 10 11 13 16 24
0 2 4 6
0 2 4 6 7 8 9 10 11 13 16 24
4 6 7 8
hours after induction of mating
pMAPK$^{Spk1}$-GFP
*
Merged
Total MAPK$^{Spk1}$-GFP
actin
byr1.DD_1
wildtype_1
byr1.DD_2
byr1.DD_1
byr1.DD_3
wildtype_3
0 3 4 6 7 8 9 11 12 13 16 24
12 13 16 24
ras1.G17V_1
wildtype_1
ras1.G17V_2
wildtype_1
wildtype_3
ras1.G17V_3
byr1.DD_1
byr1.DD ras1.G17V_1
byr1.DD_2
byr1.DD ras1.G17V_2
byr1.DD_3
byr1.DD ras1.G17V_3
scd1Δ_1
wildtype_1
scd1Δ_2
wildtype_2
scd1Δ_3
wildtype_3
0 2 4 6 8 10 12 13 16 24
0 2 4 6 8 10

**Supplementary Figure S2.**

**Original Western blotting images of WT, *byr1.DD*, *ras1.G17V*, *ras1.GV byr1.DD* and *scd1Δ* mutants used to quantify the ppMAPK$^{Spk1}$-GFP levels during the mating process in Fig. 1, 2 and 3**

MAPK$^{Spk1}$-GFP Cells were induced for sexual differentiation by the plate mating assay system as described in Supplementary Fig. S1A and the Materials and Methods. Three biological replicates presented were used for quantitation to obtain the mean value and SEM. Actin was used as a loading control (detected by Life Technologies MA1-744; mouse monoclonal RRID:AB 2223496, 1/2000 dilution), and quantitation was carried out using the Image Studio ver2.1 software (Licor Odyssey CLx Scanner). **pp**MAPK$^{Spk1}$-GFP was detected by anti-phospho-ERK antibody (#4370 Cell Signaling technology. Used at 1:2000 dilution) and total MAPK$^{Spk1}$-GFP-2xFLAG protein level was detected using anti-FLAG antibody (Sigma F1804, 1 µg/ul, 1/2000 ditution). The green asterisk indicates the background band, as seen in the supplementary Figure S1. Numbers indicate hours after induction of mating.

Western blotting quantification of three biological replicates (membranes presented in Supplementary Fig. S2)

Biological replicate 1 — Biological replicate 2 — Biological replicate 3

#### A. ppSpk1/actin (arbitrary unit)

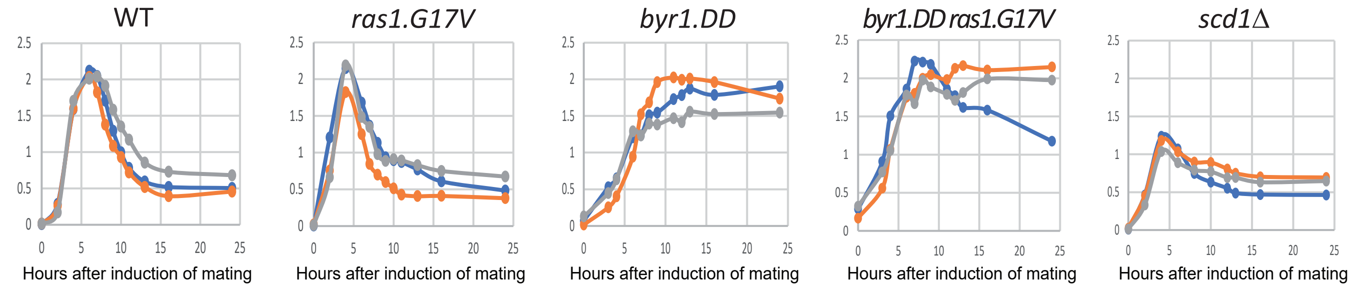


#### B. Total Spk1/actin (arbitrary unit)

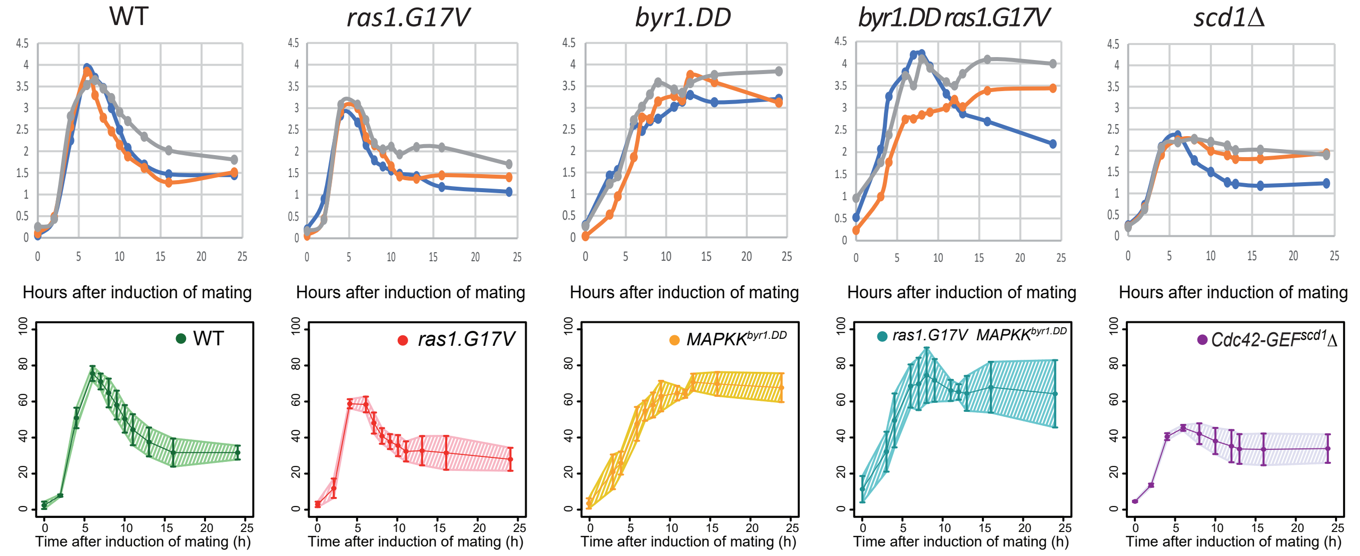


#### C. ppSpk1/total Spk1 (arbitrary unit)

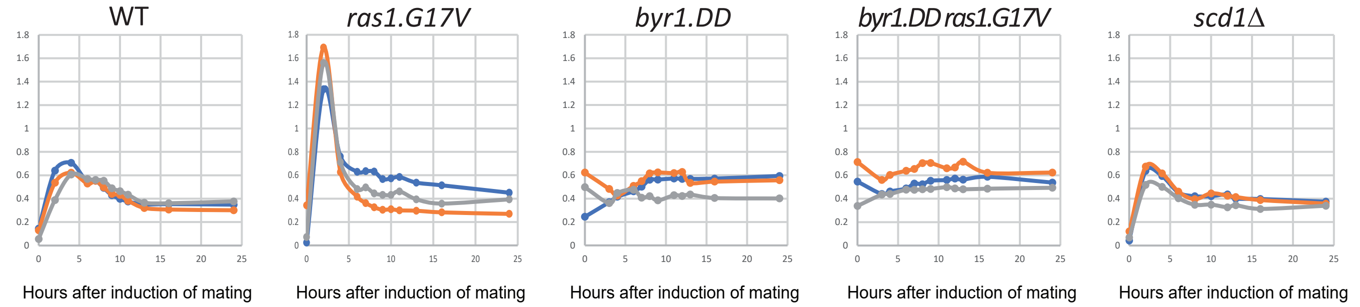


#### D. Generalised additive mixed model (GAMM) fitted to data points

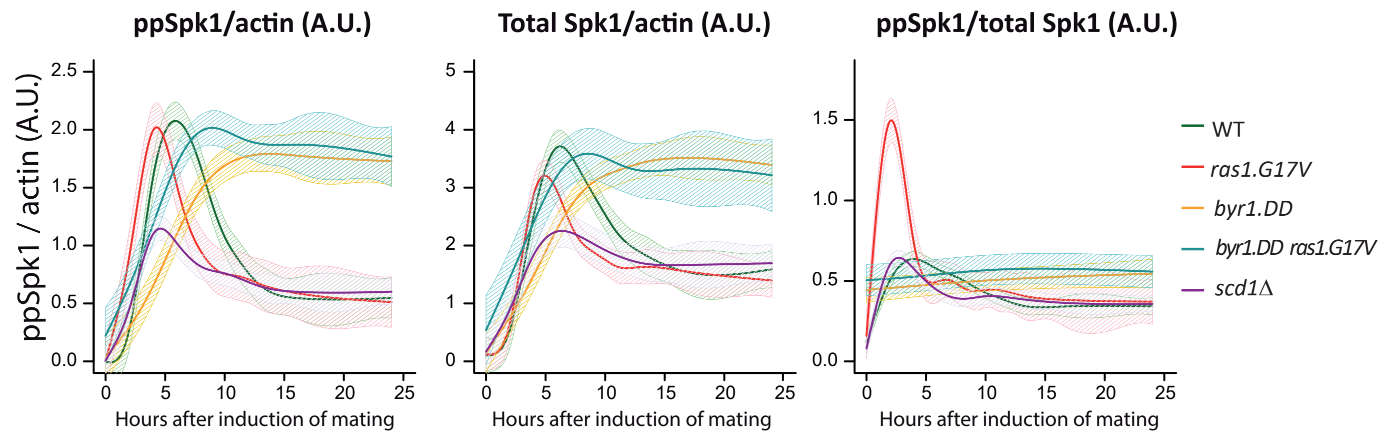


Generalised additive mixed model (GAMM) fitted to data points

**Supplementary Figure S3.**

**Quantification result of each Western blotting membrane of WT, *byr1.DD*, *ras1.G17V*, *ras1.GV byr1.DD* and *scd1Δ* mutants, presented in Supplementary Figure S2.**

(A) and (B). Relative levels of **pp**MAPK$^{Spk1}$-GFP-2xFLAG (A) and total MAPK$^{Spk1}$-GFP-2xFLAG (B) were quantified for each of the Western blotting membranes presented in Supplementary Figure S2. Three biological replicates were made, and each biological replicate data is presented in blue, orange or grey. Quantitation was carried out using the Image Studio ver2.1 software (Licor Odyssey CLx Scanner). The Y axis represents relative signal intensities in an arbitrary unit, which is set to the same scale for all the membranes. Numbers of the X-axis indicate hours after induction of mating. For the **pp**MAPK$^{Spk1}$-GFP-2xFLAG measurement, the mean values ±SEM of the three biological replicates are presented in Figures 1C, 1E, 1G, 2A and 3B. For total MAPK$^{Spk1}$-GFP-2xFLAG measurement, the mean values ±SEM of the three biological replicates are presented in the lower row of Supplementary Figure S3(B). (C) Relative ratios of **pp**MAPK$^{Spk1}$/total MAPK$^{Spk1}$ are deduced from each biological replicate and plotted. (D) Fitted values (and 95% confidence intervals) for the **pp**MAPK$^{Spk1}$ relative levels, total MAPK$^{Spk1}$ relative levels, and the ratios of **pp**Spk1/total Spk1 levels of the five strains under Generalized Additive Mixed Models (GAMM) are presented. The GAMMs indicate that the initial increase of the **pp**MAPK$^{Spk1}$ occurs distinctively earlier in the *ras1.G17V* mutant than in the wildtype strain. The levels of total MAPK$^{Spk1}$ and the **pp**MAPK$^{Spk1}$ in the *byr1.DD ras1.G17V* double mutant also increase distinctively earlier than these levels in the *byr1.DD* single mutant. The GAMMs also show that the **pp**Spk1/total Spk1 ratios increase during the first couple of hours after induction of mating in cells that do not carry the *byr1.DD* mutation. Among these cells, the increase rate in the *ras1.G17V* was distinctively higher than in the wildtype cells. Interestingly, the **pp**Spk1/total Spk1 ratio in the *ras1.G17V* cells quickly dropped to become comparable to the wildtype cells, indicating a robust damping mechanism. In contrast, in the presence of the *byr1.DD* mutant, the **pp**Spk1/total Spk1 ratios stayed almost constant throughout the mating process.

A Wildtype $h^{90}$ cells expressing MAPK$^{Spk1}$-GFP and Smd2-tdTomato (nuclear marker)

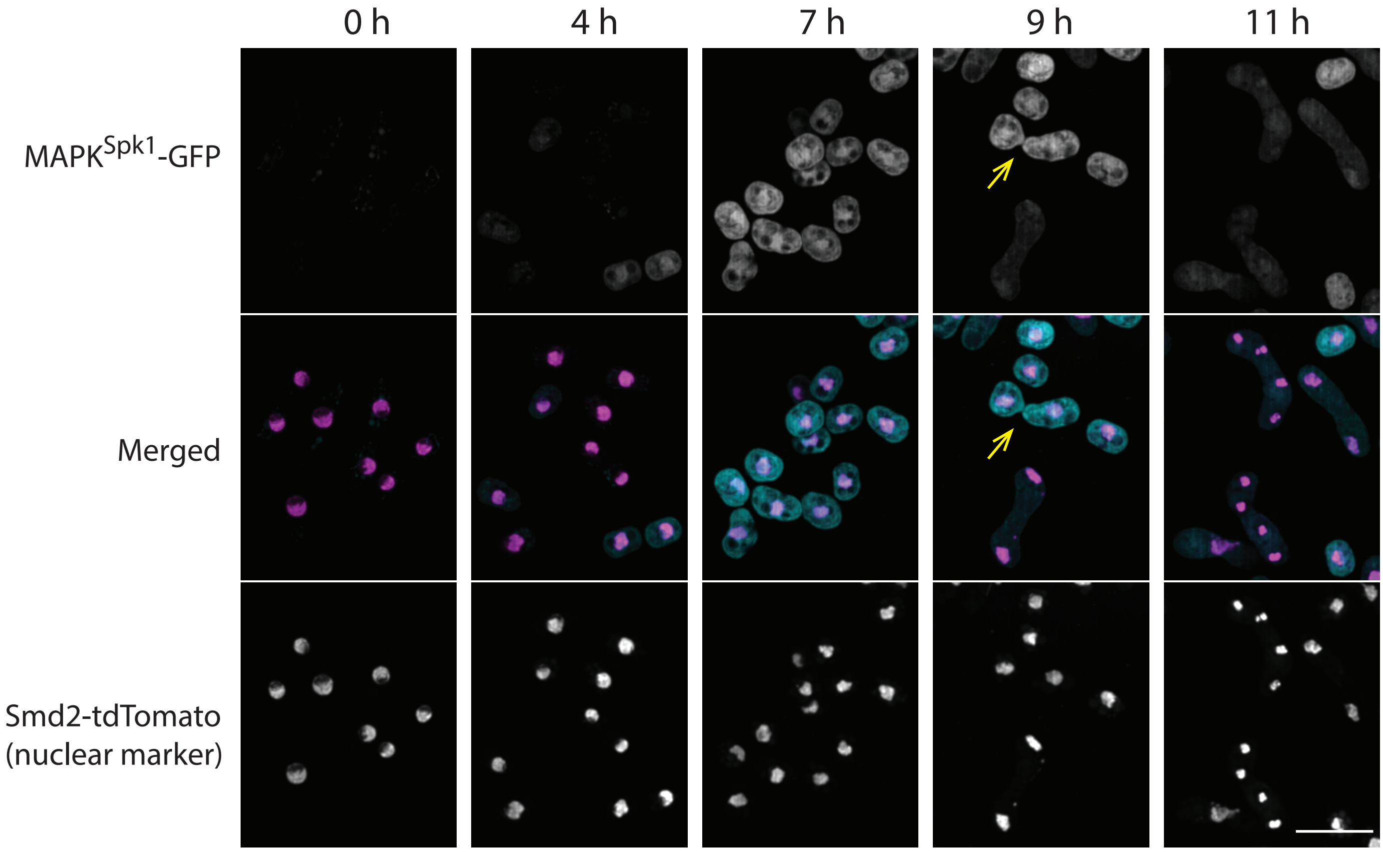


B Serial Z slices showing the MAPK$^{Spk1}$-GFP signal at the shmoo tips of a pair of mating cells

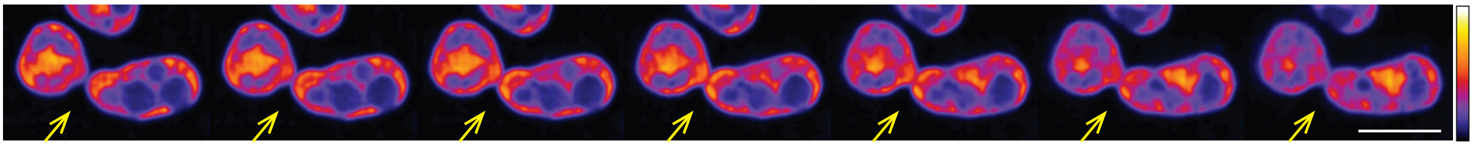

**Supplementary Figure S4. MAPK$^{Spk1}$-GFP signal intensity and localization during the wildtype mating process**

(A) MAPK$^{Spk1}$-GFP appears on the cell cortex and in the nucleus. Wildtype cells expressing MAPK$^{Spk1}$-GFP, together with Smd2-tdTomato (KT5951), a nuclear marker, were induced for sexual differentiation by the plate mating assay system as described in the Materials and Methods and Supplementary Fig. S1A. MAPK$^{Spk1}$-GFP and Smd2-tdTomato signals were captured at the indicated times after induction of sexual differentiation. Images were taken by a 2D-array scanning laser confocal microscope (Infinity 3, VisiTech) on a NikonTi-E microscope stand equipped with a Hamamatsu Flash 4.0V2 sCMOS camera and a Plan Apo 100x/1.45 objective by spanning the whole thickness of the cells using 41 Z slices with a step size of 200 µm. They were deconvolved using Huygens Essential (Scientific Volume Imaging) and Z projected (maximum intensity) and presented. MAPK$^{Spk1}$-GFP signal was found to be increased after induction of sexual differentiation, and accumulation in the nucleus was observed. Yellow arrows indicate a pair of mating cells, which are presented in (B) with a magnified format to reveal cortical accumulation of MAPK$^{Spk1}$-GFP at the shmoo tips. The scale bar represents 10 µm. (B) Individual serial Z-images of the MAPK$^{Spk1}$-GFP signal found in the pair of mating cells indicated by yellow arrows in (A) are presented in a heat-map format by applying the ImageJ Lookup table ”Fire”. Yellow arrows indicate a high-intensity accumulation of MAPK$^{Spk1}$-GFP at the shmoo tips. The scale bar represents 5 µm.

A

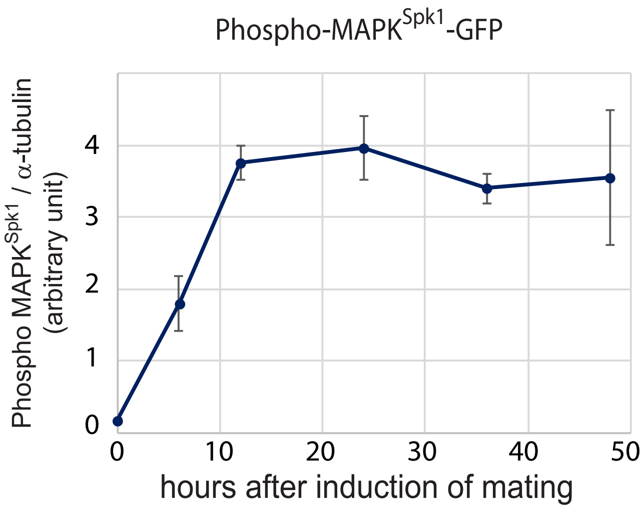
Phospho-MAPK$^{Spk1}$-GFP
Phospho MAPK$^{Spk1}$ / α-tubulin (arbitrary unit)
0
1
2
3
4
0
10
20
30
40
50
hours after induction of mating

B

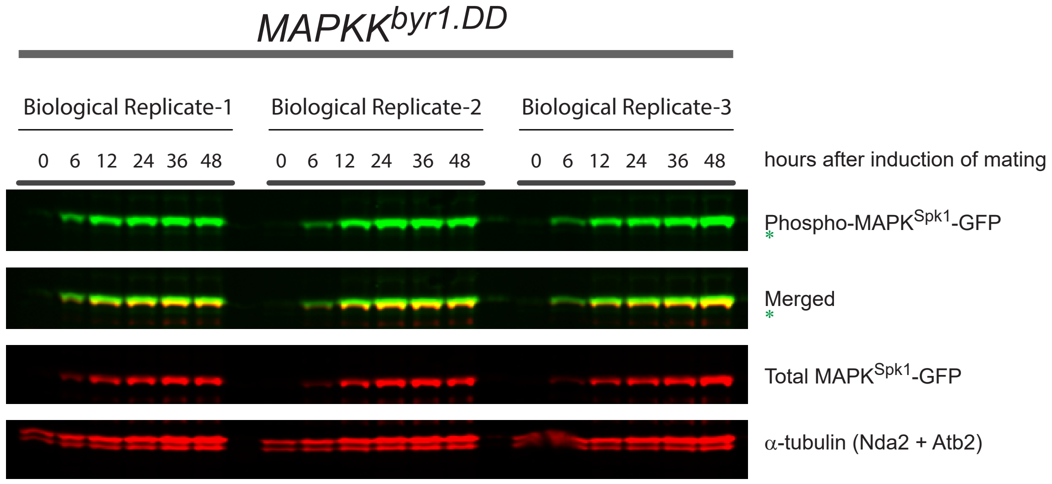
MAPKK$^{byr1.DD}$
Biological Replicate-1
Biological Replicate-2
Biological Replicate-3
0 6 12 24 36 48
0 6 12 24 36 48
0 6 12 24 36 48
hours after induction of mating
Phospho-MAPK$^{Spk1}$-GFP
*
Merged
*
Total MAPK$^{Spk1}$-GFP
α-tubulin (Nda2 + Atb2)

**Supplementary Figure S5.** ***$MAPKK^{byr1.DD}$*** **mutation causes constitutive activation of $MAPK^{Spk1}$.**

The *$MAPKK^{byr1.DD}$* mutant cells (KT3435) were induced for sexual differentiation by the plate mating assay system as described in Supplementary Fig. S1A and the Materials and Methods. Western blotting of **pp**$MAPK^{Spk1}$, total $MAPK^{Spk}$1 and α-tubulin was conducted. The α -tubulin signals were used as an internal control to estimate relative **pp**$MAPK^{Spk1}$ levels. (A) The quantitated results obtained from three biological replicates. The mean value for each time point is presented. Error bars represent SD. (B) Original membrane image of the three biological replicates used for the quantitation.

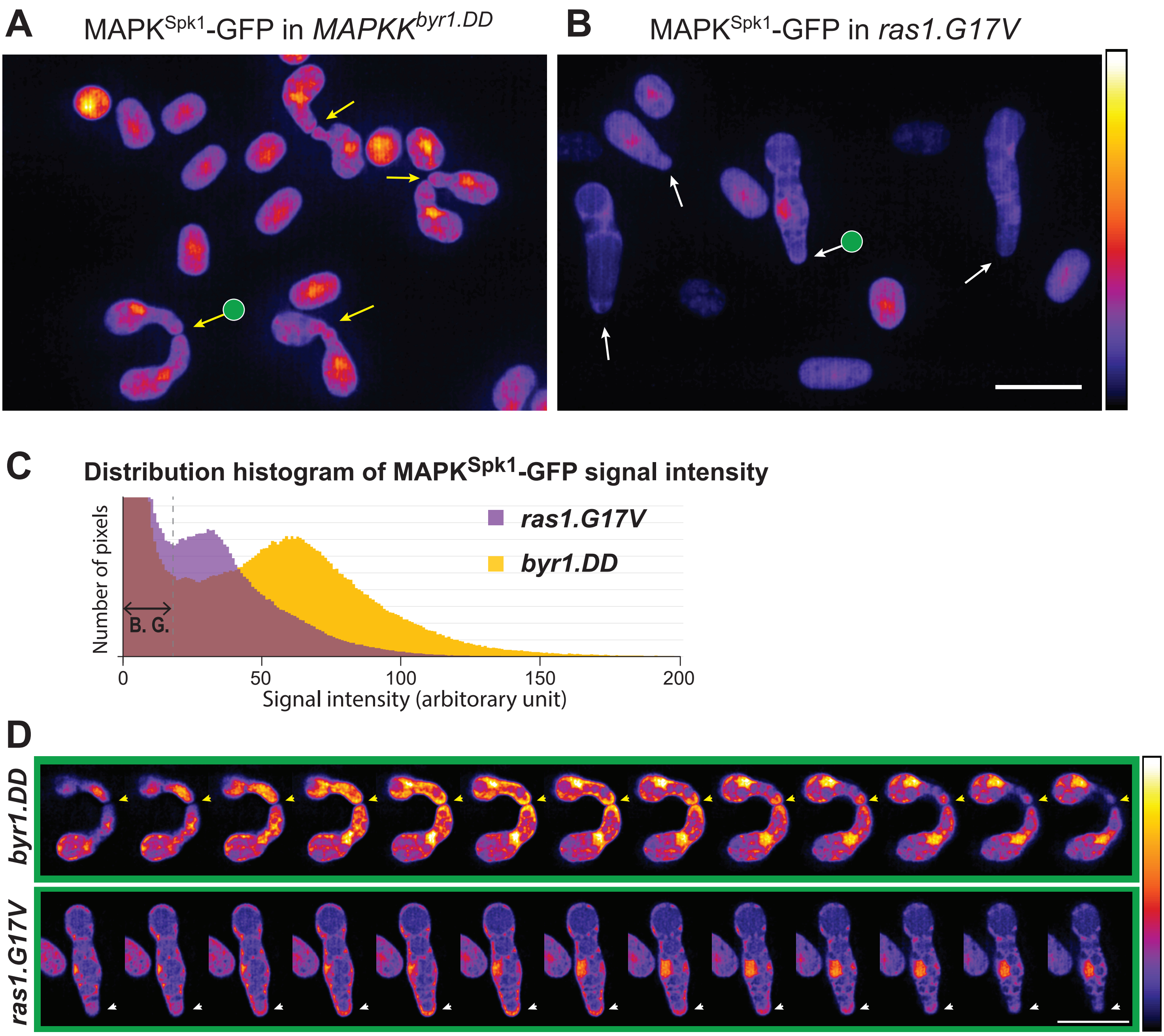
A
MAPK$^{Spk1}$-GFP in *MAPKK$^{byr1.DD}$*
B
MAPK$^{Spk1}$-GFP in *ras1.G17V*
C
Distribution histogram of MAPK$^{Spk1}$-GFP signal intensity
ras1.G17V
byr1.DD
Number of pixels
B. G.
0
50
100
150
200
Signal intensity (arbitorary unit)
D
byr1.DD
ras1.G17V

**Supplementary Figure S6. MAPK$^{Spk1}$-GFP signal intensity is decreased in the *ras1.G17V* mutant at 16 hours after induction of sexual differentiation**

The MAPK$^{Spk1}$-GFP signals were compared 16 hours after induction of sexual differentiation between the cells of *MAPKK$^{byr1.DD}$* (KT3435) (A) and the cells of *ras1.G17V* (KT3084) (B). Images were taken by a 2D-array scanning laser confocal microscope (Infinity 3, VisiTech) on a NikonTi-E microscope stand equipped with a Hamamatsu Flash 4.0V2 sCMOS camera and a Plan Apo 100x/1.45 objective by spanning the whole thickness of the cells using 31 Z slices with a step size of 200 µm. They were deconvolved using Huygens Essential (Scientific Volume Imaging). The serial Z images were Z-projected using the Image J "Sum slices" to compare the total level of MAPK$^{Spk1}$-GFP signal. Signal intensities were indicated in a heatmap format by applying the ImageJ Lookup table (LUT) "Fire" to the Z-projected images. The LUT gradient is indicated on the right-hand side of the panels. Yellow arrows in Panel A indicate the "paired" cells of the *MAPKK$^{byr1.DD}$* mutant. White arrows in Panel B indicate the "elongated" cells of the *ras1.G17V* mutant. Cells indicated by arrows with a green circle are presented as the original serial Z images in Panel (D). The scale bar represents 10µm.

(C) Distribution histograms of pixel signal intensities for *MAPKK$^{byr1.DD}$* (KT3435) (yellow) and *ras1.G17V* (KT3084) (purple). Images that covered 85 cells were subject to ImageJ analysis to measure the MAPK$^{Spk1}$-GFP signal intensity of each pixel of these images. The result is presented as distribution histograms. The peak distribution of pixels of *ras1.G17V* images was found to have a lower signal intensity compared to the one for images of *MAPKK$^{byr1.DD}$*. B.G. indicates the intensities of pixels corresponding to the background area.

(D) Cortical accumulation of MAPK$^{Spk1}$-GFP is weaker in the *ras1.G17V* mutant compared to the *MAPKK$^{byr1.DD}$* mutant at 16 hours after induction of sexual differentiation. MAPK$^{Spk1}$-GFP signals of the serial Z sections of the *MAPKK$^{byr1.DD}$* and the *ras1.G17V* mutant cells indicated with the green circles in panels A and B are presented. Signal intensities are shown in a heatmap format by applying ImageJ LUT "Fire". The LUT gradient is presented on the right-hand side of the panels. Arrowheads indicate the accumulation of MAPK$^{Spk1}$-GFP signals at the cell tips. Signal intensities at the paired *MAPKK$^{byr1.DD}$* cell tips were significantly higher than the one seen at the shmoo tip of the *ras1.G17V* cell.

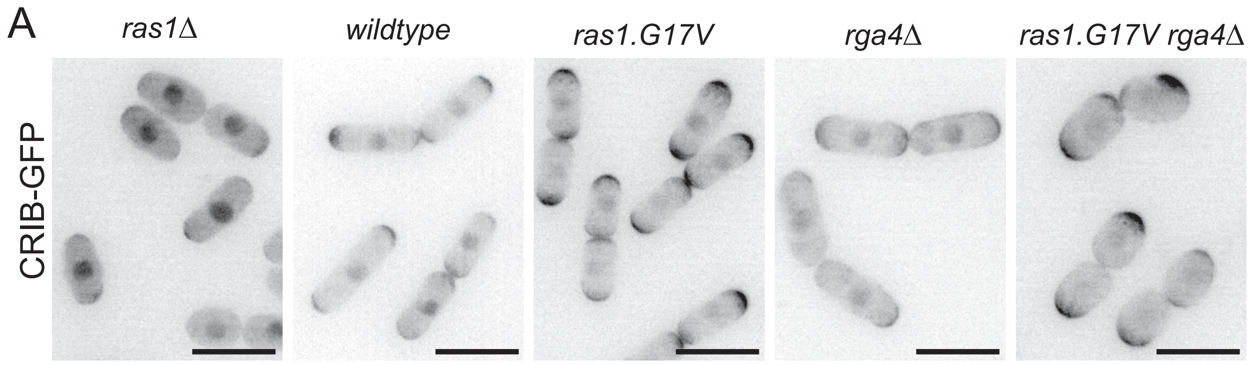
A
ras1Δ
wildtype
ras1.G17V
rga4Δ
ras1.G17V rga4Δ
CRIB-GFP

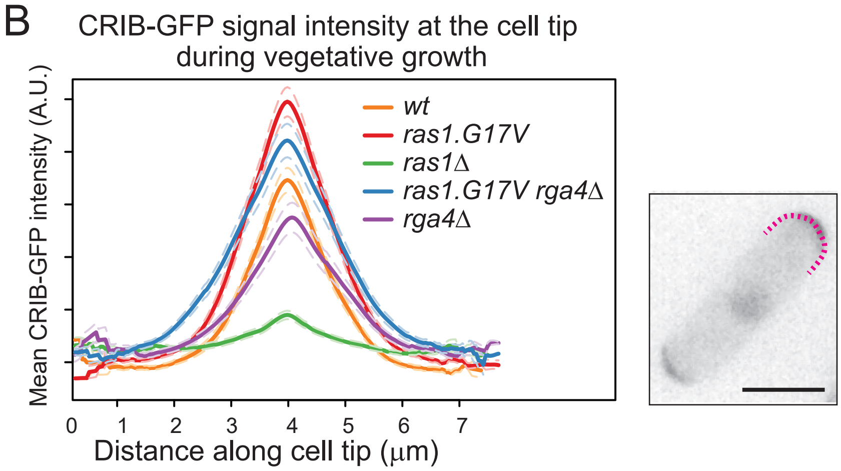
B
CRIB-GFP signal intensity at the cell tip
during vegetative growth
Mean CRIB-GFP intensity (A.U.)
wt
ras1.G17V
ras1Δ
ras1.G17V rga4Δ
rga4Δ
0
1
2
3
4
5
6
7
Distance along cell tip (μm)

**Supplementary Figure S7. Cell morphology and localization of Cdc42$^{GTP}$, indicated by CRIB-GFP signal, during vegetative growth**

(A) Representative CRIB-GFP signal images of vegetatively growing cells of wildtype (KT5077), *ras1Δ* (5107), *ras1.G17V* (KT5082), *rga4Δ* (5551) and *rga4Δ ras1.G17V* (KT5554) are presented. The scale bar is 10 µm. (B) Quantitated CRIB-GFP signals on the cell cortex of cells presented in (A). The intensity of the GFP signal on the cell cortex was measured along one of the cell tips with a stronger GFP signal as indicated as a magenta dotted line in the example image on the right (Scale bar: 10 µm) as stated in the Materials and Methods. 40 cells without septum were measured for each strain, and the average curve from all aligned traces per strain was calculated and displayed with respective standard error of the mean curves (dashed lines) as described in Materials and Methods.

Supplementary. Fig. S8

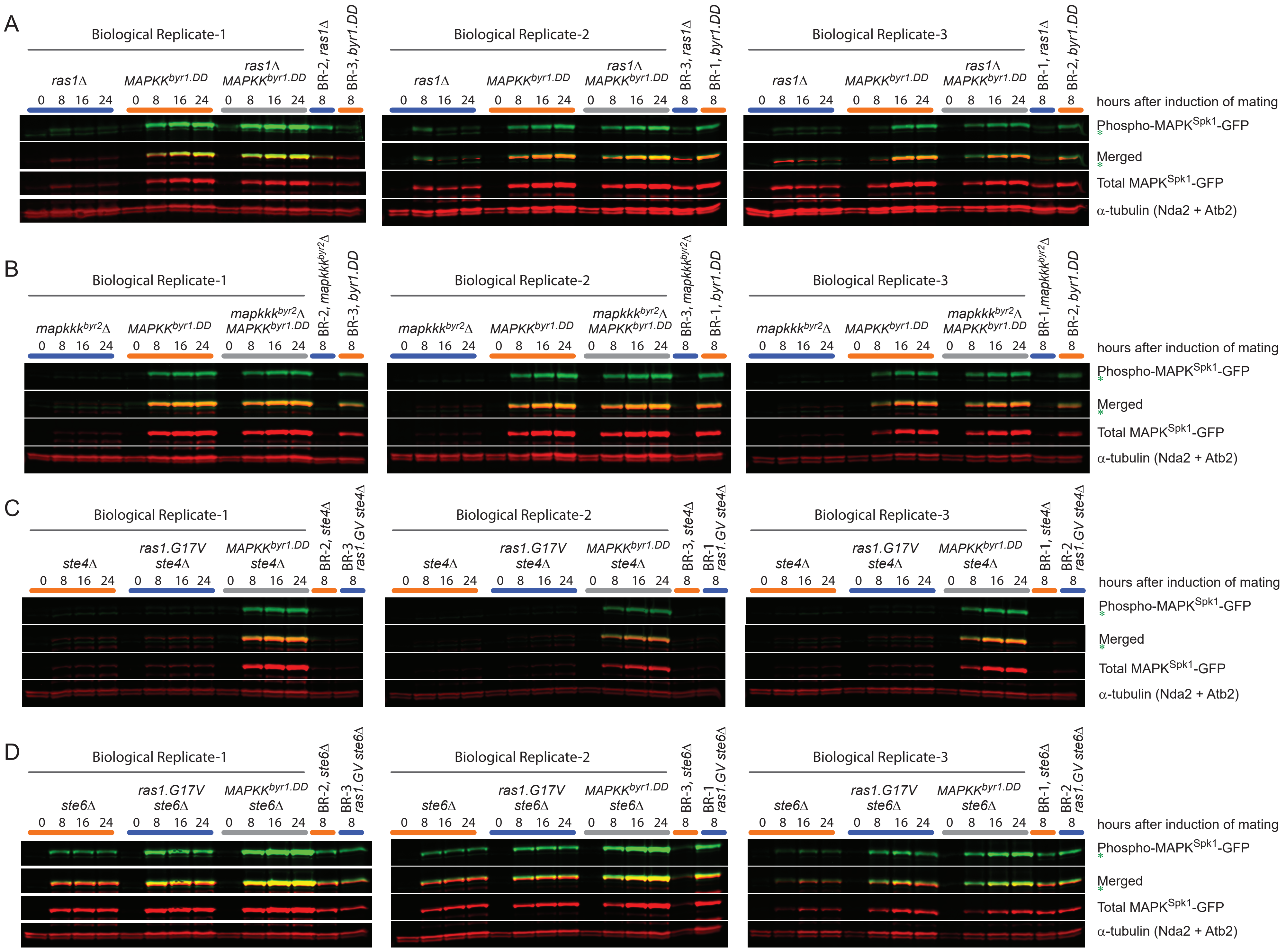

**Supplementary Figure S8. Original Western blotting images used to quantify the ppMAPK$^{Spk1}$-GFP levels during the mating process in Fig. 3 and 6**

(A) Original Western blotting membranes used to quantify the **pp**MAPK$^{Spk1}$-GFP to generate the graph presented in Figure 3D. Strains used were; *ras1Δ* (KT4323), *MAPKK$^{byr1.DD}$* (KT3435) and *ras1Δ MAPKK$^{byr1.DD}$* (KT4359) (B) Original Western blotting membranes used to quantify the **pp**MAPK$^{Spk1}$-GFP to generate the graph presented in Figure 3E. Strains used were; in *mapkkk$^{byr2}$Δ* (KT3763), *MAPKK$^{byr1.DD}$* (KT3435) and *mapkkk$^{byr2}$Δ MAPKK$^{byr1.DD}$* (KT4010). (C) Original Western blotting membranes used to quantify the **pp**MAPK$^{Spk1}$-GFP to generate the graph in Figure 6A. Strains used were; *ste4Δ* (KT4376), *ste4Δ ras1.G17V* (KT5143) and *ste4Δ MAPKK$^{byr1.DD}$* (KT5136). (D) Original Western blotting membranes used to quantify the **pp**MAPK$^{Spk1}$-GFP to generate the graph in Figure 6C. Strains used were; *ste6Δ* (KT4333), *ste6Δ ras1.G17V* (KT4998) and *ste6Δ MAPKK$^{byr1.DD}$* (KT5139). Indicated mutant cells were induced for mating by the plate mating assay system as described in the Materials and Methods. Western blotting of ppMAPK$^{Spk1}$, total MAPK$^{Spk1}$ and α-tubulin was conducted in the same way as Supplementary Figure S5. The green asterisks indicate the same background band recognized by the anti-phospho-ERK antibody (#4370 Cell Signaling Technology. Used at 1:2000 dilution) seen in Supplementary Figure S1. α-tubulin was used as a loading control, and quantitation was carried out using the Image Studio ver2.1 software (Licor Odyssey CLx Scanner).

Supplementary Fig. S9

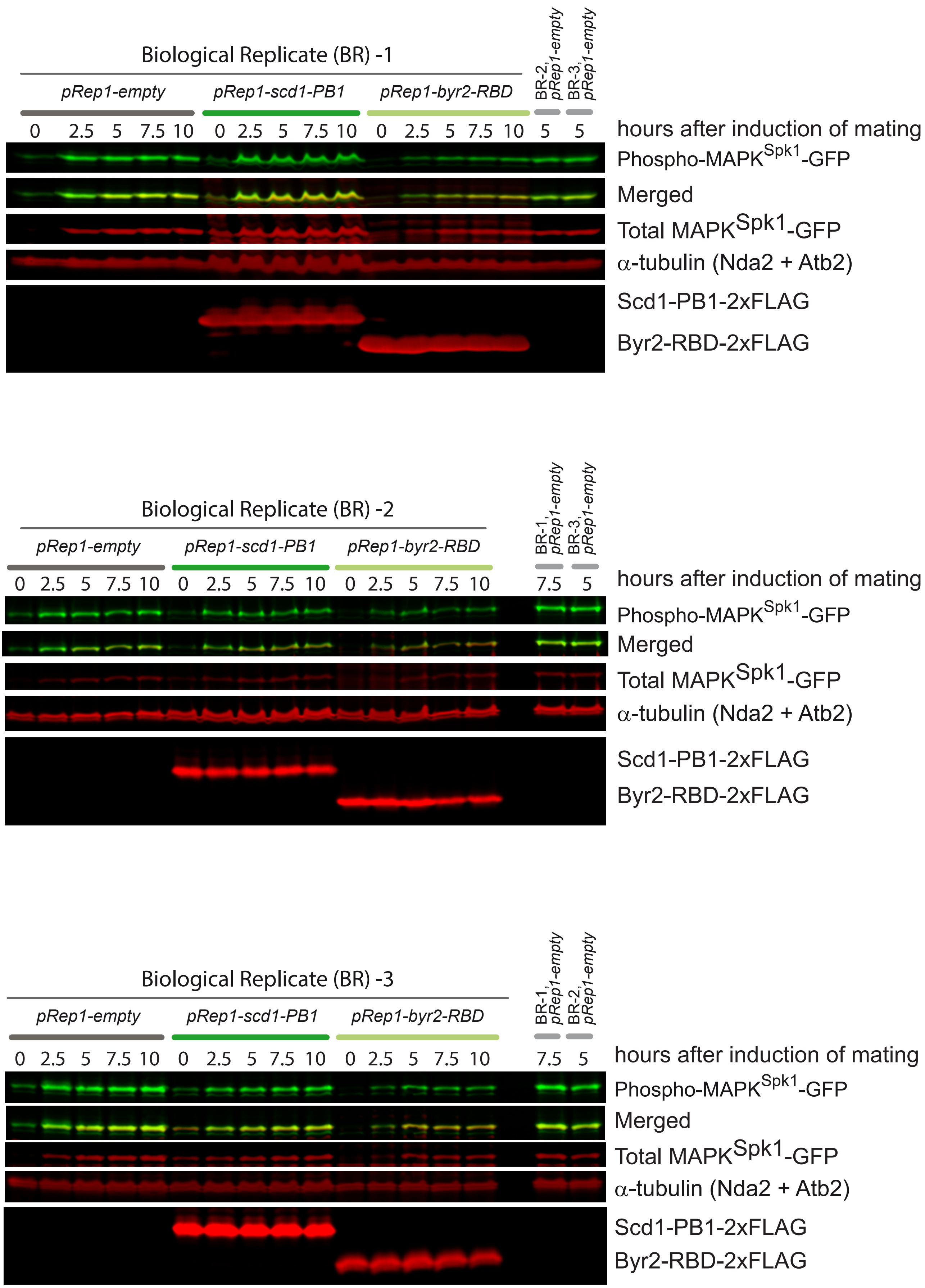

**Supplementary Figure S9. Original Western blotting membranes used in Fig. 5A.**

Original Western blotting membranes used to quantify ppMAPK$^{Spk1}$-GFP to generate the graph in Figure 5A. Cells harboring *ras1.G17V* and MAPK$^{Spk1}$-GFP-2xFLAG (KT5940) were transformed with either pRep1 empty vector, pRep1-*scd1*(760-872)-2xFLAG or pRep1-*byr2*(65-180)-2xFLAG and cultured in MM+N without thiamine for 24 hours. Cells were induced for sexual differentiation by the plate mating assay system, and the levels of ppMAPK$^{Spk1}$-GFP, total MAPK$^{Spk1}$ and $\alpha$-tubulin (loading control) were examined by Western blotting. Three biological replicates are shown. The quantitation was carried out using the Image Studio ver2.1 software (Licor Odyssey CLx Scanner). The expressions of Scd1(760-872)-2xFLAG and Byr2(65-180)-2xFLAG were confirmed by anti-FLAG antibody (Sigma F1804, 1 µg/ul, 1/2000 dilution).

Supplementary Fig. S10-A

A

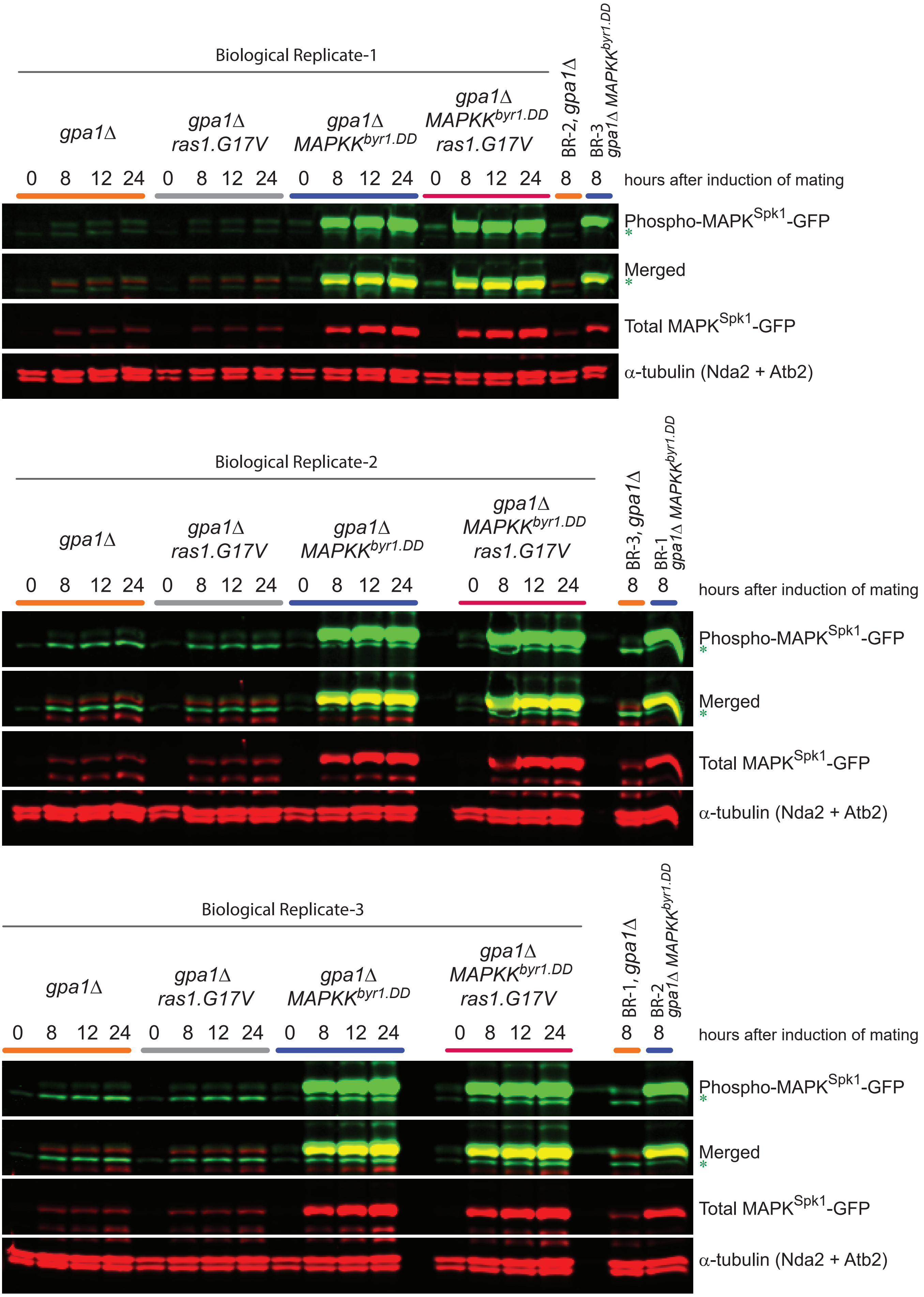

Supplementary Fig. S10-B

B

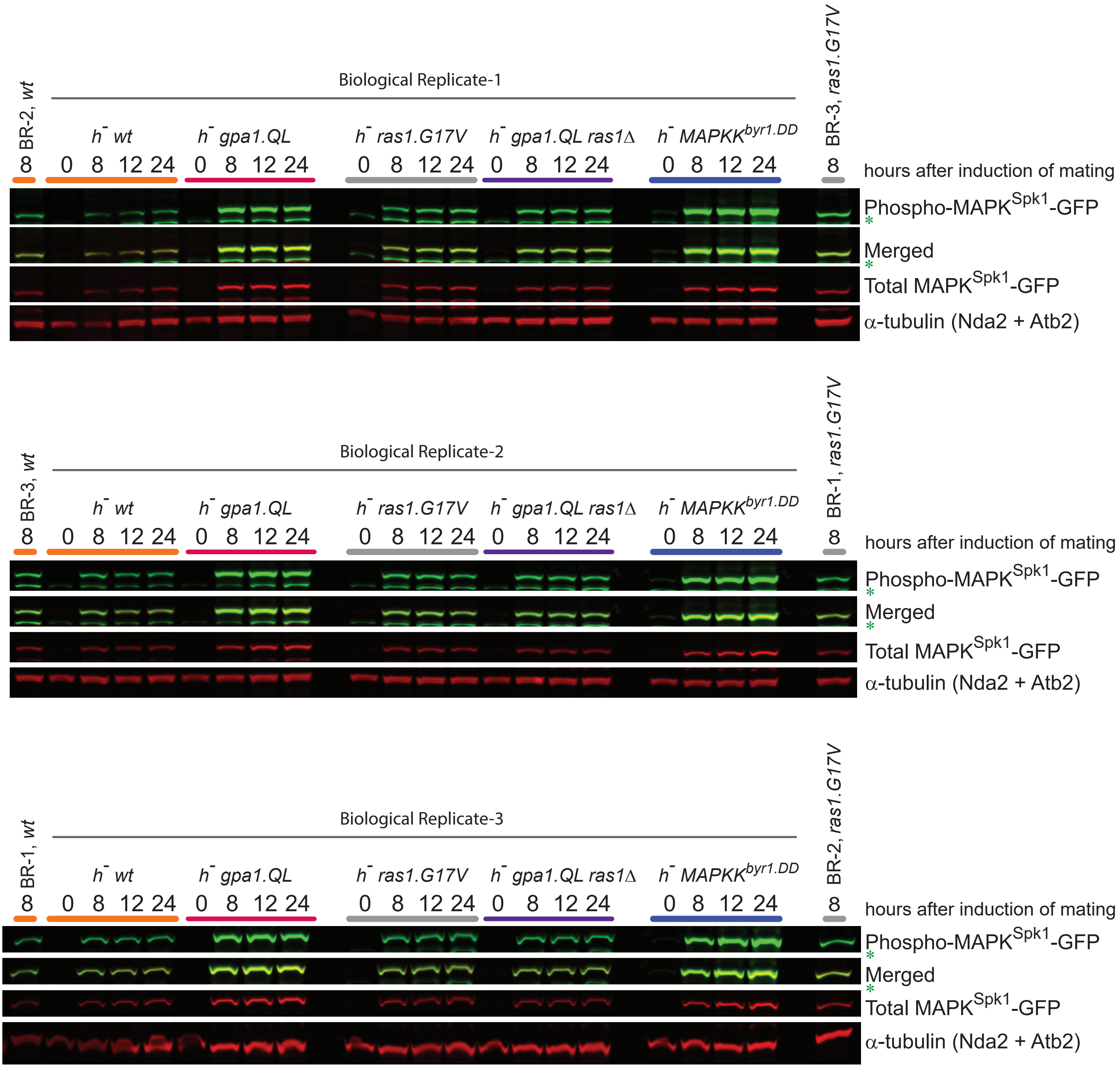

**Supplementary Figure S10. Original Western blotting membranes used in Fig. 7.**

(A) Original Western blotting membranes used to quantitate ppMAPK$^{Spk1}$-GFP to generate the graph in Figure 7A. Strains used were; *gpa1Δ* (KT4335), *gpa1Δ ras1.G17V* (KT5023), *gpa1Δ MAPKK$^{byr1.DD}$* (KT4353) and *gpa1Δ ras1.val17 MAPKK$^{byr1.DD}$* (KT5035). (B) Original Western blotting membranes used to quantitate ppMAPK$^{Spk1}$-GFP to generate the graph in Figure 7C. Strains used were; *$h^-$* WT (KT4190), *$h^-$ gpa1.QL* (KT5059), *$h^-$ ras1.G17V* (KT4233), *$h^-$ gpa1.QL ras1Δ* (KT5070), $h^-$ *MAPKK$^{byr1.DD}$* (KT4194). The green asterisk indicates a background band recognized by anti-phospho-ERK antibody (#4370 Cell Signaling technology. Used at 1:2000 dilution). α-tubulin was used as a loading control and quantitation was carried out using the Image Studio ver2.1 software (Licor Odyssey CLx Scanner).

Supplementary Fig. S11

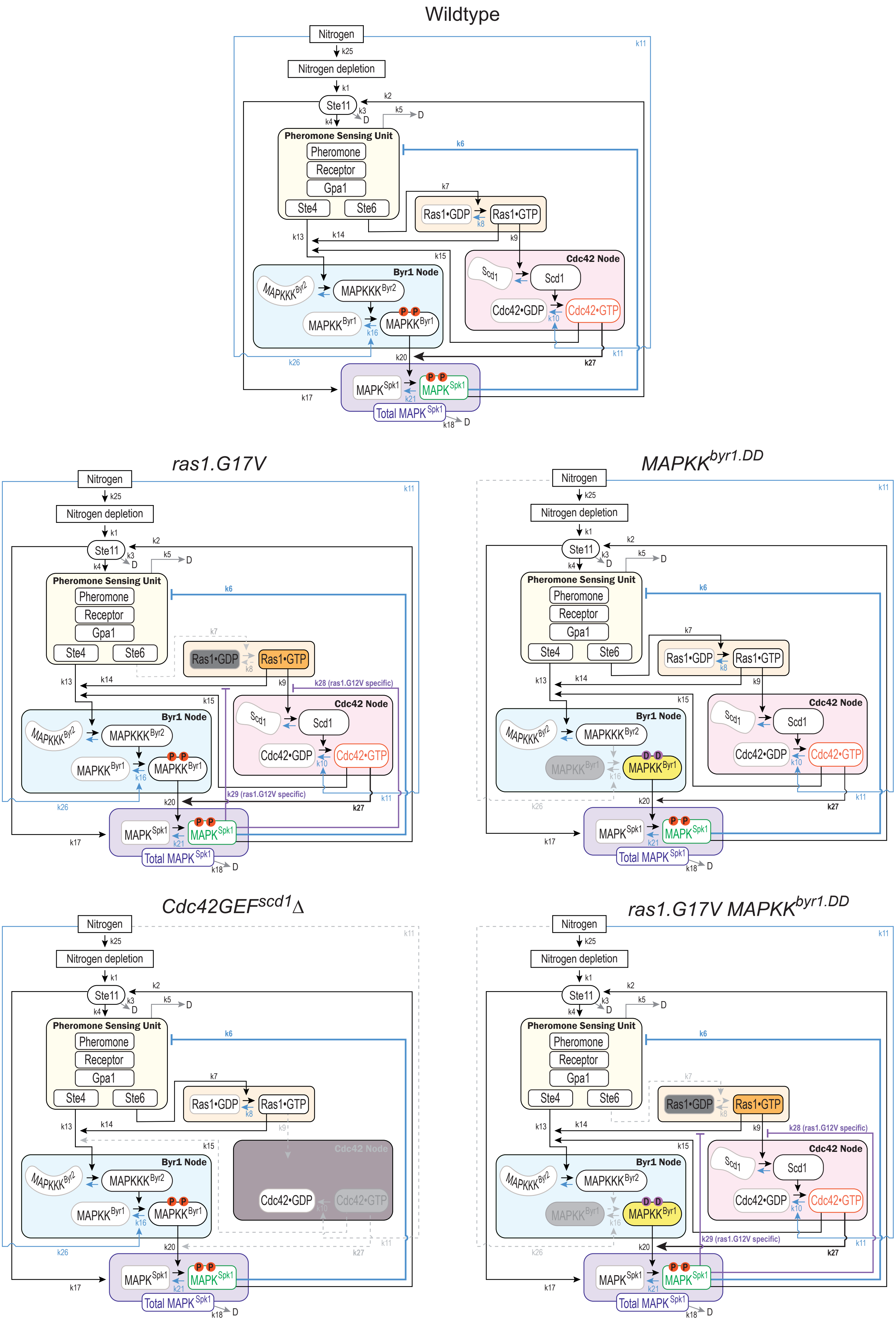

**Supplementary Figure S11. Components and framework of the mathematical model**

Components and framework of the mathematical model (Model B in this instance) of wildtype and signalling mutants, *ras1.G17V*, $MAPKK^{byr1.DD}$, *ras1.G17V* $MAPKK^{byr1.DD}$ and *scd1Δ*. Changes corresponding to each mutant are indicated as follows: Grey: removed components or interactions, orange: the activation process does not require the upstream elements. For the exact implementation of the mutants, see Materials and Method. The measured components, total $MAPK^{Spk1}$, **pp**$MAPK^{Spk1}$ and $Cdc42^{GTP}$, are indicated in purple, green and red, respectively.

Fig. S12

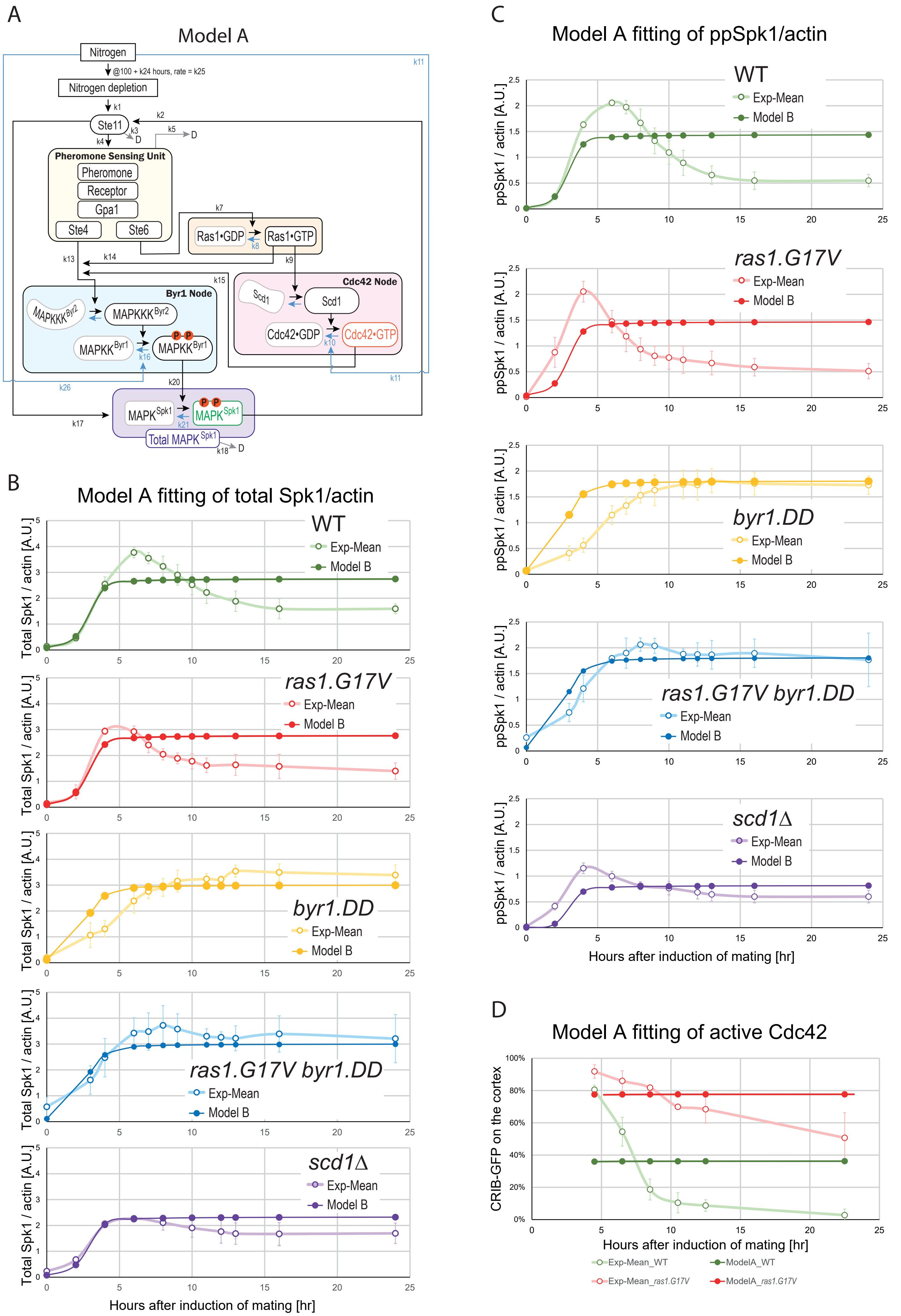
A
Model A
Nitrogen
@100 + k24 hours, rate = k25
Nitrogen depletion
Ste11
Pheromone Sensing Unit
Pheromone
Receptor
Gpa1
Ste4
Ste6
Ras1•GDP
Ras1•GTP
Byr1 Node
MAPKKK Byr2
MAPKK Byr1
Cdc42 Node
Scd1
Cdc42•GDP
Cdc42•GTP
MAPK Spk1
Total MAPK Spk1
k1
k2
k3
k4
k5
k7
k8
k9
k10
k11
k13
k14
k15
k16
k17
k18
k20
k21
k26
D
B
Model A fitting of total Spk1/actin
Total Spk1 / actin [A.U.]
WT
ras1.G17V
byr1.DD
ras1.G17V byr1.DD
scd1Δ
Exp-Mean
Model B
Hours after induction of mating [hr]
C
Model A fitting of ppSpk1/actin
ppSpk1 / actin [A.U.]
WT
ras1.G17V
byr1.DD
ras1.G17V byr1.DD
scd1Δ
Exp-Mean
Model B
Hours after induction of mating [hr]
D
Model A fitting of active Cdc42
CRIB-GFP on the cortex
Hours after induction of mating [hr]
Exp-Mean_WT
ModelA_WT
Exp-Mean_ras1.G17V
ModelA_ras1.G17V

**Supplementary Figure S12. Models A and its fittings for the total MAPK$^{Spk1}$, ppMAPK$^{Spk1}$, and active Cdc42 levels**

(A) Model A components and framework. For the detailed implementation of the mutants, see Supplementary Figure S11 and Materi als and Methods. The measured components, total MAPK$^{Spk1}$, **pp**MAPK$^{Spk1}$ and Cdc42$^{GTP}$, are shown in purple, green and red, respectively. (B) and (C) Model A fittings for the total MAPK$^{Spk1}$ (B) and **pp**MAPK$^{Spk1}$ (C) levels in wildtype, *ras1.G17V*, *MAPKK$^{byr1.DD}$*, *ras1.G17V MAPKK$^{byr1.DD}$* and *Cdc42GEF$^{scd1}$Δ* mutants. Mean values (open circles) and SD values (error bars) of the experimental results are shown in pale colours, and the model-fitted values are shown as filled circles in darker colours. (D) Model A fitting of the active Cdc42 levels in wildtype and *ras1.G17V* mutants. Mean values (open circles) and SD values (error bars) of the experimental results are shown in pale colours, and the model-fitted values are shown in filled circles in darker colours.

Fig. S13

### Model B fitting of total Spk1/actin

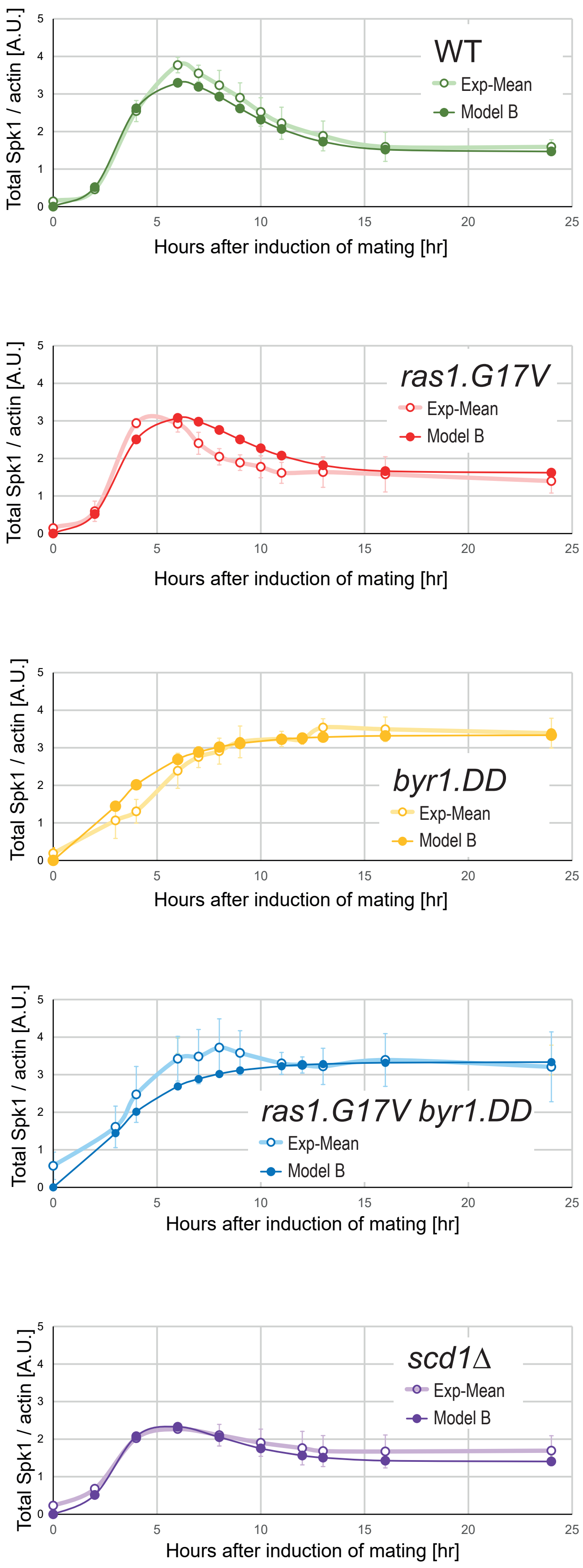

**Supplementary Figure S13. Models B fittings for the total MAPK$^{Spk1}$ levels**

Model B fittings for the total MAPK$^{Spk1}$ levels in wildtype, *ras1.G17V*, *MAPKK$^{byr1.DD}$*, *ras1.G17V MAPKK$^{byr1.DD}$* and *Cdc42GEF$^{scd1}$Δ* mutants. Mean values (open circles) and SD values (error bars) of the experimental results are shown in pale colours, and the model-fitted values are shown as filled circles in darker colours.

Fig. S14

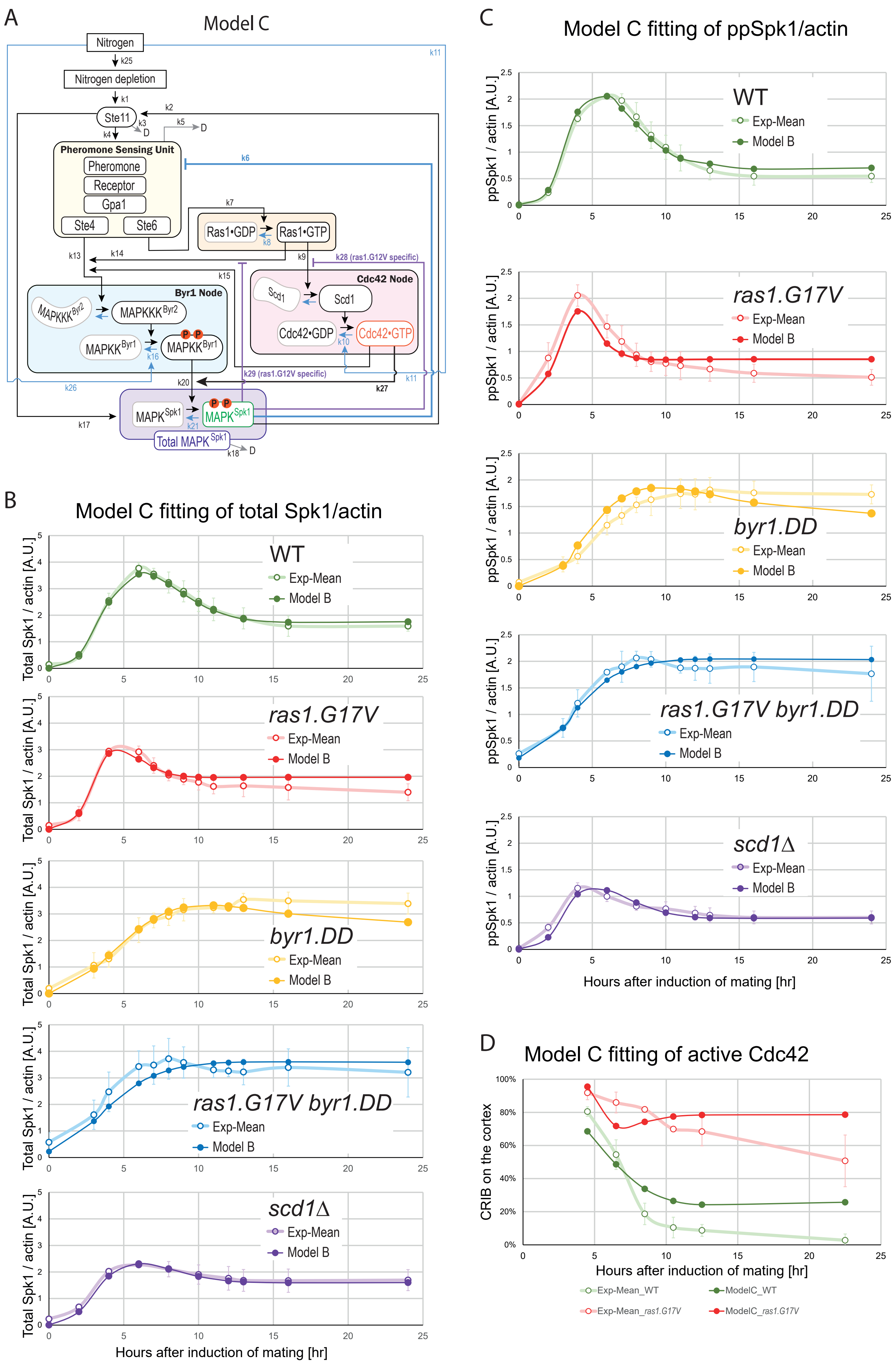
A
Model C
Nitrogen
k25
Nitrogen depletion
k1
Ste11
k2
k3
D
k4
k5
D
Pheromone Sensing Unit
Pheromone
Receptor
Gpa1
Ste4
Ste6
k6
k7
Ras1•GDP
Ras1•GTP
k8
k9
k28 (ras1.G12V specific)
k13
k14
k15
Byr1 Node
MAPKKK Byr2
MAPKKK Byr2
MAPKK Byr1
MAPKK Byr1
k16
Cdc42 Node
Scd1
Scd1
Cdc42•GDP
Cdc42•GTP
k10
k11
k26
k20
k29 (ras1.G12V specific)
k27
k17
MAPK Spk1
MAPK Spk1
k21
Total MAPK Spk1
k18
D
B
Model C fitting of total Spk1/actin
Total Spk1 / actin [A.U.]
WT
Exp-Mean
Model B
ras1.G17V
byr1.DD
ras1.G17V byr1.DD
scd1Δ
Hours after induction of mating [hr]
C
Model C fitting of ppSpk1/actin
ppSpk1 / actin [A.U.]
WT
ras1.G17V
byr1.DD
ras1.G17V byr1.DD
scd1Δ
Exp-Mean
Model B
Hours after induction of mating [hr]
D
Model C fitting of active Cdc42
CRIB on the cortex
Hours after induction of mating [hr]
Exp-Mean_WT
ModelC_WT
Exp-Mean_ras1.G17V
ModelC_ras1.G17V

**Supplementary Figure S14. Models C and its fittings for the total MAPK$^{Spk1}$, ppMAPK$^{Spk1}$, and active Cdc42 levels**

(A) Model C components and framework. For the detailed implementation of the mutants, see Supplementary Figure S11 and Materi als and Methods. The measured components, total MAPK$^{Spk1}$, **pp**MAPK$^{Spk1}$ and Cdc42$^{GTP}$, are shown in purple, green and red, respectively. (B) and (C) Model C fittings for the total MAPK$^{Spk1}$ (B) and **pp**MAPK$^{Spk1}$ (C) levels in wildtype, *ras1.G17V*, *MAPKK$^{byr1.DD}$*, *ras1.G17V MAPKK$^{byr1.DD}$* and *Cdc42GEF$^{scd1}$Δ* mutants. Mean values (open circles) and SD values (error bars) of the experimental results are shown in pale colours, and the model-fitted values are shown as filled circles in darker colours. (D) Model C fitting of the active Cdc42 levels in wildtype and *ras1.G17V* mutants. Mean values (open circles) and SD values (error bars) of the experimental results are shown in pale colours, and the model-fitted values are shown in filled circles in darker colours.

Fig. S15

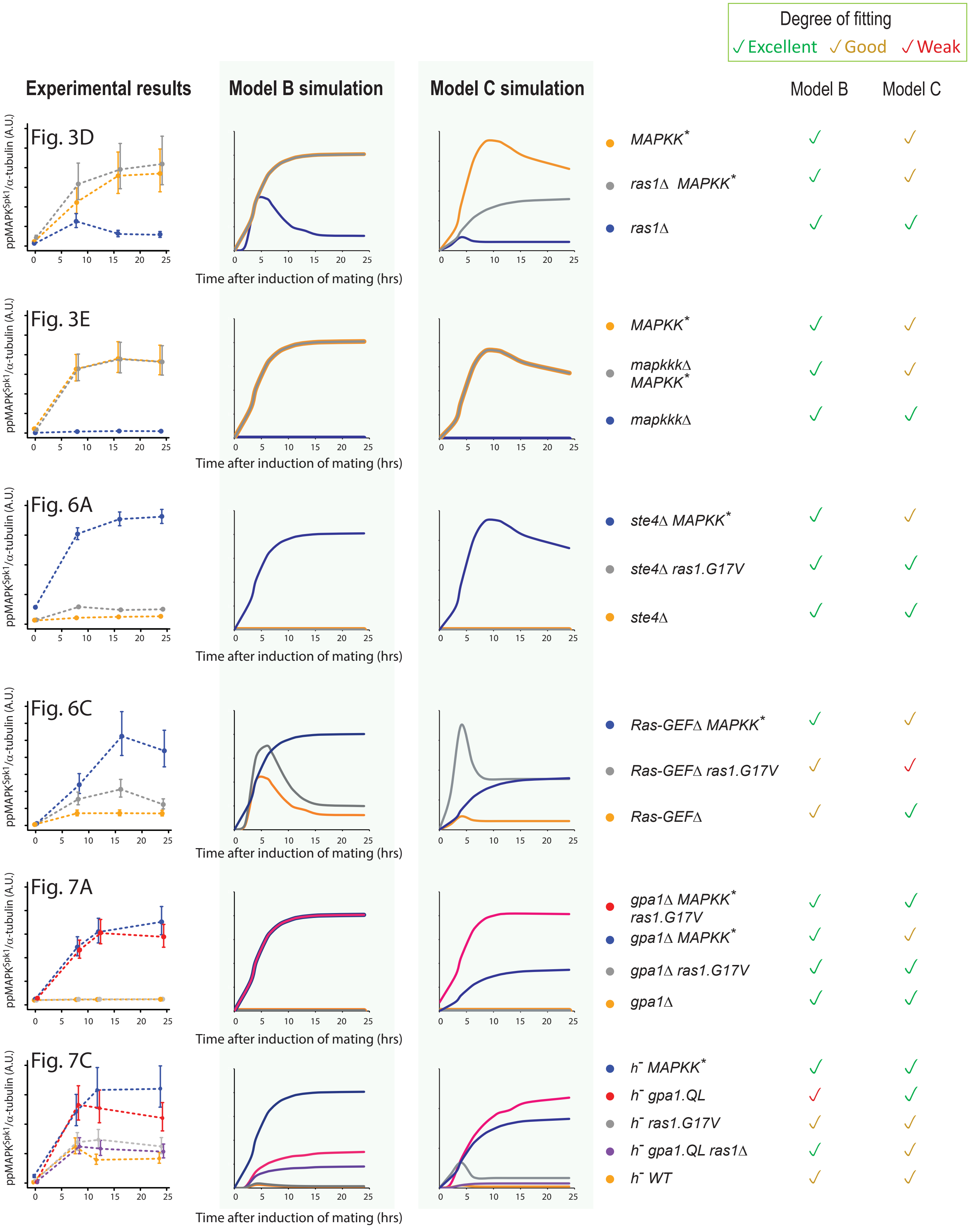
Degree of fitting
✓Excellent ✓Good ✓Weak
Experimental results
Model B simulation
Model C simulation
Model B
Model C
Fig. 3D
MAPKK*
ras1Δ MAPKK*
ras1Δ
Fig. 3E
MAPKK*
mapkkkΔ MAPKK*
mapkkkΔ
Fig. 6A
ste4Δ MAPKK*
ste4Δ ras1.G17V
ste4Δ
Fig. 6C
Ras-GEFΔ MAPKK*
Ras-GEFΔ ras1.G17V
Ras-GEFΔ
Fig. 7A
gpa1Δ MAPKK* ras1.G17V
gpa1Δ MAPKK*
gpa1Δ ras1.G17V
gpa1Δ
Fig. 7C
h- MAPKK*
h- gpa1.QL
h- ras1.G17V
h- gpa1.QL ras1Δ
h- WT
ppMAPK^Spk1/α-tubulin (A.U.)
Time after induction of mating (hrs)
0 5 10 15 20 25

**Supplementary Figure S15. Prediction ability of Models B and C**

Model B and C were used to predict the experimental results shown in Fig. 3D, Fig. 3E, Fig. 6A, Fig. 6C, Fig. 7A and Fig. 7C, involving 20 fission yeast strains. First, the **pp**Spk1 of *MAPKK*$^{byr1\Delta}$ (Supplementary Fig. S1F) was predicted by fixing the amount of $MAPKK^{Byr1}$ to zero in Model B/C. The models predicted a minimum ppSpk1 level in *MAPKK*$^{byr1\Delta}$, consistent with the experiment (data not shown).

**Fig. 3D prediction involving *MAPKK*$^{byr1.DD}$, *ras1Δ*, and *ras1Δ MAPKK*$^{byr1.DD}$**: the amount of Ras1 was fixed to zero for *ras1Δ*. Model B predicted the **pp**$MAPK^{Spk1}$ relative levels in these three strains both qualitatively and quantitatively well. Model C also made a qualitatively good prediction, but it predicted the *MAPKK*$^{byr1.DD}$ single mutation to generate a higher level of **pp**$MAPK^{Spk1}$ compared to the *ras1Δ MAPKK*$^{byr1.DD}$ double mutant, although the experimental result did not show a significant difference between these two strains. This discrepancy was because that Model C was constructed on an assumption that Ras1 positively contributes to the **pp**Spk1 production independent of Byr1 at the rate *k27* (to cause an earlier increase of the **pp**Spk1 in the *ras1.G17V MAPKK*$^{byr1.DD}$ double mutant compared to the *MAPKK*$^{byr1.DD}$ single mutant as suggested in Supplementary Fig. S3D). The Model C result suggests that the predicted Ras1 contribution to the $MAPK^{Spk1}$ activation is more complex than we considered in the proposed models.

**Fig. 3E prediction involving *MAPKK*$^{byr1.DD}$, *mapkkk*$^{byr2}$*Δ MAPKK*$^{byr1.DD}$, and *mapkkk*$^{byr2}$*Δ***: the activation of $MAPKK^{Byr1}$ was set to zero for the strains carrying the *mapkkk*$^{byr2}$*Δ* allele. Both Model B and C predicted the experimental result qualitatively and quantitively.

**Fig. 6A prediction involving *ste4Δ MAPKK*$^{byr1.DD}$, *ste4Δ ras1*$^{G17V}$, and *ste4Δ***: the activation of $MAPKK^{Byr1}$ was set to zero for the strains carrying the *ste4Δ* allele. Therefore, the *ste4Δ* model should behave identically to the *mapkkk*$^{byr2}$*Δ* model examined above. Both Model B and C predicted the experimental result qualitatively and quantitively.

**Fig. 6C prediction involving *ste6Δ MAPKK*$^{byr1.DD}$, *ste6Δ ras1*$^{G17V}$, and *ste6Δ***: the activation of Ras1 was set to zero for the strains carrying the *ste6Δ* allele. Both Model B and C predicted a higher **pp**Spk1 for *ste6Δ MAPKK*$^{byr1.DD}$. Model C also predicted a lower ppSpk1 throughout the time course, which agrees with the experimental result. Model B predicts a lower **pp**Spk1 for *ste6Δ ras1*$^{G17V}$ and *ste6Δ* for the later time points (10-25 hours after induction of mating), which were consistent with the experiment. However, Model B's prediction of substantial **pp**Spk1 increase in the earlier time points in the *ste6Δ* $ras1^{G17V}$ and *ste6Δ* mutants was not seen in the experimental result. Model C predicts a high **pp**Spk1 for the *ste6Δ ras1.G17V*, which is not the case in the experimental result. The result indicates that Ste6 is involved in activating **pp**Spk1 downstream of Ras1, which is not included in our current model.

**Fig. 7A prediction involving *gpa1Δ ras1.G17V* $MAPKK^{byr1.DD}$ triple mutant, *gpa1Δ ras1.G17V* double mutant, *gpa1Δ* $MAPKK^{byr1.DD}$ double mutant, and *gpa1Δ* single mutant:** the activation of both Ras1 and Byr1 was set to zero for the strains carrying the *gpa1Δ* allele. Model B predicted the experimental result very well. Model C also reproduced the experimental result of the *gpa1Δ ras1.G17V* double mutant and the *gpa1Δ* single mutant, predicting the minimum level of the **pp**Spk1. Meanwhile, for the *gpa1Δ ras1.G17V* $MAPKK^{byr1.DD}$ triple mutant and the *gpa1Δ* $MAPKK^{byr1.DD}$ double mutant, Model C predicted a higher **pp**Spk1 level for the *gpa1Δ ras1.G17V* $MAPKK^{byr1.DD}$ triple mutant than the *gpa1Δ* $MAPKK^{byr1.DD}$ double mutant. This is because Model C postulates that Ras1 can boost the **pp**Spk1 level independent of $MAPKK^{Byr1}$ at a rate constant k27, as discussed above. This Ras1 contribution should be smaller than we assumed in Model C.

**Fig. 7C prediction involving $h^-$ $MAPKK^{byr1.DD}$, $h^-$ *gpa1.QL*, $h^-$ *ras1.G17V*, $h^-$ *gpa1.QL ras1Δ* and $h^-$ wildtype:** in order to reflect the lack of pheromone signalling in these $h^-$ strains, the activation levels of Ras1 and Byr1 were reduced to 1/100 of the $h^{90}$ wild type values, and the amount of active Ste4/Ste6 (i.e. pheromone sensing unit) was set to the value of 0.2. Both Model B and C recapitulated the experimental result qualitatively by predicting relatively higher **pp**Spk1 levels for $h^-$ $MAPKK^{byr1.DD}$ and $h^-$ *gpa1.QL* strains, whereas the rest of the strains are predicted to produce lower, but non-zero, **pp**Spk1 levels. Model C predicted the $h^-$ *ras1.G17V* mutant to produce an initial **pp**Spk1 increase, which is higher than the experiment, indicating that the activity of Ras1.G17V was estimated too high.

**Table S1. Model parameters and opimized values**

| | | Model A | Model B | Model C |
|---|---|---|---|---|
| ppSpk1: constant to normalize the efficiency of the Western blot (compared to that of tSpk1) | k12 | 0.87 | 1.54 | 2.02 |
| aCdc42: constant to convert CRIB-GFP[%] to the Western blot signal (of tSpk1) | k19 | 0.89 | 14.59 | 9.83 |
| time: nitrogen starvation become effective at this time [x0.5 hr] after the removal of nitrogen in the experiments | k24 | 3.88 | 3.36 | 2.75 |
| time: the steapness parameter of the nitrogen starvation | k25 | 0.65 | 0.0012 | 0.0048 |
| aRas1: [aRas1] at the ras1.G17V mutant | k22 | 9.58 | 0.02 | 0.25 |
| ppByr1: [ppByr1] at the byr1.DD mutant | k23 | 1.40 | 0.05 | 0.16 |
| Ste11: activation constant due to nitrogen starvation | k1 | 1.97 | 4.09 | 5.79 |
| Ste11: activation constant due to ppSpk1 | k2 | 0.66 | 6.78 | 20.53 |
| Ste11: basal inactivation constant (due to degradation) | k3 | 2.93 | 8.70 | 24.41 |
| Pherome Sensing Unit (PSU) including Ste4/6: activation constant due to Ste11 | k4 | 7.15 | 14.80 | 28.59 |
| PSU including Ste4/6: basal inactivation constant | k5 | 17.28 | 17.55 | 7.22 |
| aRas1: activation constant due to PSU including Ste4/6 | k7 | 5.81 | 25.22 | 2.83 |
| aRas1: basal inactivation constant | k8 | 0.11 | 0.24 | 0.25 |
| aCdc42: activation constant due to aRas1 | k9 | 3.31 | 14.32 | 0.36 |
| aCdc42: basal inactivation constant | k10 | 4.60 | 6.04 | 0.26 |
| aCdc42: inactivation constant due to nitrogen | k11 | 0.26 | 1.02 | 0.01 |
| ppByr1: activation constant due to PSU including Ste4/6 | k13 | 30.19 | 18.06 | 5.47 |
| ppByr1: activation constant due to aRas1 | k14 | 0.13 | 0.21 | 322.85 |
| ppByr1: activation constant due to aCdc42 | k15 | 48.10 | 21.08 | 0.12 |
| ppByr1: basal inactivation constant | k16 | 16.81 | 0.10 | 0.54 |
| ppByr1: inactivation constant due to nitrogen | k26 | 0.10 | 0.60 | 0.48 |
| tSpk1: activation constant due to Ste11 | k17 | 4.10 | 2.76 | 2.18 |
| tSpk1: basal inactivation constant | k18 | 1.55 | 1.23 | 0.66 |
| ppSpk1: activation constant due to ppByr1 | k20 | 14.26 | 12.50 | 1.20 |
| ppSpk1: basal inactivation constant | k21 | 8.88 | 0.92 | 0.69 |
| PSU including Ste4/6: inactivation constant due to ppSpk1 | k6 | | 36614.68 | 3447.85 |
| ppSpk1: activation constant due to aCdc42 | k27 | | | 11.95 |
| aCdc42: inactivation constant due to ppSpk1 in the ras1.G17V mutant | k28 | | | 3.62 |
| ppByr1: inactivation constant due to ppSpk1 in the ras1.G17V mutant | k29 | | | 18.46 |
| maximum log-likelihood | lmax | -610.75 | -141.84 | -109.07 |
| the number of parameters | k | 25 | 26 | 29 |
| AIC (Akaike Information Criterion) | AIC | 1271 | 336 | 276 |

**Supplementary Table S2 : Strains used in this study**

**Figure 1**

**C, D**

**KT3082** *$h^{90}$ ade6.M216 leu1.32 spk1-GFP-2xFLAG::$Kan^R$*

**E, F**

**KT3435** *$h^{90}$ ade6-M216 leu1.32 byr1.DD::$Hyg^R$ spk1-GFP-2xFLAG::$Kan^R$*

**G, H**

**KT3084** *$h^{90}$ ade6.M210 leu1.32 ras1.G17V::$LEU2^+$ spk1-GFP-2xFLAG::$Kan^R$*

**Figure 2**

**A, B**

**KT3439** *$h^{90}$ ade6.M210 leu1.32 ras1.G17V::$LEU2^+$ byr1.DD::$Hyg^R$ spk1-GFP-2xFLAG::$Kan^R$*

**C**

**KT4300** *$h^{90}$ ade6.M216 leu1.32 byr1::$ClonNAT^R$ spk1-GFP-2xFLAG::$Kan^R$*

**KT5215** *$h^{90}$ ade6.M210 leu1.32 ras1.G17V::$LEU2^+$ byr1::$ClonNAT^R$ spk1-GFP-2xFLAG::$Kan^R$*

**Figure 3**

**A**

**KT3082** *$h^{90}$ ade6.M216 leu1.32 spk1-GFP-2xFLAG::$Kan^R$*

**KT4061** *$h^{90}$ ade6.M216 leu1.32 scd1::$ClonNAT^R$ spk1-GFP-2xFLAG::$Kan^R$*

**KT4056** *$h^{90}$ ade6.M210 leu1.32 ras1.G17V::$LEU2^+$ scd1::$ClonNAT^R$ spk1-GFP-2xFLAG::$Kan^R$*

**B**

**KT3082** *$h^{90}$ ade6.M216 leu1.32 spk1-GFP-2xFLAG::$Kan^R$*

**KT4061** *$h^{90}$ ade6.M216 leu1.32 scd1::$ClonNAT^R$ spk1-GFP-2xFLAG::$Kan^R$*

**C**

**KT4047** *$h^{90}$ ade6.M216 leu1.32 byr1.DD::$Hyg^R$ scd1::$ClonNAT^R$ spk1-GFP-2xFLAG::$Kan^R$*

**D**

**KT4323** *$h^{90}$ ade6.M216 leu1.32 ras1::$ClonNAT^R$ spk1-GFP-2xFLAG::$Kan^R$*

**KT3435** *$h^{90}$ ade6-M216 leu1.32 byr1.DD::$Hyg^R$ spk1-GFP-2xFLAG::$Kan^R$*

**KT4359** *$h^{90}$ ade6.M216 leu1.32 byr1.DD::$Hyg^R$ ras1::$ClonNAT^R$ spk1-GFP-2xFLAG::$Kan^R$*

**E**

**KT3763** *$h^{90}$ ade6.M216 leu1.32 byr2::$ClonNAT^R$ spk1-GFP-2xFLAG::$Kan^R$*

**KT3435** *$h^{90}$ ade6-M216 leu1.32 byr1.DD::$Hyg^R$ spk1-GFP-2xFLAG::$Kan^R$*

**KT4010** *$h^{90}$ ade6-M216 leu1.32 byr1.DD::$Hyg^R$ byr2::$ClonNAT^R$ spk1-GFP-2xFLAG::$Kan^R$*

**F**

**KT4323** h$^{90}$ ade6.M216 leu1.32 ras1::ClonNAT$^{R}$ spk1-GFP-2xFLAG::Kan$^{R}$

**KT4359** h$^{90}$ ade6.M216 leu1.32 byr1.DD::Hyg$^{R}$ ras1::ClonNAT$^{R}$ spk1-GFP-2xFLAG::Kan$^{R}$

**KT3763** h$^{90}$ ade6.M216 leu1.32 byr2::ClonNAT$^{R}$ spk1-GFP-2xFLAG::Kan$^{R}$

**KT4010** h$^{90}$ ade6-M216 leu1.32 byr1.DD::Hyg$^{R}$ byr2::ClonNAT$^{R}$ spk1-GFP-2xFLAG::Kan$^{R}$

**Fig. 4**

**A, B**

**KT5077** h$^{90}$ ade6.M210 leu1.32 ura4.294::[Pshk1:ScGIC2 CRIB:GFP3:ura4$^{+}$]

**KT5082** h$^{90}$ ade6.M216 leu1.32 ras1.G17V::LEU2$^{+}$ ura4.294::[Pshk1:ScGIC2 CRIB:GFP3:ura4$^{+}$]

**Fig. 5**

**A**

**KT5940** h$^{90}$ ade6.M216 leu1.32 ras1.G17V::Hyg$^{R}$ spk1-GFP-2xFLAG::Kan$^{R}$

**B**

**KT5938** h$^{90}$ ade6.M216 leu1.32 ras1.G17V::Hyg$^{R}$ ura4.294::[Pshk1:ScGIC2 CRIB:GFP3:ura4$^{+}$]

**Fig. 6**

**A, B**

**KT4376** h$^{90}$ ade6.M216 leu1.32 ste4::ClonNAT$^{R}$ spk1-GFP-2xFLAG::Kan$^{R}$

**KT5143** h$^{90}$ ade6.M210 leu1.32 ste4::ClonNAT$^{R}$ ras1.G17V::LEU2$^{+}$ spk1-GFP-2xFLAG::Kan$^{R}$

**KT5136** h$^{90}$ ade6-M216 leu1.32 ste4::ClonNAT$^{R}$ byr1.DD::Hyg$^{R}$ spk1-GFP-2xFLAG::Kan$^{R}$

**C, D**

**KT4333** h$^{90}$ ade6.M216 leu1.32 ste6::ClonNAT$^{R}$ spk1-GFP-2xFLAG::Kan$^{R}$

**KT4998** h$^{90}$ ade6.M210 leu1.32 ste6::Hyg$^{R}$ ras1.G17V::LEU2$^{+}$ spk1-GFP-2xFLAG::Kan$^{R}$

**KT5139** h$^{90}$ ade6-M216 leu1.32 ste6::ClonNAT$^{R}$ byr1.DD::Hyg$^{R}$ spk1-GFP-2xFLAG::Kan$^{R}$

**Fig. 7**

**A, B**

**KT4335** h$^{90}$ ade6.M216 leu1.32 gpa1::ClonNAT$^{R}$ spk1-GFP-2xFLAG::Kan$^{R}$

**KT5023** h$^{90}$ ade6.M210 leu1.32 gpa1::ClonNAT$^{R}$ ras1.G17V::LEU2$^{+}$ spk1-GFP-2xFLAG::Kan$^{R}$

**KT4353** h$^{90}$ ade6-M216 leu1.32 gpa1::ClonNAT$^{R}$ byr1.DD::Hyg$^{R}$ spk1-GFP-2xFLAG::Kan$^{R}$

**KT5035** h$^{90}$ ade6.M210 leu1.32 gpa1::ClonNAT$^{R}$ ras1.G17V::LEU2$^{+}$ byr1.DD::Hyg$^{R}$ spk1-GFP-2xFLAG::Kan$^{R}$

**C, D**

**KT4190** h$^{-}$ ade6.M216 leu1.32 spk1-GFP-2xFLAG::Kan$^{R}$

**KT5059** h$^{-}$ ade6.M216 leu1.32 ura4.d18 gpa1.QL::ura4$^{+}$ spk1-GFP-2xFLAG::Kan$^{R}$

**KT4233** *h*$^{-}$ *ade6.M216 leu1.32 ras1.G17V::LEU2*$^{+}$ *spk1-GFP-2xFLAG::Kan*$^{R}$

**KT5070** *h*$^{-}$ *ade6.M216 leu1.32 ura4.d18 ras1::ClonNAT*$^{R}$ *gpa1.QL::ura4*$^{+}$ *spk1-GFP-2xFLAG::Kan*$^{R}$

**KT4194** *h*$^{-}$ *ade6.M216 leu1.32 byr1.DD::Hyg*$^{R}$ *spk1-GFP-2xFLAG::Kan*$^{R}$

**Supplementary Figure S1**

**D, E and F**

**KT301** *h*$^{90}$ *ade6-M216 leu1.32*

**KT3082** *h*$^{90}$ *ade6.M216 leu1.32 spk1-GFP-2xFLAG::Kan*$^{R}$

**G**

**KT3082** *h*$^{90}$ *ade6.M216 leu1.32 spk1-GFP-2xFLAG::Kan*$^{R}$

**KT4300** *h*$^{90}$ *ade6.M216 leu1.32 byr1::ClonNAT*$^{R}$ *spk1-GFP-2xFLAG::Kan*$^{R}$

**Supplementary Figure S2 and S3**

**KT3082** *h*$^{90}$ *ade6.M216 leu1.32 spk1-GFP-2xFLAG::Kan*$^{R}$

**KT3435** *h*$^{90}$ *ade6-M216 leu1.32 byr1.DD::Hyg*$^{R}$ *spk1-GFP-2xFLAG::Kan*$^{R}$

**KT3084** *h*$^{90}$ *ade6.M210 leu1.32 ras1.G17V::LEU2*$^{+}$ *spk1-GFP-2xFLAG::Kan*$^{R}$

**KT3439** *h*$^{90}$ *ade6.M210 leu1.32 ras1.G17V::LEU2*$^{+}$ *byr1.DD::Hyg*$^{R}$ *spk1-GFP-2xFLAG::Kan*$^{R}$

**KT4061** *h*$^{90}$ *ade6.M216 leu1.32 scd1::ClonNAT*$^{R}$ *spk1-GFP-2xFLAG::Kan*$^{R}$

**Supplementary Figure S4**

**A and B**

**KT5951** *h*$^{90}$ *ade6.M216 leu1 spk1-GFP-2xFLAG::Kan*$^{R}$ *smd2-tdTomato::Hyg*$^{R}$

**Supplementary Figure S5**

**A and B**

**KT3435** *h*$^{90}$ *ade6-M216 leu1.32 byr1.DD::Hyg*$^{R}$ *spk1-GFP-2xFLAG::Kan*$^{R}$

**Supplementary Figure S6**

**A and D**

**KT3435** *h*$^{90}$ *ade6-M216 leu1.32 byr1.DD::Hyg*$^{R}$ *spk1-GFP-2xFLAG::Kan*$^{R}$

**B and D**

**KT3084** *h*$^{90}$ *ade6.M210 leu1.32 ras1.G17V::LEU2*$^{+}$ *spk1-GFP-2xFLAG::Kan*$^{R}$

**Supplementary Figure S7**

**KT5107** *h*$^{90}$ *ade6.M210 leu1.32 ras1::ClonNAT*$^{R}$ *ura4.294::[Pshk1:ScGIC2 CRIB:GFP3:ura4*$^{+}$*]*

**KT5077** *h*$^{90}$ *ade6.M210 leu1.32 ura4.294::[Pshk1:ScGIC2 CRIB:GFP3:ura4*$^{+}$*]*

**KT5082** *$h^{90}$ ade6.M216 leu1.32 ras1.G17V::LEU2$^{+}$ ura4.294::[Pshk1:ScGIC2 CRIB:GFP3:ura4$^{+}$]*

**KT5551** *$h^{90}$ ade6.M210 leu1.32 rga4::Hyg$^{R}$ ura4.294::[Pshk1:ScGIC2 CRIB:GFP3:ura4$^{+}$]*

**KT5554** *$h^{90}$ ade6.M216 leu1.32 ras1.G17V::LEU2$^{+}$ rga4::Hyg$^{R}$ ura4.294::[Pshk1:ScGIC2 CRIB:GFP3:ura4$^{+}$]*

**Supplementary Figure S8**

**A**

**KT4323** *$h^{90}$ ade6.M216 leu1.32 ras1::ClonNAT$^{R}$ spk1-GFP-2xFLAG::Kan$^{R}$*

**KT3435** *$h^{90}$ ade6-M216 leu1.32 byr1.DD::Hyg$^{R}$ spk1-GFP-2xFLAG::Kan$^{R}$*

**KT4359** *$h^{90}$ ade6.M216 leu1.32 byr1.DD::Hyg$^{R}$ ras1::ClonNAT$^{R}$ spk1-GFP-2xFLAG::Kan$^{R}$*

**B**

**KT3763** *$h^{90}$ ade6.M216 leu1.32 byr2::ClonNAT$^{R}$ spk1-GFP-2xFLAG::Kan$^{R}$*

**KT3435** *$h^{90}$ ade6-M216 leu1.32 byr1.DD::Hyg$^{R}$ spk1-GFP-2xFLAG::Kan$^{R}$*

**KT4010** *$h^{90}$ ade6-M216 leu1.32 byr1.DD::Hyg$^{R}$ byr2::ClonNAT$^{R}$ spk1-GFP-2xFLAG::Kan$^{R}$*

**C**

**KT4376** *$h^{90}$ ade6.M216 leu1.32 ste4::ClonNAT$^{R}$ spk1-GFP-2xFLAG::Kan$^{R}$*

**KT5143** *$h^{90}$ ade6.M210 leu1.32 ste4::ClonNAT$^{R}$ ras1.G17V::LEU2$^{+}$ spk1-GFP-2xFLAG::Kan$^{R}$*

**KT5136** *$h^{90}$ ade6-M216 leu1.32 ste4::ClonNAT$^{R}$ byr1.DD::Hyg$^{R}$ spk1-GFP-2xFLAG::Kan$^{R}$*

**D**

**KT4333** *$h^{90}$ ade6.M216 leu1.32 ste6::ClonNAT$^{R}$ spk1-GFP-2xFLAG::Kan$^{R}$*

**KT4998** *$h^{90}$ ade6.M210 leu1.32 ste6::Hyg$^{R}$ ras1.G17V::LEU2$^{+}$ spk1-GFP-2xFLAG::Kan$^{R}$*

**KT5139** *$h^{90}$ ade6-M216 leu1.32 ste6::ClonNAT$^{R}$ byr1.DD::Hyg$^{R}$ spk1-GFP-2xFLAG::Kan$^{R}$*

**Supplementary Figure S9**

**KT5940** *$h^{90}$ ade6.M216 leu1.32 ras1.G17V::Hyg$^{R}$ spk1-GFP-2xFLAG::Kan$^{R}$*

**Supplementary Figure S10**

**A**

**KT4335** *$h^{90}$ ade6.M216 leu1.32 gpa1::ClonNAT$^{R}$ spk1-GFP-2xFLAG::Kan$^{R}$*

**KT5023** *$h^{90}$ ade6.M210 leu1.32 gpa1::ClonNAT$^{R}$ ras1.G17V::LEU2$^{+}$ spk1-GFP-2xFLAG::Kan$^{R}$*

**KT4353** *$h^{90}$ ade6-M216 leu1.32 gpa1::ClonNAT$^{R}$ byr1.DD::Hyg$^{R}$ spk1-GFP-2xFLAG::Kan$^{R}$*

**KT5035** *$h^{90}$ ade6.M210 leu1.32 gpa1::ClonNAT$^{R}$ ras1.G17V::LEU2$^{+}$ byr1.DD::Hyg$^{R}$ spk1-GFP-2xFLAG::Kan$^{R}$*

**B**

**KT4190** *$h^{-}$ ade6.M216 leu1.32 spk1-GFP-2xFLAG::Kan$^{R}$*

**KT5059** *$h^{-}$ ade6.M216 leu1.32 ura4.d18 gpa1.QL::ura4$^{+}$ spk1-GFP-2xFLAG::Kan$^{R}$*

**KT4233** *h$^-$ ade6.M216 leu1.32 ras1.G17V::LEU2$^+$ spk1-GFP-2xFLAG::Kan$^R$*

**KT5070** *h$^-$ ade6.M216 leu1.32 ura4.d18 ras1::ClonNAT$^R$ gpa1.QL::ura4$^+$ spk1-GFP-2xFLAG::Kan$^R$*

**KT4194** *h$^-$ ade6.M216 leu1.32 byr1.DD::Hyg$^R$ spk1-GFP-2xFLAG::Kan$^R$*
